# Dissecting functional regulatory convergence over 160 million years of therian evolution

**DOI:** 10.1101/2025.07.10.662670

**Authors:** Navya Shukla, Laura E. Cook, Davide M. Vespasiani, Andrew J. Pask, Irene Gallego Romero

## Abstract

Understanding the molecular mechanisms underpinning convergent traits can provide insight into the predictability of the evolutionary process. The thylacine, an extinct marsupial carnivore, has been long known to have a very similar craniofacial morphology to that of eutherian canids, despite having diverged ***≈***160 million years ago. Using a massively parallel reporter assay (MPRA), we tested the regulatory potential of a set of previously identified highly conserved craniofacial enhancers that showed convergent sequence acceleration in thylacine and wolf (TWARs). We compared orthologous sequences from six different taxa, including outgroup taxa with non-convergent craniofacial phenotypes (Tasmanian devil and giant panda) and ancestral reconstructions for marsupial and placental carnivores (Dasyuromorphia and Carnivora). Dense 10bp tiling allowed us to thoroughly examine features associated with activity, including percentage GC content and transcription factor binding motifs turnover. We identified marked conservation of levels and patterns of regulatory activity across all six taxa, as well as multiple cases of differentially active TWARs within each clade, with both thylacine and wolf driving functional divergence. However, evidence of convergent divergence was limited to a set of neighbouring TWARs near the neural crest gene *Phox2b* – one of which exhibited reduced activity in the wolf, and the other in the thylacine. Ultimately our findings suggest that shared gene regulatory potential is not a feature of these convergently accelerated regions in the thylacine and wolf.

## Introduction

Convergent evolution is the independent emergence of the same traits in multiple lineages or populations, possibly in response to shared selective pressures. Echolocation, for example, is thought to have independently originated in marine mammals and bats as an adaptation to their low visibility habitats [1, 2], with parallel amino-acid substitutions in the cochlear motor protein *Prestin* in these animals pointing to its key specialised role in enabling the phenotype[3, 4]. However, other traits, such as the development of winged structures in birds and bats, exhibit convergence at the phenotypic level, but not necessarily at the level of independent base substitutions, highlighting the flexibility of developmental toolkits and ontogenetic processes [5]. Identifying and studying the genetic basis of convergent traits can therefore provide unique insight into the role of adaptation and constraint in trait evolution, and the repeatability of evolutionary processes.

The extinct marsupial thylacine (*Thylacinus cynocephalus*, also known as the Tasmanian tiger, and the Tasmanian dog by early European colonisers), displays high levels of morphological convergence with placental canids [6], especially in the craniofacial region. In particular, morphometric analyses have shown high feature overlap in craniodental structures [7–9]. Although its specific predatory behaviours and diet are disputed, the thylacine is universally agreed to have been a hypercarnivore, like most placental canids [8, 10–12]. This phenotypic convergence is thus hypothesized to be an adaptation to a shared feeding ecology requiring shearing of prey flesh.

We recently identified a set of 339 candidate cis regulatory elements (CREs) active during murine craniofacial development that showed increased rates of substitution in the thylacine and the grey wolf (*Canis lupus*) relative to 61 vertebrate taxa. These thylacine-and-wolf-accelerated-regions, or TWARs [13], are promising candidates for explaining craniofacial convergence across these taxa, but their effects on craniofacial development cannot be directly tested in either species. Instead, here we have employed a massively parallel reporter assay (MPRA), a high-throughput approach to examine the gene regulatory potential of genomic regions [14, 15], to test for functional convergence in the TWARs and identify putative functional CREs across thylacine, wolf, and outgroup taxa. Using this approach we identified evidence of functional conservation across species in spite of significant sequence divergence, as well as examples of divergent regulatory activity associated with sequence acceleration in both the thylacine and wolf. Convergence in regulatory activity of orthologous sequences was rare, however, suggesting that the flexibility of the regulatory architecture in craniofacial development facilitates the emergence of diverse paths to same trait.

## Results

### Designing an MPRA to compare marsupial and placental enhancer sequences

To investigate the genetic basis of convergent evolution in the thylacine and wolf, we performed an episomal MPRA testing the regulatory potential of 295 orthologous TWAR sequences from the two species, alongside orthologous sequences from two species with a non-convergent morphology and similar divergence times from the focal species, the Tasmanian devil, *Sarcophilius harrisi* (thylacine-Tasmanian devil TMRCA: 25-35 MYA [16]) and the giant panda, *Ailuropoda melanoleuca* (panda-wolf TMRCA = 40-50 MYA, [17]) and ancestral reconstructions of the two parent clades (Dasyuromorphia and Carnivora) (Figure 1A). We also included the murine orthologous sequence of all tested TWARs to account for any effects driven by the mouse *trans*-regulatory environment, as given a lack of suitable marsupial cell lines we performed the assay in mouse preosteoblasts and cranial neural crest cells (CNCCs), both involved in craniofacial development [18–20].

**Figure 1.**
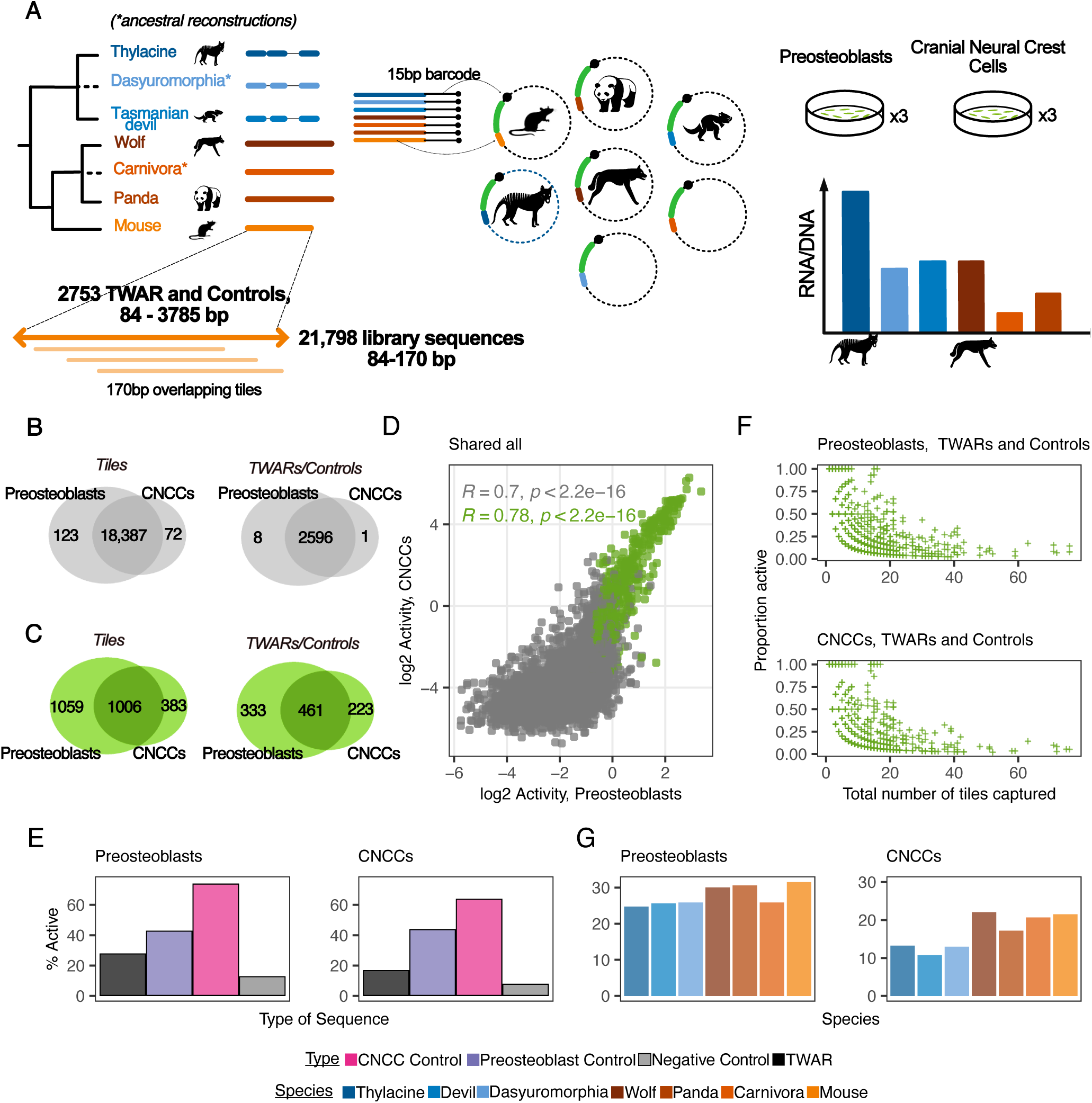
Performing an MPRA to test convergence of of regulatory activity. **A.** Orthologous TWAR sequences from seven different taxa — two test (thylacine and wolf), two outgroup (Tasmanian devil and panda), two ancestral (Dasyuromorphia and Carnivora) and one control (mouse) taxa were tested for activity in an episomal MPRA. Elements longer than 170 bp were covered with overlapping tiles in 10 bp steps. The cloned library was transfected into preosteoblasts and CNCCs. TWARs were then tested for convergent differential activity within each clade. **B.** Number of tiled library sequences and un-tiled TWARs and controls captured in each cell line. **C.** Number of active tiled library sequences and active un-tiled TWARs and controls captured in each cell line. **D.** Correlation of *log*_2_ activity for library sequences in preosteoblasts and CNCCs (with significantly active library sequences in green). **E.** Proportion of TWARs, cell-line specific positive controls and scrambled negative controls determined to be active in each cell line. **F.** Proportion of active tiles versus the total number of tiles captured for each multi-tiled TWAR and control sequence, for each cell line. **G.** Proportion of TWARs from each taxa determined to be active in each cell line.

TWAR lengths ranged from 84 to 1,001 bp; thus we tiled across elements longer than 170 bp with a 10 bp sliding window, enabling us to examine the regulatory potential of the whole TWAR at high resolution. However, because TWAR orthologs can display variable lengths and alignment, we note that tiles did not always map 1-to-1 across taxa. Our library consisted of 21,798 unique oligos (representing 2,757 distinct TWAR and control elements) (**Supplementary Table 1**). The synthesized library was tagged with a median of 337 barcodes/oligo, assembled into an episomal vector, and transfected into 3 x replicates per cell line. After sequencing and quality control, we found that *≈* 85% of the library was captured in both cell lines (preosteoblasts: 18,510, CNCCs: 18,459) (Figure 1B), with 2,065 significantly active sequences (11.2% of tested) in preosteoblasts (*log*_2_ activity between –0.82 and 3.36) and 1,389 (7.5%) in CNCCs (*log*_2_ activity between –3.5 and 6.28) (Figure 1C). The lower range in activity values in the CNCCs is due to a technical artefact in the sequencing of mRNA from CNCCs (Supplementary Figure 1 and 2, see Methods); despite this, 72.4% (1006) of the library sequences active in CNCCs were also active in preosteoblasts, demonstrating high reproducibility. Spearman’s correlation between activity values in the two cell lines was 0.70, increasing to 0.78 for shared active sequences (Figure 1D).

In addition, both the proportion of the negative controls that were active, as well as the activity values for these controls were lower than those for the positive controls and the TWARs (Figure 1E Supplementary Figure 3A and C, pairwise t-tests results in **Supplementary Table 2**).

Our overlapping tile design helped compensate for the loss of individual library sequences during cloning/transfection. By examining sequence coverage across complete TWAR and control sequences, we found that 2569 (98.7%) and 2563 (98.7%) have at least 80% coverage in preosteoblasts and CNCCs datasets, respectively (Supplementary Figure 4). We observed that regulatory potential is not uniform across tested sequences — for both TWARs and positive controls the overall region length was inversely correlated with the proportion of active tiles (Figure 1F); for instance, the mean and median number of active tiles for multi-tiled TWARs was 2.0 and 2.6 in the preosteoblasts and 1.0 and 2.5 in the CNCCs. When focusing on the subset of active TWARs in each cell line for which i) all tiles were captured and ii) more than one tile was found active (146 in preosteoblasts and 78 in CNCCs), the median distance between active tiles was estimated to be 10 nt, i.e. most active tiles within this subset were neighbouring each other and shared most of their sequence (Supplementary Figure 5). These observations are consistent with a role for small transcription factor binding motifs in driving regulatory activity within promoter and enhancer regions.

Finally, to assess the impact of the murine *trans*-regulatory environment, we compared TWAR activity for each of the test taxa with that of the orthologous mouse sequence. Activity values were not significantly different between the mouse and any of the other taxa in either cell line (pairwise t-test results in **Supplementary Table 3**, Supplementary Figure 3B and D). The percentage of active TWARs did not differ significantly among the placental taxa, with pairwise differences ranging from 0.9-5.6% in preosteoblasts and 0.5-4.3% in CNCCs (Figure 1G, two-proportion tests in **Supplementary Table 4**). All placental taxa, excluding the Carnivora in preosteoblasts, had a greater percentage of active TWARs than the three marsupial taxa (pairwise differences ranging 4.1-6.8% in preosteoblasts and 4.0-11.3 % in CNCCs). This could indicate a slight placental bias arising from the placental *trans* environment; however, these differences were not statistically significant (two-proportion tests in **Supplementary Table 4**).

### Genomic features underlying TWAR activity

To identify genomic features driving activity in our assay, we first used FIMO to scan TWARs for transcription factor binding motifs (TFBMs) from the HOCOMOCO Mouse Core v11, summarising all motifs inside a TWAR into a single metric that ranges from 0-1 and can be directly compared across all orthologs of a given TWAR (see Methods). Active TWARs have significantly higher aggregated FIMO scores than inactive TWARs (Figure 2A) (pairwise t-test, preosteoblasts *p*-value: 3.386 *×* 10*^−^*^17^, CNCCs *p*-value: 4.794 *×* 10*^−^*^17^). We also observed that active TWARs also had significantly higher percentage GC content, consistent with the well-known association of GC-rich DNA with open chromatin, promoters and broadly active enhancers [21, 22] (Figure 2B) (t-test preosteoblasts *p*-value: 3.027 *×* 10*^−^*^10^, CNCCs *p*-value: 1.459 *×* 10*^−^*^18^). Finally, we classified TWARs within 1 kb of a mouse (GRCm38) TSS as promoters and the rest as enhancers (26 and 271, respectively), to test for any differences in activity between distal and proximal elements. There was no enrichment for activity in proximal sequences relative to distal ones (Fisher’s exact test, preosteoblasts *p*-value: 0.447, CNCCs *p*-value: 0.454) (Figure 2C). Furthermore, there was only a weak association between the absolute distance from a transcription start site in mouse to the proportion of taxa in which a TWAR was determined to be active, in either cell line (see Methods).

**Figure 2.**
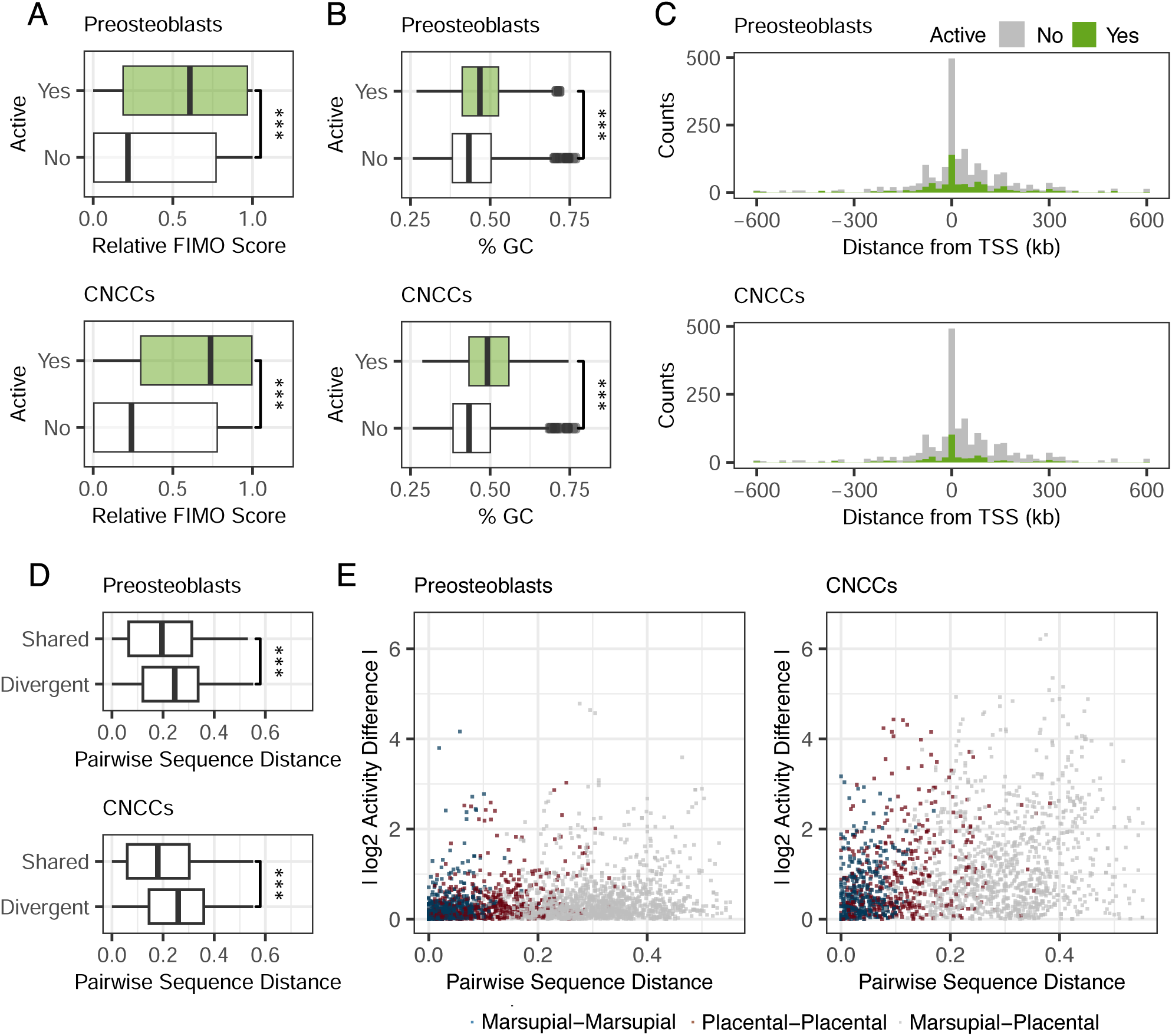
Examining sequence features driving activity of TWARs. Distribution of **A.** aggregated FIMO scores and **B.** Percentage GC content for active and non-active TWARs, for both cell lines. **C.** Distribution of active and non-active TWARs in cell lines by their distance from a transcription start site in the mouse genome. **D.** Distribution of pairwise sequence distances between each pair of TWAR orthologs that display shared or divergent activity. **E.** Correlation between pairwise sequence distance and absolute activity differences for each pair of TWAR orthologs.

We then examined the role of sequence divergence in influencing patterns of TWAR activity. Overall, 167 TWARs in preosteoblasts and 147 in the CNCCs were active in at least one of the six tested taxa. Of these 117 (70.0%) and 83 (56.5%), respectively, were active in more than 1 taxon (Supplementary Figure 6). Considering only TWARs with demonstrated activity in at least one taxa, we observed significantly higher pairwise sequence distances between pairs of orthologous TWARs that show divergent activity (one active, one inactive) than pairs of orthologous TWARs for which activity was shared (both active or inactive) (Figure 2D) (pairwise t-test preosteoblasts *p*-value: 1.950 *×* 10*^−^*^10^, CNCCs *p*-value: 1.440 *×* 10*^−^*^19^). Differences in relative FIMO scores were also significantly larger between pairs of orthologous TWARs with divergent activity (Supplementary Figure 7) (t-test preosteoblasts *p*-value: 0.0004, CNCCs *p-value*: 4.630 *×* 10*^−^*^15^). Spearman’s *ρ* between pairwise sequence distance and absolute activity differences was 0.21 in preosteoblasts and 0.28 in CNCCs (Figure 2E). We observed higher correlations for the placental-placental comparisons (*ρ* 0.30 in preosteoblasts, 0.28 in CNCCs) than the marsupial-marsupial comparisons (*ρ* 0.22 in preosteoblasts, 0.15 in CNCCs). This is likely due to the overall greater sequence similarity between the marsupial orthologs (mean pairwise sequence distance thylacine vs Tasmanian devil: 0.068, thylacine vs Dasyuromorphia: 0.015, Tasmanian devil vs Dasyuromorphia: 0.054, wolf vs panda: 0.166, wolf vs. Carnivora: 0.154 and panda vs Carnivora: 0.043) (Supplementary Figure 8).

### Examining patterns of TWAR activity across taxa

Seventeen TWARs displayed significant activity across all six taxa in the preosteoblasts, with 6 being also universally active in CNCCs (Figure 3A, **Supplementary Table 5**); variation across taxa in TWAR lengths and thus tile numbers confounded 1:1 tile activity comparisons but activity-by-tile plots of all conserved elements are available as **Supplementary File 1**. The top-ranked TWAR by activity in all taxa was TWAR17.chr6; 85% of tiles in this TWAR were significantly active in all taxa. Our sliding window design identified a consistent pattern of decreasing activity when moving across tiles (5’ to 3’) in the TWAR (Figure 3B). Although only the first six panda and seven of the first eight Carnivora tiles were present in the final library, the orthologous mouse sequence had activity patterns nearly identical to those of the wolf, with a peak of activity in tile 12 (Wolf—Mouse Spearman’s *ρ* preosteoblasts: 0.86, CNCCs: 0.91) (Supplementary Figure 9). Zooming in, our FIMO analyses identified a cluster of TFBMs within the first 70 bp of TWAR17.chr6 (Supplementary Figure 10), while the peak in activity in tile 12 for the wolf ortholog (spanning positions 111-280 in the sequence) overlapped a placental-specific binding site for LYL1 (at positions 254-267). TWAR17.chr6 intersects with a super-enhancer in human embryonic stem cells for the neighbouring *Atf7ip* [23], known to inhibit premature differentiation of progenitor cells (including the preosteoblasts used in our study [24–26]). Thus it is likely that the endogenous gene regulatory network that regulates *Atf7ip* expression is highly active in our cell lines and driving these activity patterns.

**Figure 3.**
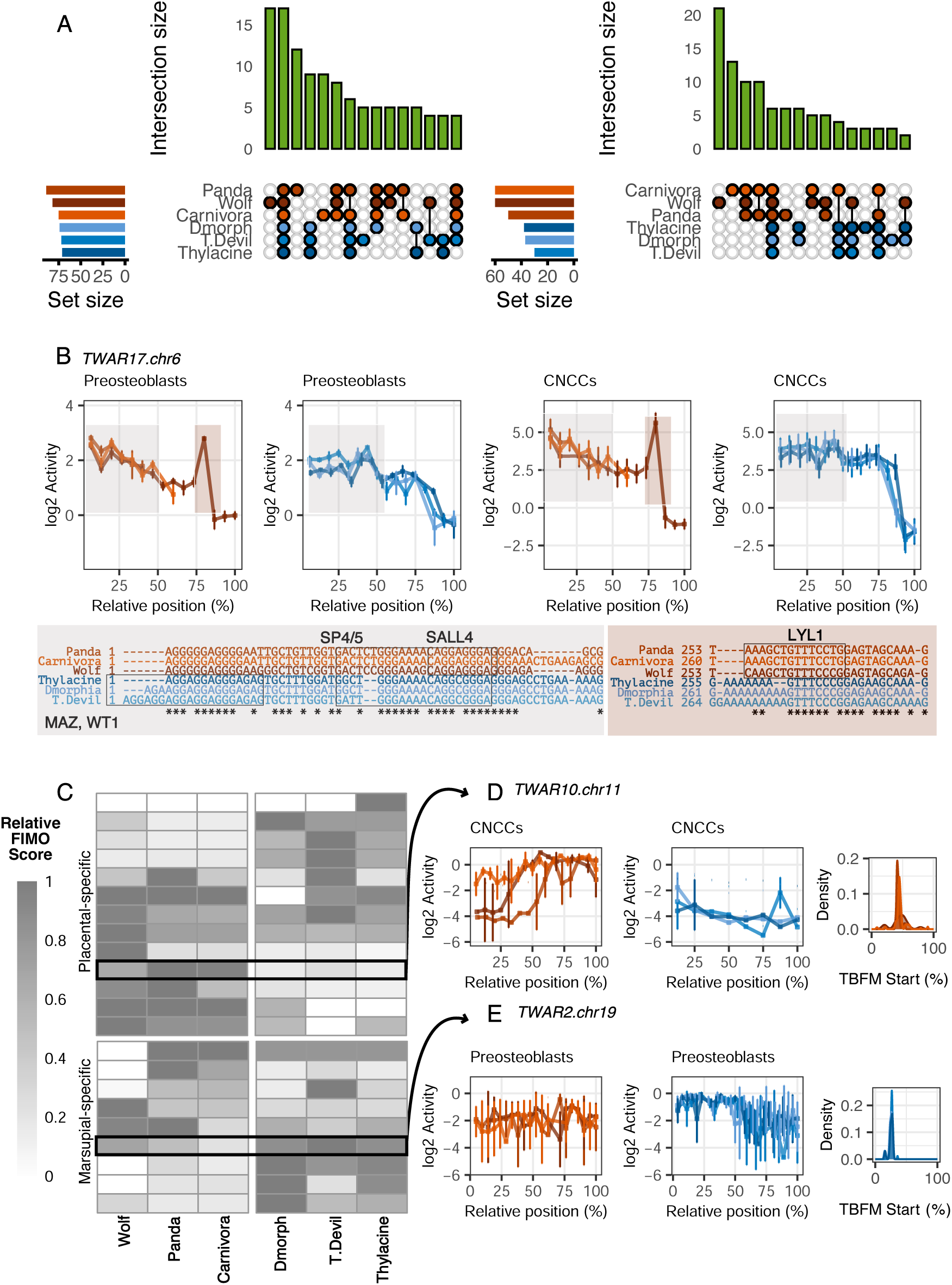
Functional conservation of activity across taxa. **A.** Top 15 intersections of active TWARs across taxa for each cell line. **B.** Patterns of activity across tiles for TWAR17.chr6 in marsupial and placental taxa. Due to variation in the number of tiles across taxa, tile number has been summarized as relative to the total number of tiles (“Relative position”). Boxes identify TFBMs motifs present in the sequences for this TWAR highlighted in the plots. **C.** Heatmap of relative FIMO scores for every TWAR with marsupial- or placental-specific activity in either cell line. **D.** Activity across tiles of the placental-active TWAR10.chr11 for all taxa in the CNCCs, along with distribution of binding motifs across sequence length for the placentals. **E.** Activity across tiles of the marsupial-active TWAR2.chr19 for all taxa in the preosteoblasts, along with distribution of binding motifs across sequence length specifically for the marsupials.

As originally defined, all TWARs overlapped H3K9ac and H3K27ac peaks in mouse craniofacial tissues collected at E11.5 in [27]). Recently, we published a novel H3K27ac and H3K4me3 ChIP-seq dataset from craniofacial tissue of newborn fat-tailed dunnarts (*Sminthopis crassicaudata*), a carnivorous dasyurid marsupial model species [28] with a TMRCA to the thylacine of 25-35 million years [16]. Of the 17 universally active TWARs, 12, including TWAR17.chr6 overlapped or were adjacent to dunnart H3K27ac and/or H3K4me3 peaks, also confirming their activity in the craniofacial region of a developing dasyurid and their broad enhancer potential across therians.

Having established that our MPRA could distinguish TWARs with broad gene regulatory potential, we sought to establish its capacity to detect regulatory changes over long evolutionary distances. We identified TWARs with marsupial and placental-specific regulatory behaviour — 8 TWARs in preosteoblasts and 1 in CNCCs, were exclusively active in the marsupial carnivores, and 6 TWARs in preosteoblasts and 8 in CNCCs (one overlapping) were exclusively active in the placental carnivores (**Supplementary Table 5** and **Supplementary File 2**). Aggregated FIMO scores were higher in the marsupials for marsupial-active TWARs (mean difference marsupials—placentals 0.180, pairwise t-test *p*-value 0.055) and higher in placentals for the placental active TWARs (mean difference 0.128, *p*-value 0.128). While these differences were not significant overall, clustering elements by their aggregated FIMO scores highlights individual examples of clade-specific regulatory activity associated with clade-specific TFBMs (Figure 3C). For instance, TWAR10.chr11 displayed placental-specific activity in CNCCs (Figure 3D), with identical tile activity patterns across all three placental taxa. These correlated with the presence of TFBMs for Sp and KLF family proteins in these sequences. Similarly, the first half of TWAR2.chr19 displayed marsupial-specific activity (Figure 3E), consistent with a cluster of TFBMs for ELF5, ETV2, KLF15, MAZ, Sp5 and Sp1 in the region. TWAR2.chr19 was also distinguished by a stronger H3K27ac and H3K4me3 signal in the dunnart (which shares these motifs) than the mouse (Supplementary Figure 11). Across the clade-specific active TWARs, 5 (63%) marsupial-specific and 4 (31%) placental-specific TWARs overlapped or were adjacent to H3K27ac and/or H3K4me3 peaks in dunnart craniofacial tissue, confirming the ability of our MPRA to detect less conserved activity patterns.

### Characterising differential TWAR activity in the thylacine and wolf

We then asked whether any TWARs exhibited activity patterns unique to the wolf or thylacine sequences. By assessing binary active/inactive trends, we found a distinct signal of unique behaviour for the wolf — 23 TWARs in preosteoblasts and 24 in CNCCs were either only active or inactive in this species (Supplementary Figure 6). Due to the generally lower levels of pairwise sequence divergence between marsupials than placentals (Supplementary Figure 8) in our TWAR, we hypothesised that differences in this clade may be subtler, which was borne out by our data: the thylacine was associated with only 7 uniquely active or inactive TWARs in preosteoblasts and 4 in CNCCs.

Moving beyond binary patterns, we tested for differential TWAR activity within each clade by adapting an ANOVA-based approach from [29] (see Methods). Given the large evolutionary and sequence distance separating our two focal species, we decided to carry out these analyses separately within each clade, thus asking whether wolf TWAR activity differed from that of the panda and Carnivora and, in parallel, whether thylacine TWAR activity differed from that of the Tasmanian devil and Dasyuromorphia. Furthermore, due to the lack of direct 1-to-1 orthology between tiles, we performed all tests by examining the most active tile in a TWAR for each species. Restricting our analyses to the subset of TWARs that were active in at least one taxon, we found 7 TWARs (out of 110) to be differentially active (DA) in preosteoblasts and 11 (out of 50) were DA in CNCCs for marsupials; 3 were shared across the two cell lines and 2 had the same direction of effect in both cell lines. Across the placental taxa, 22 (out of 134) TWARs were DA in preosteoblasts and 29 in CNCCs (out of 103); 12 were shared across the two cell lines including 11 with the same direction of effect.

We characterised effect sizes relative to our two focal species to disentangle the effect of thylacine and wolf acceleration. The range of effect sizes was smaller in the marsupial DA TWARs (preosteoblasts: –2.77 to 0.89, CNCCs: –3.17 to 2.93) than the placental ones (preosteoblasts: –2.21 to 2.59, CNCCs: –4.16 to 4.436), confirming that subtle activity differences were more frequent among the marsupials. Across DA TWARs, we observed linear and positive correlations for both marsupials and placentals in effect sizes in the focal species relative to the outgroup and the ancestor (Pearson’s R 0.72-0.91; Figure 4A). This agreement suggests that the thylacine and wolf elements were driving differentially activity within their clade, as expected given how the TWARs were initially identified. Supporting this, activity differences between the wolf and the panda, and the wolf and Carnivora were both significantly larger than activity differences between the panda and Carnivora (Figure 4B). While activity differences between the thylacine and Tasmanian devil were larger than activity differences between the Tasmanian devil and Dasyuromorphia in both cell lines, the comparison was only significant in CNCCs, due to the fewer number of observations in preosteoblasts. Lastly, differences in activity between the thylacine and Dasyuromorphia and between the Tasmanian devil and Dasyuromorphia were comparable in both cell lines. Finally, we asked whether TWARs with higher levels of sequence divergence between our focal and comparison taxa also exhibited differential activity patterns more often. Indeed, pairwise sequence distance between the wolf and the other placental TWARs was significantly greater in DA TWARs than in non-DA TWARs (Figure 4C). This was not the case in marsupials, again likely due to overall higher sequence similarity between marsupial orthologous sequences (Supplementary Figure 8). piou909879

**Figure 4.**
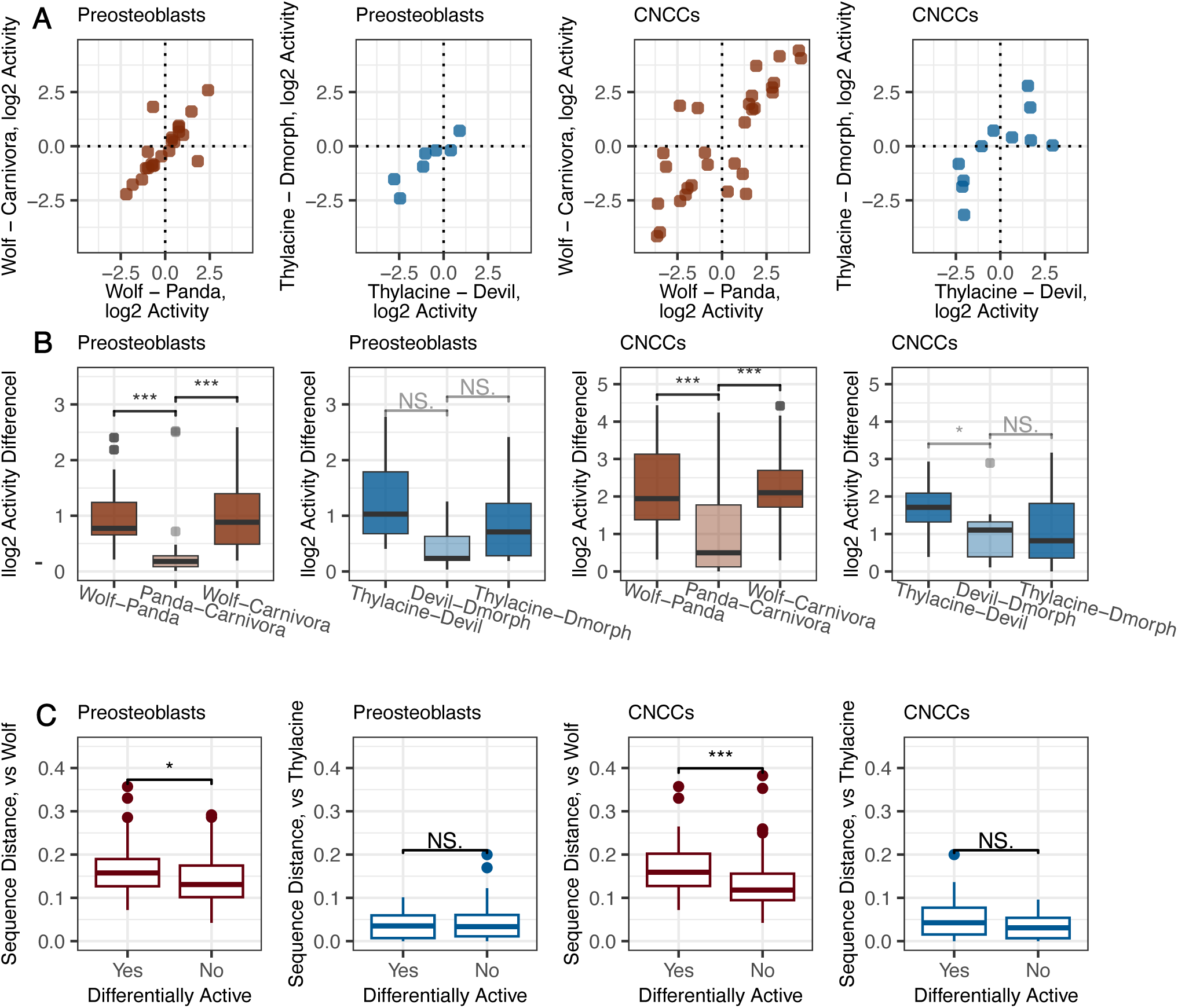
Differential activity of TWARs within the marsupials and placentals. **A.** Comparisons of activity differences between the test and outgroup taxa to the differences in activity between the test and ancestral taxa for marsupial and placental differentially active (DA) TWARs in both cell lines **B.** Distributions of activity differences between each pair of taxa for marsupial and placental DA TWARs in both cell lines. **C.** Distribution of pairwise sequence distances between the focal taxon and the outgroup and ancestral taxa for DA and non-DA TWARs in both cell lines.

### Tiling allows the identification of TFBMs associated with divergent regulatory activity

To identify associations between the presence of TFBMs and differential TWAR activity, we recalculated our FIMO scores for each TWAR relative to the most active ortholog separately for each clade. FIMO scores were overall comparable between DA TWARs than non-DA TWARs for both marsupials and placentals (Supplementary Figure 12). For placental DA TWARs, the direction of effect was significantly associated with differences in FIMO scores — TWARs with increased activity in the wolf relative to other placental taxa also have higher FIMO scores in the wolf than the other two taxa, and vice versa (Figure 5A). We proceeded to cluster DA TWARs in each clade by their FIMO scores, and identified several examples for which direction of activity could be linked to variation in FIMO scores (Figure 5B).

**Figure 5.**
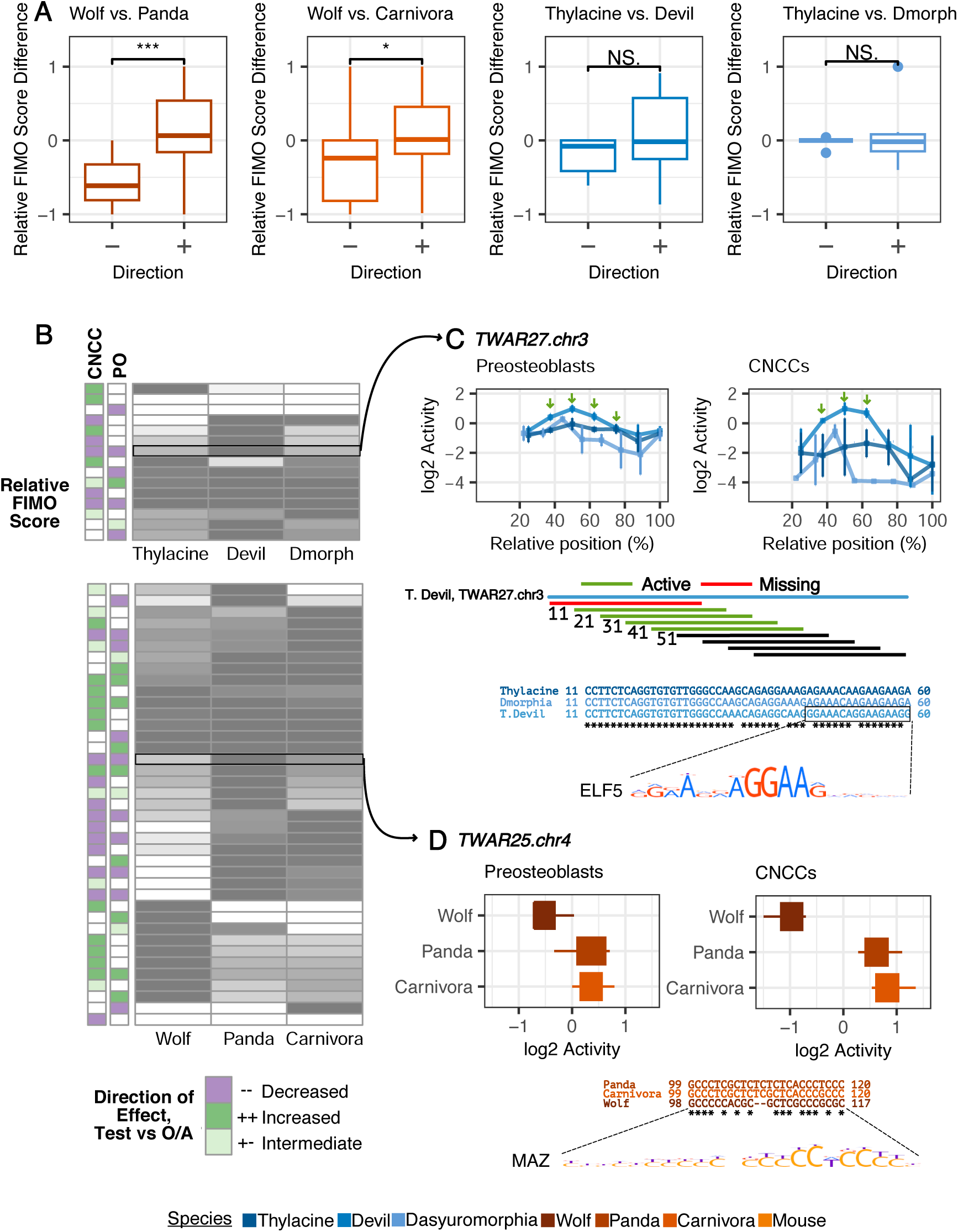
Transcription factor binding and differential activity. **A.** Differences in FIMO scores between the test species and the outgroup or ancestral taxon, for DA TWARs that have gained (+) or lost (-) activity in the focal species. **B.** Relative FIMO scores for differentially active (DA) TWARs in the marsupials and placentals for both cell lines. Direction of effect highlights if the thylacine/wolf (“test”) TWARs have gained activity relative to both outgroup (“O–)and ancestor (“A”) (++), lost active relative to both outgroup and ancestor (–) or have intermediate activity to that of the outgroup and ancestor (+-). Note the concordance in direction for DA TWARs across cell lines. **C.** Activity across tiles of the differentially active TWAR27.chr3 for marsupial taxa; highlighted are the Tasmanian devil-specific binding motifs associated with Tasmanian devil-active tiles for this TWAR. **D.** Activity values of the differentially activity TWAR25.chr4 for the placental taxa. The sequences alignments shows the wolf specific deletion and consequent loss of binding motifs, with one example provided.

We also took advantage of the fact that our tiling design traverses TWARs 10bp at a time to examine how activity patterns and binding motifs change across a TWAR’s length at high resolution. For instance, TWAR27.chr3 was highly differentially active between marsupials (Figure 5C; adjusted *p*-value <1 x 10*^−^*^6^ in all replicates, **Supplementary Table 6**-**Supplementary Table 9**). Both Tasmanian devil and Dasyuromorphia TWAR sequences were more active than the thylacine one, with higher differences in activity between Tasmanian devil and thylacine (mean *log*_2_ effect size preosteoblasts –1.029, CNCCs –2.342) than between the thylacine and Dasyuromorphia (mean *log*_2_ effect size preosteoblasts –0.340, CNCCs, –0.817). In the Tasmanian devil activity peaked between tiles 2-5; examining sequence covered uniquely by these tiles (11-60 nt) we found 3 fixed single nucleotides differences in the Tasmanian devil relative to the other marsupials, which span predicted TFBMs for ELF5, ETS2, ETV2, MAZ and SP5. We observed a similar trend in TWAR25.chr4, a single-tile TWAR with significantly decreased activity in the wolf relative to the other placentals in both cell lines (Figure 5D). Motifs for three different transcription factors (KLF5, MAZ, SP5) were found in both panda and Carnivora but not wolf, potentially attributed to 1-2 bp gaps in the wolf sequence that disrupt these motifs. These examples illustrate how combining evidence of activity and tiling patterns from the MPRA with sequence changes and FIMO results gave us with base-pair level insights into drivers of functional change within individual non-coding elements.

### Limited evidence of convergence in differential activity between marsupials and placentals

Our initial hypothesis was that shared sequence acceleration of the TWARs for thylacine and wolf would be reflected in convergent patterns of regulatory activity. However, only two TWARs were DA across both the marsupials and placentals, TWAR6.chr14 and TWAR5.chr7 (Figure 6A). In both cases, we observed opposite directions of effect for the thylacine and the wolf (Figure 6B&C, Supplementary Figure 13). TWAR6.chr14 had decreased activity in the thylacine and increased activity in the wolf in both cell lines (adjusted *p*-value <1 x 10*^−^*^10^ in every replicate for both clades); TWAR5.chr7 had increased activity in the thylacine in the CNCCs and decreased activity in the wolf in both cell lines (adjusted *p*-value <0.002 in every replicate).

**Figure 6.**
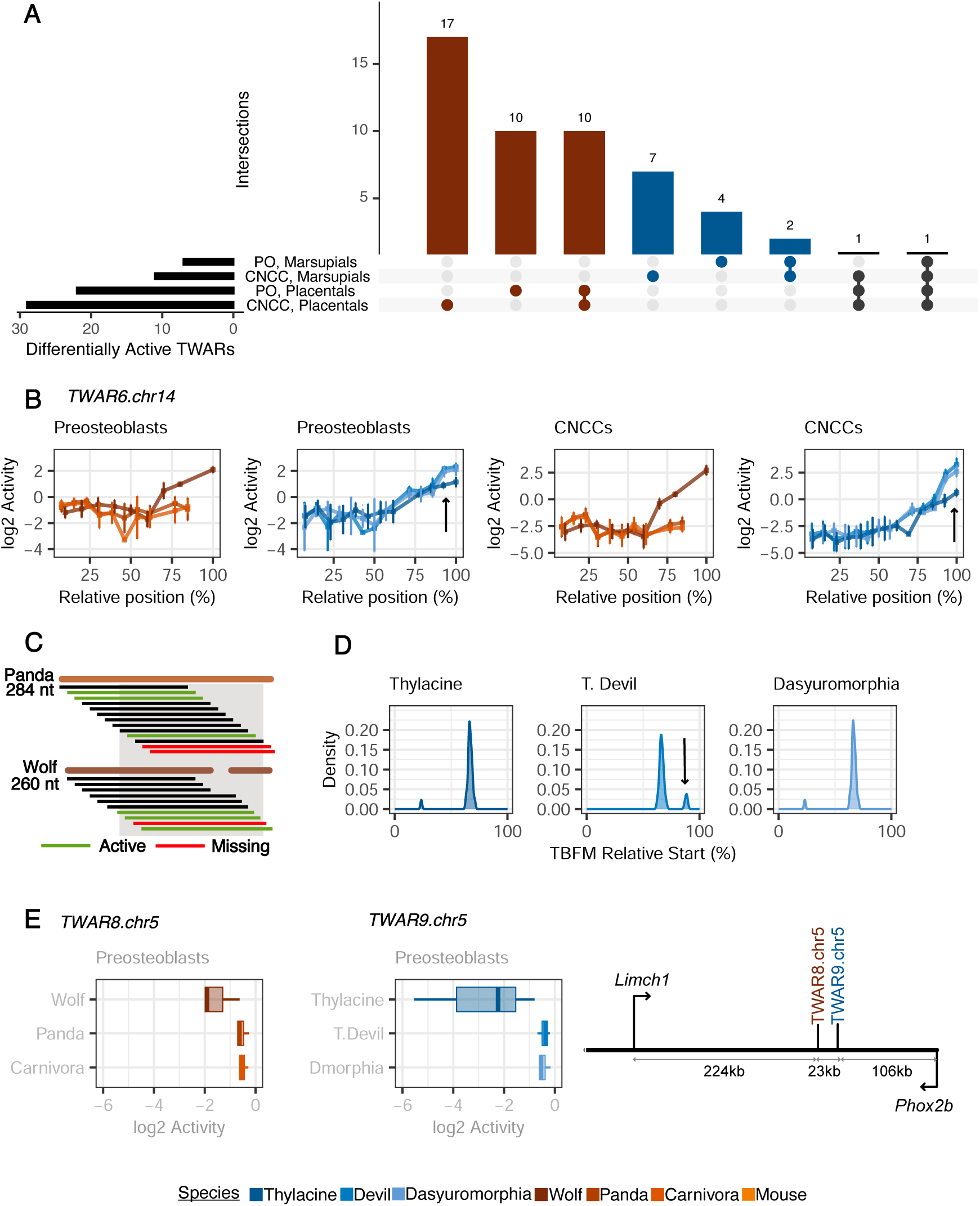
Convergent differential activity. **A.** Intersections of differentially active (DA) TWARs across the marsupials and placentals in both cell lines. **B.** Activity across tiles of TWAR6.chr14 for marsupials and placentals in each cell line. The arrows corresponds to tiles with increased activity in the Tasmanian devil which also have Tasmanian devil-specific biding motifs. **C.**The schematic compares the constitutive tiles of TWAR6.chr14 for the wolf and panda; it describes how although we have lost the last two panda tiles for this TWAR, panda sequences orthologous to wolf tiles that have significant activity are indeed present in our dataset. **D.** Distribution of binding motifs for the marsupial taxa have also been provided, with the Tasmanian devil-specific binding motif indicated by the arrow. **E.** Differential activity of TWAR8.chr5 in the placentals and TWAR9.chr5 in the marsupials. The diagram shows the proximity of these TWARs in the mouse genome and their two nearest genes, *Phox2b* and *Limch1*.

Within each clade, these patterns could be associated to distinct sequence features. For instance, increased activity in the Tasmanian devil and Dasyuromorphia TWAR6.chr14 relative to the thylacine was limited to their last two tiles; this correlated with a cluster of sequence differences in the last 13bp of their sequences, as well as TFBMs from the *Sp* family that mapped to these positions only in the Tasmanian devil sequence (Figure 6D). Likewise, gain-of-activity in the wolf TWAR6.chr14 was associated to a 22bp deletion which introduces 2 unique binding motifs for ZBT17 and ZN143 absent in the other therian taxa (Supplementary Figure 14). On the other hand, wolf TWAR5.chr7 had no detectable activity in either cell lines, likely due to a 29 bp deletion absent from the panda and the Carnivora sequence. TWAR6.chr14 is in the promoter region of *Chd8*, which is involved in the patterning and differentiation of cranial neural crest cells, while TWAR5.chr7 is located 64 kb downstream from *Rras2* and 73 kb upstream from *Copb1*, both of which have been linked to rare craniofacial pathologies [30–34]. Additionally, both TWARs overlap H3K27ac and H3K4me3 peaks in dunnart craniofacial tissue. As such, there is some evidence from our results that these TWARs may have functional roles in marsupials and placentals as well as divergent regulatory function in the thylacine and wolf due to specific sequence alterations; however, the lack of concordance in direction of effect complicates interpretability across clades.

Given the limited evidence for convergence at individual loci, we finally turned to searching for overlaps in molecular functions or pathways amongst genes near the other marsupial and placental DA TWARs. We found one instance of regulatory convergence at the gene-level — TWAR8.chr5 had significantly reduced expression in the wolf (adjusted *p*-value <1 x 10*^−^*^9^ in every replicate), whereas TWAR9.chr5 has significantly reduced expression in the thylacine (adjusted *p*-value <0.02 in all preosteoblast replicates; Figure 6E). The two TWARs are 23kb apart, and located upstream of *Phox2b* (+129kb for TWAR8.chr5, +107kb for TWAR9.chr5), which is essential for differentiation into the autonomic nervous system, including the cranial nerves and the cranial sensory ganglia [35, 36]. Ultimately, overlap in DA TWARs was limited, especially as the overall number of DA TWARs was much lower in marsupials, pointing towards a limited role for nucleotide-level genetic convergence in giving rise to the observed phenotypic similarities between the two taxa.

## Discussion

We have used an MPRA to evaluate the activity of 295 conserved non-coding elements that display convergent sequence acceleration in the marsupial thylacine and the placental wolf. To our knowledge, this is the first high-throughput functional dissection of non-coding elements from lineages this distant (divergence time *≈*160 MYA) [37]. Our approach demonstrates how MPRAs can be employed to ask complex evolutionary questions. Our strategy of including outgroup and ancestral taxa facilitated our comparisons by controlling for sequence decay observed over longer divergence times. For instance, we were able to identify tiles and TWARs with evidence of gene regulatory activity specific to the focal species as well as conserved patterns of activity across multiple TWARs active in all tested taxa, the latter suggestive of higher-level evolutionary constraint. We observed substantial functional divergence amongst the placental taxa but less so amongst the marsupials; however, both thylacine and wolf sequences frequently drove differential activity within their respective clades. However, only 2 TWARs were differentially active in both the marsupials and placentals; in both cases these had opposing directions of effect in thylacine and wolf. Within each clade we did observed multiple differentially active TWARs near key developmental genes such as *Lmo4* (TWAR27.chr3, marsupial DA), *Axin2* (TWAR14.chr11, placental DA) and *Hes1* (TWAR3.chr16, placental DA) [38–43].

The function of promoters and enhancers can be robust to sequence turnover [44–46]. Correspondingly, 17 (6.8% of the 250 TWARs captured in all taxa) displayed functional conservation of gene regulatory potential for all marsupials and placentals in one or both cell lines. Nonetheless, patterns of conservation and divergence were broadly associated with sequence features, as in previous MPRAs [29, 47, 48]. Within placentals, DA TWARs displayed greater sequence divergence than non-DA TWARs, and wolf sequences had divergent regulatory behaviour that overlapped with the gain and loss of TFBMs. This trend was not replicated in the marsupials, where we observed a paucity of differential activity.

Multiple factors factors can explain this observation. First, our finding of consistently small sequence distances between the thylacine and other marsupials for ostensibly thylacine accelerated regions could be rooted in how acceleration was initially defined. The TWARs were derived from a 61-species whole genome alignment of vertebrates that contained 41 placentals and 4 marsupials (opossum, wallaby, Tasmanian devil and thylacine). Of these, only the Tasmanian devil is within the same order as the thylacine (Dasyuromorphia), whereas the alignment includes two representatives from Carnivora besides the wolf [13]. Dense sampling, especially of closely related taxa, is integral to the characterisation of lineage-specific and convergent changes [49–51]. The undersampling of marsupials coupled with a placental-biased phylogeny likely overestimates sequence acceleration that is unique to the thylacine and potentially explains the lack of DA TWARs amongst marsupials. In addition, our analyses hint towards a limited impact of the murine *trans*-regulatory environment, with fewer TWARs active in marsupials than placentals, which could also obscure true signal. However, there are currently no validated craniofacial lineage cell lines from marsupials that would enable us to directly test this hypothesis, and our previous results have shown that the expression of craniofacial development genes is highly conserved between during early developmental in the mouse and the marsupial fat-tailed dunnart [28], suggesting that the possibility of widespread rewiring of the developmental regulatory network is unlikely.

In conjunction with this, the low sequence and functional divergence in craniofacial non-coding elements among marsupials could be reflective of their unique biology. Unlike placentals, marsupials give birth to highly altricial young that complete development *ex utero* in the maternal pouch. To enable the transition from newborn to suckling young, marsupials undergo accelerated development in both limbs and orofacial features [52–54]. This developmental requirement has been hypothesized to impose constraints on the evolution of early forming features in marsupials[55–60]. Though still subject to debate, this theory suggests that the craniofacial development machinery would also be more constrained, with functional craniofacial non-coding elements only able to tolerate sequence changes with no or subtle effects on regulatory activity.

Overall, we found limited support for the hypothesis that convergent acceleration within a given TWAR underlies phenotypic convergence between thylacine and wolf. This result aligns with previous findings on the lack of homoplasic substitutions in protein coding sequences between wolf and thylacine [9]. Our work instead suggests that morphological convergence between the thylacine and wolf arises through distinct genetic changes, and demonstrate that shared acceleration in sequence may not be reflective of shared regulatory function. Considering overall features of gene regulatory architecture, our findings are not unprecedented. As non-coding elements are characterized by a flexible organization and frequent redundancy [61, 62], identical genetic changes are not necessary to produce similar regulatory outcomes. Our decision to focus on TWARs that showed convergent evidence of acceleration was motivated by the desire to test the hypothesis of locus-level convergence. However, these intersecting elements were only a fraction of the complete set of thylacine accelerated regions (2.7%) and wolf accelerated regions (15.3%) defined by [13]. As suggested by our finding of parallel changes near *Phox2b*, a broader look at non-convergent, species-specific regulatory elements with these massively parallel approaches could perhaps be more fruitful in understanding the molecular origins of their convergent biology.

## Methods

### Library design

*Sequence preparation* We extracted previously identified TWARs from a 61-way whole genome alignment based on the mouse mm10 coordinates from [13] using maf_parse [63]. We obtained corresponding sequences for the thylacine, gray wolf, panda and Tasmanian devil with GNU grep for the species name and excluded elements if they were shorter than 90 bp in any taxa, as elements larger than this shared >80% sequence length between taxa. This left a total of 295 TWARs to be tested.

*Ancestral reconstruction* We used tree files for Carnivora and Dasyuromorphia from vertlife ( http://vertlife.org/phylosubsets) to generate independent tree models for each TWAR using the phyloFit utility from PHAST [63]. We then applied PREQUEL [63] (prequel i FASTA n s anc) with the output FASTA parameter to generate the ancestral sequences for each TWAR for Carnivora and Dasyuromorphia. Species included in the ancestral reconstructions are listed in **Supplementary Table 10** and **Supplementary Table 11**. For each reconstruction, we built a BLAST [64] database using the reconstructed sequences to compare sequence identity relative to the focal species. Fewer marsupial species (4) than placental taxa (11) were available as input for these reconstructions; consequently higher sequence sharing is observed between the Dasyuromorphia and the marsupial taxa (pairwise sequence identity 96% with thylacine and 94% with Tasmanian devil) than between Carnivora and the placental taxa (94.2% with the wolf and 85.4% with the panda).

*MPRA controls* We added mouse orthologs for each TWARs as controls for the murine cell lines used in our analysis, the MC3T3-E1 preosteoblasts [18] and O9-1 cranial neural crest cells [20].To obtain a set of preosteoblast-specific positive controls, we first downloaded a dataset of ChIP peaks for H3K27ac in mouse craniofacial (nose, maxillary and mandibular) tissue at E13.5 and E15.5 [65]. This was intersected with ChIP-seq peaks for *Runx2*, a master regulator of osteoblast formation in MC3T3-E1 preosteoblasts cells to obtain 194 overlapping regions (100-260 bp) [66]. We further intersected the craniofacial H3K27ac ChIP-seq peaks with a DNAse I hypersensitive site (DHS) sequencing dataset, also in MC3T3-E1 preosteoblasts cells, to get an additional 196 sequences (148-1,844 bp) [67]. Lastly, we added 6 enhancer sequences (98-215 bp) found to be highly active in human primary fetal osteoblasts in a previous MPRA [68]. For our cranial neural crest-specific positive controls, we extracted elements from the VISTA enhancer database known to be active in the branchial arch, cranial nerves and or the facial mesenchyme at E11.5 [69]. These were intersected with ChIP-seq peaks for *UTX*, involved in maintaining neural crest viability [70], as well as peaks for p300 in O91 mouse neural crest cells [71]. A set of 74 regions (196-3,785 bp in length) were found to be both strongly expressed in the craniofacial regions and intersecting with one or both of the ChIP-seq datasets. 170 bp scrambled sequences were generated as negative controls, and we used BLAT [64] and MEME FIMO [72] to confirm these sequences were not found in the mouse genome or predicted to have a transcription factor binding motif.

*Tiling* Given oligosynthesis constraints, we could only test 170bp of sequence at a time. When TWAR sequences were longer, we tiled across elements using a 10 bp sliding window, for a total of 22,110 tiles. Tiles can be identical when a TWAR is highly homologous across taxa; considering this redundancy, the library consisted of 21,798 unique tiles, of which 301 were shared between two species and 11 across three. This was carefully accounted for in our analysis.

### Library amplification and plasmid assembly

*PCR Amplification* The full sequence library of 21,798 unique sequences flanked by shared 15 bp adapter sequences at either end (5’ ACTGGCCGCTTGACG| CACTGCGGCTCCTGC 3’) was synthesized by Twist Bioscience. We followed a 2-step amplification protocol, as recommended in Gordon et al. [73]. An initial 8-cycle PCR was first performed on the single-stranded oligo pool with the *F_adapter* and *R_adapter primers*, with the NEBNext Ultra II Q5 Master Mix (M0544, NEB) and the double-stranded library was cleaned up with 1.8x AMPure XP beads (A63881, Beckman Coulter). We subsequently performed parallel 8-cycle PCR reactions with *F_oligo_PCR* and the degenerate *R_oligo_PCR*, to add a random 15 bp barcode to each oligo, as well as SfiI and BsiWI restriction sites and HiFi assembly sites for plasmid assembly. All primers used have been listed in **Supplementary Table 12**.

*First cloning step* The purified barcoded library and the pMPRA1 plasmid (https://www.addgene.org/49349/, gifted by Tarjei Mikkelsen) [74] were both digested with the SfiI restriction enzyme (R0123S, NEB) for 2 hours at 50C. The pMPRA1 backbone was gel-extracted (T1020, NEB) while the oligo library was purified with a QiaQuick PCR Purification Kit (28104, Qiagen) to minimize loss of library sequences. We then ligated the plasmid backbone and library overnight at 16C with T4 DNA ligase (M020S, NEB), using a molar insert:vector ratio of 1:3.

The cloned library was transformed into competent bacterial E. coli cells (JM109, Promega) with heat-shock (42C for 2 minutes). The transformed bacteria were plated on Luria-Bertani (LB) agar plates containing 100 ug/mL of ampicillin (BP1760-5, Thermo Fisher Scientific) and cultured overnight. Bacterial dilutions (1:10, 1:100 and 1:1000) were also plated and used to estimate the library complexity. Bacterial colonies were collected from plates and the plasmids were extracted with the ZymoPure Plasmid Maxi-Prep Kit (D4202, Zymo Research). The oligo and tag region were amplified from the competent plasmid library in 12-cycle PCR reactions (NEBNext Ultra II Q5 Master Mix), purified (QiaQuick PCR Purification Kit) and sent for NGS library preparation followed by amplicon sequencing (Illumina 2×150 bp) with Azenta Life Sciences. We performed this cloning step 3x, and sequenced each resulting library to a minimum depth of 100 million reads. The three cloned plasmid libraries were then combined for the following procedures.

*Second cloning step* We digested the oligo-pMPRA1 cloned plasmid with the BsiWI-HF restriction enzyme (R3553S, NEB) overnight at 37C, resulting in a linearised library with overhanging HiFi assembly sites. The reporter construct was synthesized as a gBlock^TM^ (Integrated DNA Technologies). This construct consisted of a minimal promoter in front of the open reading frame for the enhanced green fluorescent protein, with HiFi assembly overhangs flanking the sequence. We used the NEB HiFi DNA Assembly Master Mix (E2621S, NEB) to assemble the final plasmid, with a 1:2 ratio of vector to insert.

The reporter gene cloned plasmid was electroporated into 10-beta electrocompetent E. coli cells (C3020K, NEB) with the recommended settings (2.0 kV, 200 Ω and 25 *µ*F) and plated in LB agar plates containing 100 ug/mL of ampicillin. They were then cultured overnight, along with a dilution plate (1:10, 1:100 and 1:1000). We used our oligo-tag sequencing results and Poisson modelling to estimate the number of individual bacterial colonies required to capture the oligo library with a sufficient number of barcodes (approximately 20-25 x 10^6^ colonies). Bacterial cells were lifted from the cultured plates with LB, and the plasmid library was extracted with ZymoPure Plasmid GigaPrep Kit (D4202, Zymo Research).

### Cell culture, transfection and harvesting

*Preosteoblasts* We cultured MC3T3-E1 Subclone 4 preosteoblasts (CRL-2593, ATCC) in MEM *α* (no ascorbic acid) (A1049001, Gibco) with 10% Fetal Bovine Serum (FBS) (FBSFR-S00JF, Scientifix). Prior to transfection, cells were seeded in 7 x 150 mm plates. Cells were disassociated with TrypLE (12605010, Thermo Scientific) for passaging and harvesting.

*Cranial neural crest cells* O9-1 cranial neural crest cells (SCC049, Merck) were cultured in Complete Embryonic Cell Media (15% FBS and LIF) (ES-101, Sigma-Aldrich) supplemented with 100 ug/mL of recombinant human bFGF (100-18B, PeproTech). Prior to transfection, cells were seeded in 4 x 150 mm plates coated with 5 mg/ml of Matrigel base membrane matrix (354234, Corning). Cells were disassociated with 0.25% Trypsin EDTA (25200072, Thermo Scientific) for passaging and harvesting.

*Transfection* At 60-70% confluence, we transfected 17.5 ug of the competent plasmid library into each dish (total *≈*70-90 million per replicate, per cell line) with the jetOptimus transfection reagent (101000006, Polyplus). A jetOptimus reagent to DNA ratio of 1.5 was used. The media was replaced 4 hours post-transfection.

*Harvest* For each replicate, we disassociated cells 24 hours post-transfection, re-suspended them in dPBS (14190136, Thermo Fisher Scientific), and centrifuged them for 5 minutes (100xg MC3T3-E1; 300xg O9-1). The resulting pellets were re-suspended again in media and all the cell suspensions were combined. 10% of the total cell suspension was split equally into 2 microcentrifuge tubes, pelleted and stored at –80C, for use in DNA extraction. The remainder of the cell suspension was centrifuged and stored at –80C, for use in RNA extraction.

### DNA and RNA extraction and library preparation

*DNA extraction* We extracted DNA with the Qiagen DNeasy Blood and Tissue Kit (69504, Qiagen), following the spin-column protocol for purification from animal cells, including the optional treatment with RNase A (19101, Qiagen) to obtain RNA-free genomic DNA.

*RNA extraction and cDNA synthesis* We extracted RNA with the Qiagen RNeasy Midi Kit (75144, Qiagen), following the protocol for the isolation of total RNA from animal cells, including the on-column DNase digestion. The eluted RNA was then treated again with Turbo DNase (AM1907, Thermo Fisher Scientific). We used an input of between 25-50 ug of RNA per replicate library for first-strand synthesis with Superscript IV (18090200, Thermo Fisher Scientific), with the reverse transcription primer (*Reverse_transcription_primer_R*). We additionally treated the first-strand synthesis product with 1 uL of RNAse A (7000 units) (Qiagen, 19101) to remove excess leftover RNA.

*Library amplification* We performed qPCR reactions in triplicate per RNA and DNA library with SYBR Green 1 (628014, Thermo Fisher Scientific) to determine the number of cycles required for efficient amplification of the barcode sequence (15-17 cycles for gDNA libraries and 22-24 for cDNA libraries). The barcode region was then amplified in each gDNA and cDNA library with NEBNext Q5 Ultra II Master Mix and *GFP_F* and *Multipurpose_Buffer_R* primers. The amplicons were purified with the Qiagen PCR Minelute Kit (Qiagen, 28004) and sent for amplicon sequencing (Illumina 2×150 bp, >200M reads/library) with Azenta Life Sciences. All libraries were sequenced to a minimum depth of 200 million reads.

### MPRA Data Analysis

*Oligo-Barcode Association* We used Cutadapt (version 4.2) to trim sequencing adapters and filter the plasmid sequencing reads, keeping read pairs with Q>20 and length>75bp. The MPRAmatch pipeline developed by Ryan Tewhey (https://github.com/tewhey-lab/MPRA_oligo_barcode_pipeline/MPRAmatch.wdl) [75, 76] was used to associate barcodes with the oligos in our library. A custom python script was used in place of pull_barcodes.pl to accommodate our library design.

*Barcode counting* Cutadapt (version 4.2) was used to trim Illumina adapters from the barcode sequencing reads from the cDNA and pDNA libraries. Merged reads were then processed with the MPRAcount workflow from Ryan Tewhey (https://github.com/tewhey-lab/MPRA_oligo_barcode_pipeline/blob/master/MPRAcount.wdl) [75, 76]. The pipeline maps the reads to the created oligo-barcode dictionary in the previous step, generating a table of raw barcode counts for each cDNA and pDNA library. A custom python script was used in place of make_counts.py, to accommodate our library design. The number and percentage of reads that were successfully filtered and mapped is in **Supplementary Table 13**). We retained a lower read depth for the cDNA libraries of two of our cranial neural crest replicates; upon closer observation, sequences filtered out mapped to the mouse genome (mm10), and suggested the presence of excess residual endogenous RNA in these sequenced libraries.

*Normalization, filtering and QC* All analyses were in performed R (version 4.2.1) unless otherwise stated. We normalized the raw count data by library size to get counts per million (CPM), followed by a *log*_2_ transformation. We removed barcode observations with extremely low pDNA counts across replicates in each cell line, using the density distribution of the *log*_2_ normalized pDNA to determine the cutoff at which read counts stabilized (preosteoblasts: –5.50; CNCCs: –5.75) (Supplementary Figure 15), as in [29]. We further filtered out oligos with fewer than 10 barcodes in each cell line. We mapped oligos passing all filters to the untiled library with minimap2 (version 2.17) [77], and used samtools (version 1.16) [78] to calculate the percentage coverage of our untiled sequences (**Supplementary Table 14** and **Supplementary Table 15**).

*Activity calculation and testing* We calculated activity across all barcode observations as the ratio of normalized *log*_2_ cDNA CPM to normalized *log*_2_ pDNA CPM. We tested for statistically significant activity individually in each replicate. To do so, we applied a one-sided t-test per oligo (as in Uebbing et al. [29]), comparing the distribution of activity across barcodes to the overall mean activity observed in a replicate. Per replicate, we applied a Benjamini-Hochberg multiple testing correction. For each cell line, we defined active oligos as those that passed a false discovery rate (FDR) of 5% in at least one of three replicates and 10% in at least two (**Supplementary Table 16** and **Supplementary Table 17**). To summarize activity values for a library sequence we took the median activity across barcodes and averaged activity across the three replicates.

*Activity analysis* To compare activity distributions between TWARs, cell-line specific positive controls and negative controls, we first obtained the median *log*_2_ activity of the most active tile in each TWAR and control. We defined “active” TWARs and controls as having at least 1 active tile. In defining regions with marsupial- or placental-active TWARs, we required sequences to display good coverage (80% or higher) in at least 2 out of 3 species in the clade that was not active. GC content for each TWAR sequence was calculated with the seqinr package (version 4.2-36) in R. For each TWAR we determined the proportion of sites that differed between each pair of sequences (the pairwise sequence distance) with the ape package (version 5.8-1) in R (function dist.dna(), model = “raw”). Distance to nearest TSS in the mouse had been previously determined with GREAT in [13] (**Supplementary Table 18**). We classified TWARs within 1kb of the a TSS as promoters, and the rest as enhancers.

*Differential activity testing* For each TWAR, we first ranked the component tiles by average median *log*_2_ activity across replicates. The most active tile in a TWAR, for each species, is used for our comparisons. Per clade in each cell line, we took a subset of TWARs that were active in at least one of the three taxa and present in two of the three taxa. Thus in preosteoblasts, we tested differential activity in 110 and 134 TWARs for marsupials and placentals respectively; in CNCCs we tested differentially activity in 50 and 103 TWARs for marsupials and placentals respectively. We filtered out any TWARs where more than 1 taxa had low coverage (<80%). Following these steps, we used an ANOVA to assess for differential activity, with the taxon as the factor and *log*_2_ activity across barcodes as the response variable (as in Uebbing et al. [29]). We then applied a Benjamini-Hochberg multiple testing correction; differentially active TWARs were required to pass a FDR of 5% in at least one of the three replicates and 10% the at least two.

*Nearest gene analysis* Feigin et al. [9] had previously identified genes nearest to each TWAR in the mouse genome (mm10) with GREAT [79]. Here we designated TWARs within 1000 bp of a TSS promoters and the remainder as enhancers. Per cell line, we applied a t.test comparing the absolute distance to a TSS for inactive and active TWARs (MC3T3: *p*-value = 0.451; O91: *p*-value = 0.020) A caveat of this test however was that we only had information for the mouse genome; as such each orthologous TWAR was assigned the same absolute distance to a TSS. We thus also applied a linear modelling approach, with absolute distance to a TSS as our explanatory variable and proportion of active orthologs for a TWAR as our response variable (MC3T3: *p*-value = 0.629; O91: *p*-value = 0.128).

*Transcription Factor Motif Scan and Analysis* We used FIMO [72] to scan our oligo library for non-redundant vertebrate transcription factor binding motifs from the HOCOMOCO Core Mouse Collection (v11) [80] (**Supplementary File 3**). A *q*-value threshold of 0.10 was applied. When a single oligo was associated with multiple binding motifs for the same TF, we retained the motif with highest binding score. We then counted the number of unique transcription factor motifs discovered in each oligo, as well as the mean binding score across these motifs. Multiplying the TFBM counts and the mean binding score, we obtained a summary FIMO score. To compare FIMO scores for a TWAR across taxa (all six or within a clade), we divided its summary score by the maximum possible score for an ortholog of that TWAR.

*Marsupial craniofacial ChIP-seq* We used BLAST [81] to identify orthologous regions in the dunnart genome [28] using the Tasmanian devil sequence for each TWAR. Uniquely mapped regions from Tasmanian devil to dunnart were then blasted back to the dunnart genome to ensure reciprocal mapping. The dunnart coordinates for each mapped TWAR were extracted and intersected with existing dunnart craniofacial ChIP-seq peaks [28] using bedtools intersect [82]

## Data and code availability

Raw sequencing reads have been deposited to ENA under accession number PRJEB90363. All analysis scripts are available at https://gitlab.unimelb.edu.au/igr-lab/tycnmpra.

## Acknowledgements

The work was partly funded by Australian Research Council Discovery Project DP210100505 to AJP. AJP is also funded in part by Colossal Biosciences. St Vincent’s Institute acknowledges the infrastructure support it receives from the National Health and Medical Research Council Independent Research Institutes Infrastructure Support Program and from the Victorian Government through its Operational Infrastructure Support Program. The funders had no role in study design, data collection and analysis, decision to publish, or preparation of the manuscript. The authors thank Chengyu Deng and Severin Uebbing for advice on experimental protocols. We also thank members of the Gallego Romero lab for valuable comments on the manuscript.

## Competing interests

AJP is funded in part by Colossal Biosciences, a company engaged in de-extinction and genetic restoration projects, including work on the thylacine. However, Colossal Biosciences had no direct influence on this project design, data collection, analysis, interpretation, or conclusions presented in this paper.

## Author contributions

- Conceptualization: LEC, IGR, AJP
- Data Curation: NS
- Formal analysis: NS, LEC
- Funding Acquisition: AJP, IGR
- Investigation: NS
- Methodology: LEC, NS, DVM
- Resources: AJP, IGR
- Supervision: IGR
- Visualization: NS
- Writing – Original Draft Preparation: NS, IGR, LEC
- Writing – Review & Editing: All

## Supplementary Figures

**Supplementary Figure 1.**
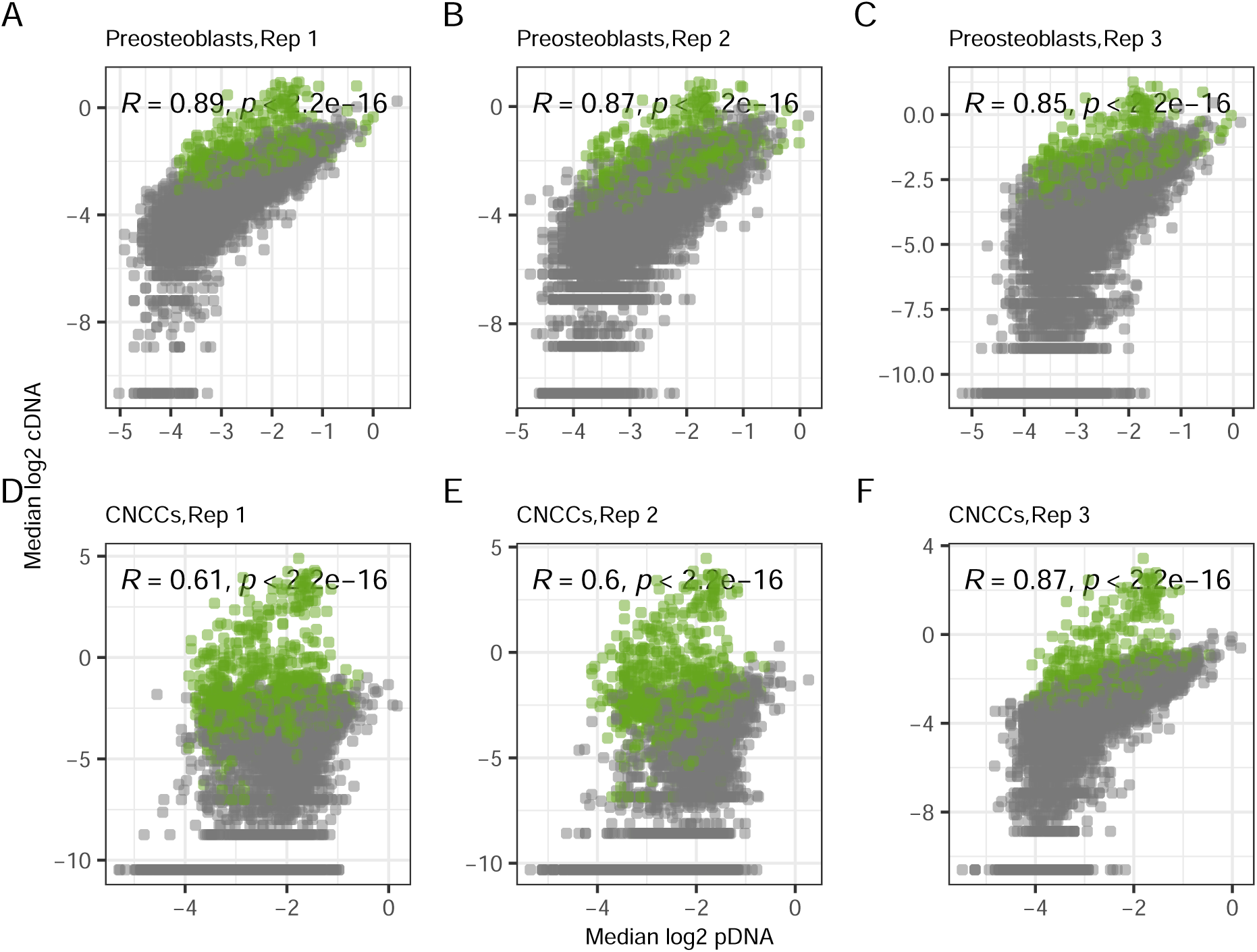
Scatterplot of median *log*_2_ CPM across barcodes for library sequences in the pDNA and cDNA libraries per cell line replicate, with Spearman’s rho correlations. Active sequences are indicated in green.

**Supplementary Figure 2.**
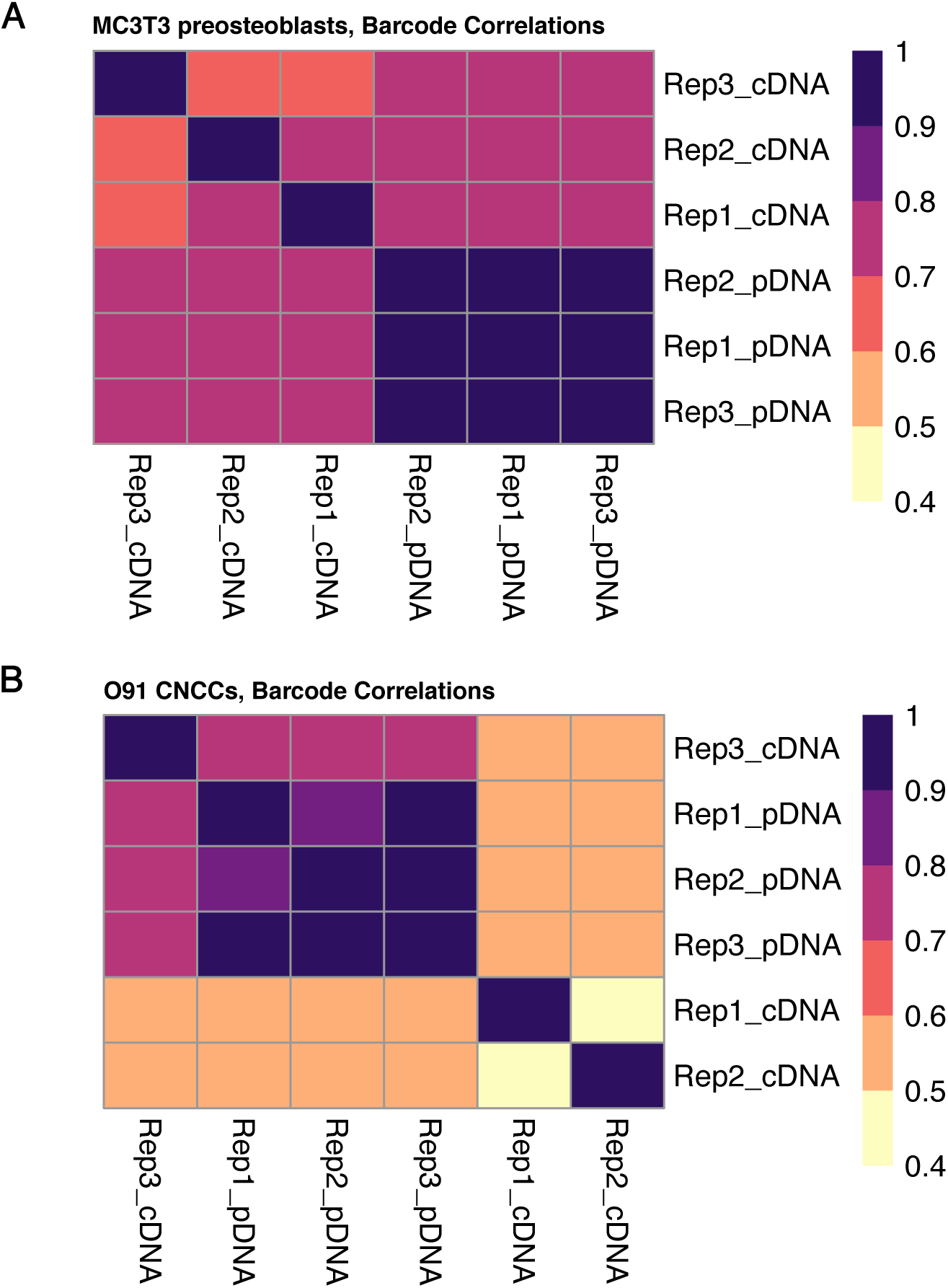
Correlations of barcode *log*_2_ CPM between all pDNA and cDNA libraries, per cell line replicate.

**Supplementary Figure 3.**
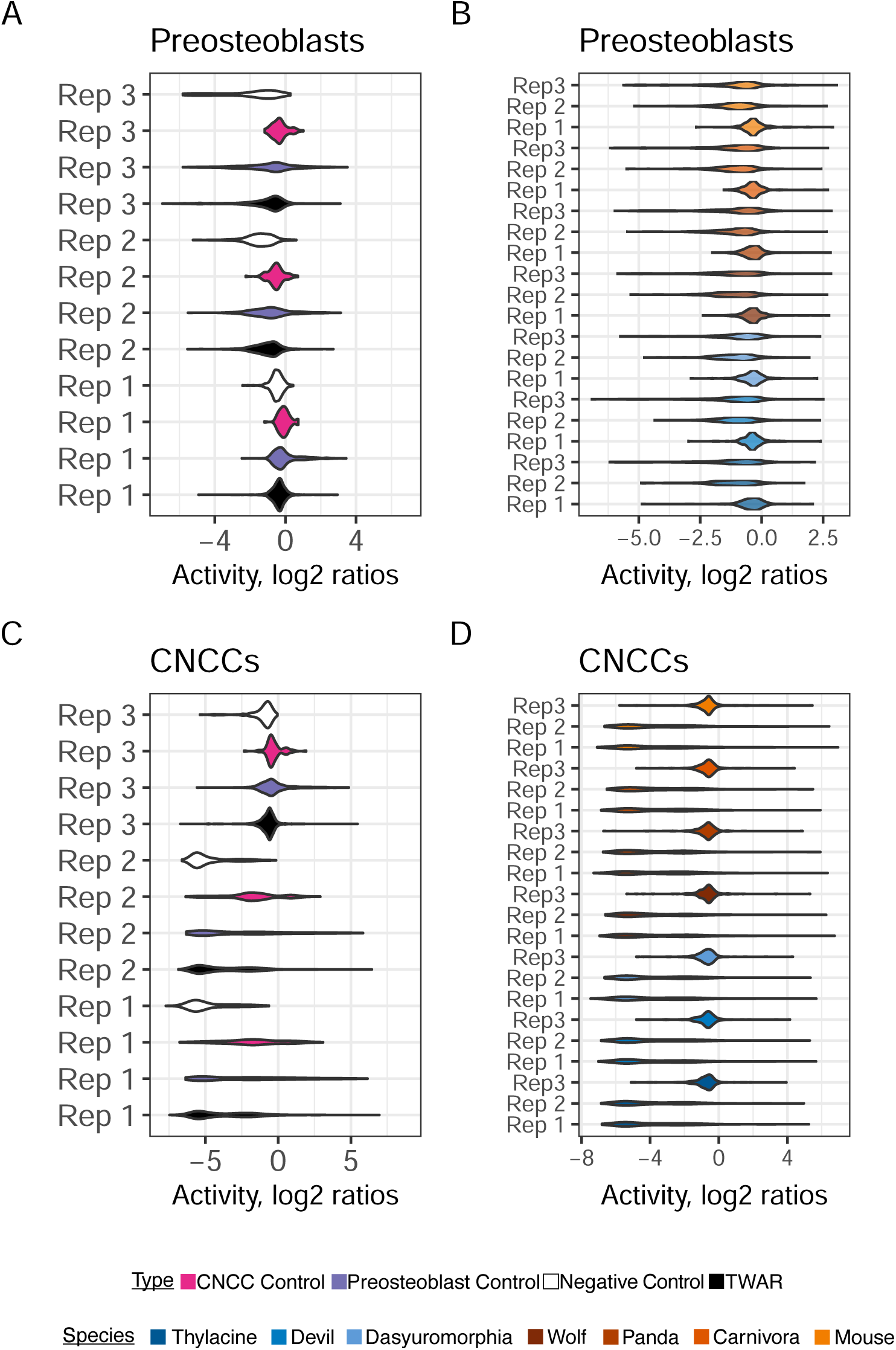
Distribution of activity values in each cell line replicate for **A and C.** all TWARs, cell-line specific positive controls and negative controls, and for **B and D.** TWARs from each of the six test taxa and the control mouse.

**Supplementary Figure 4.**
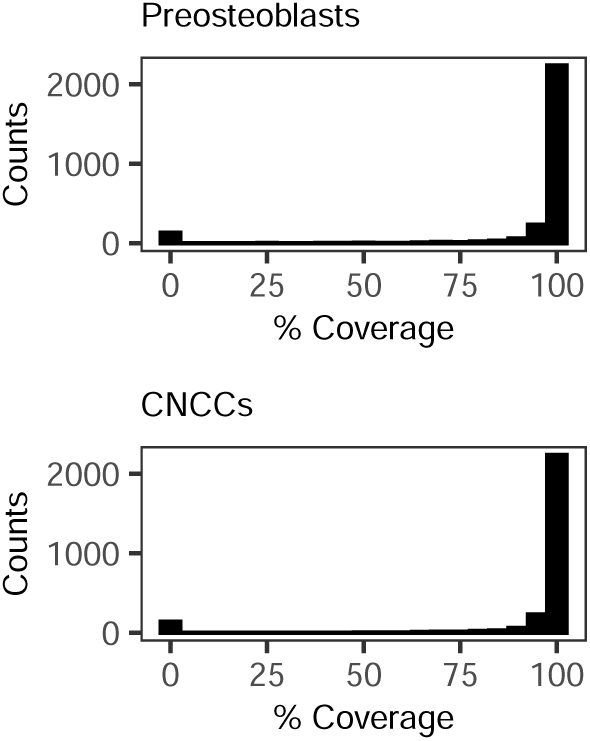
Percentage sequence coverage of un-tiled TWAR and control sequences that were captured in each cell line.

**Supplementary Figure 5.**
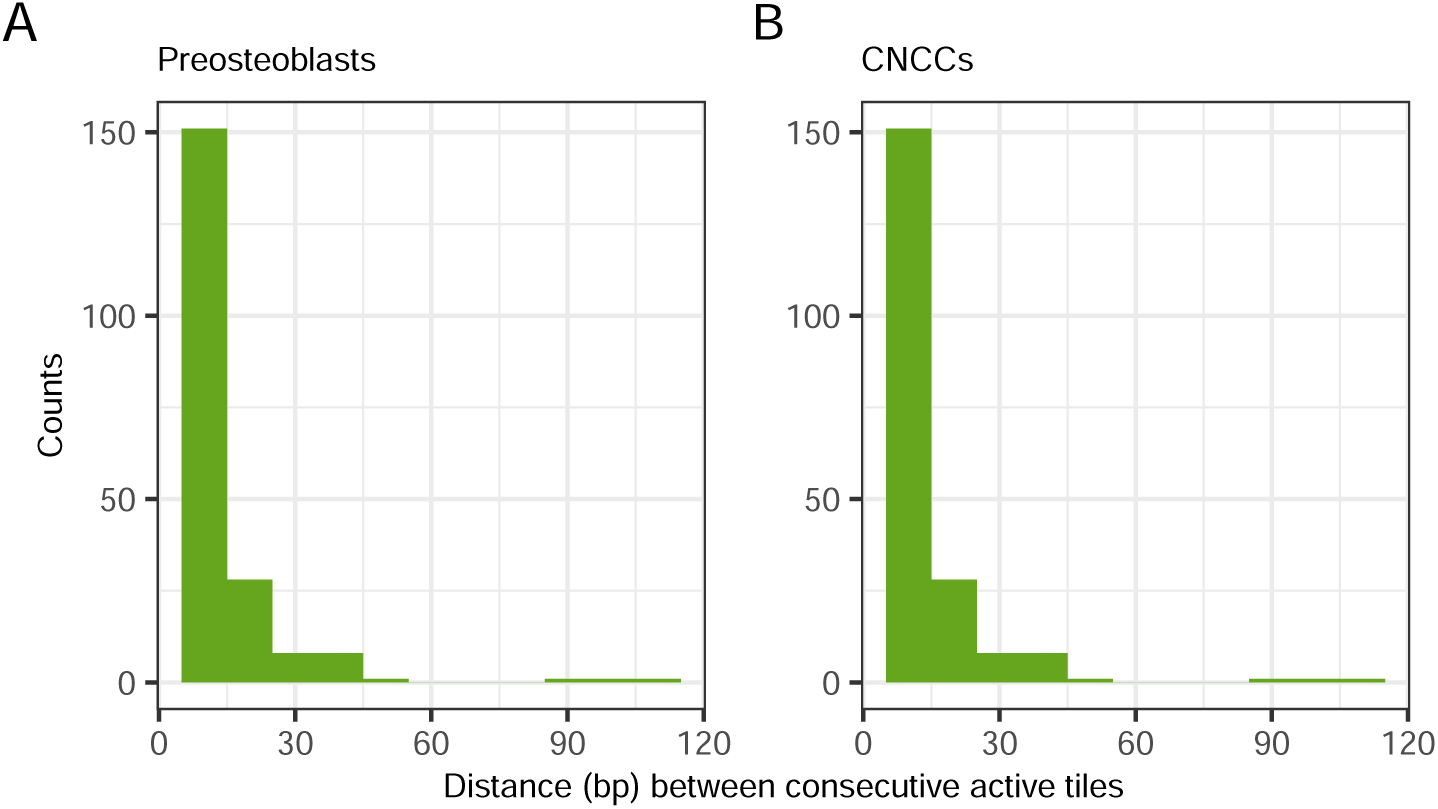
Distribution of distances between neighbouring active tiles for tiled TWARs with 100% coverage in our dataset.

**Supplementary Figure 6.**
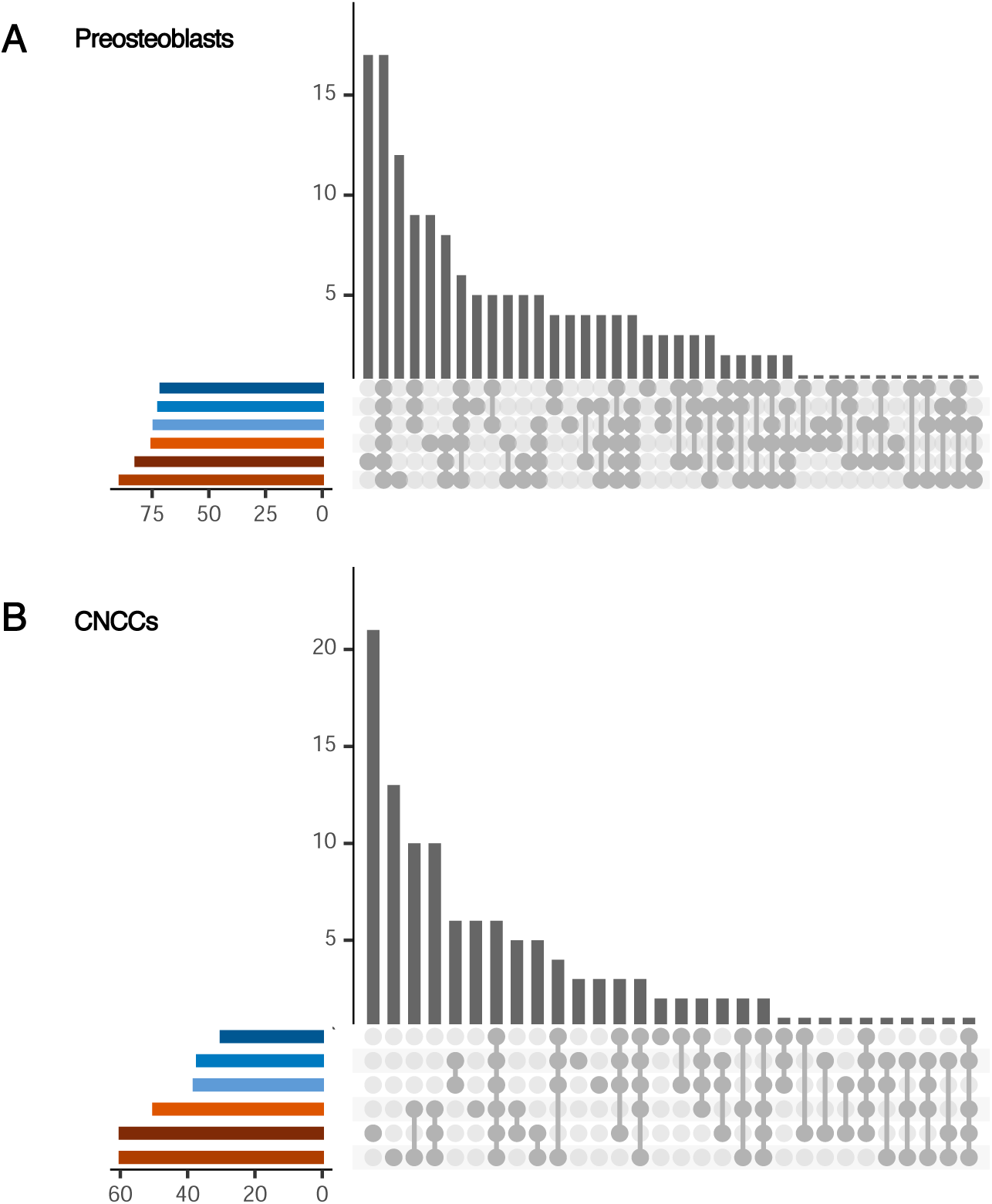
All intersections of active TWARs across species for each cell line.

**Supplementary Figure 7.**
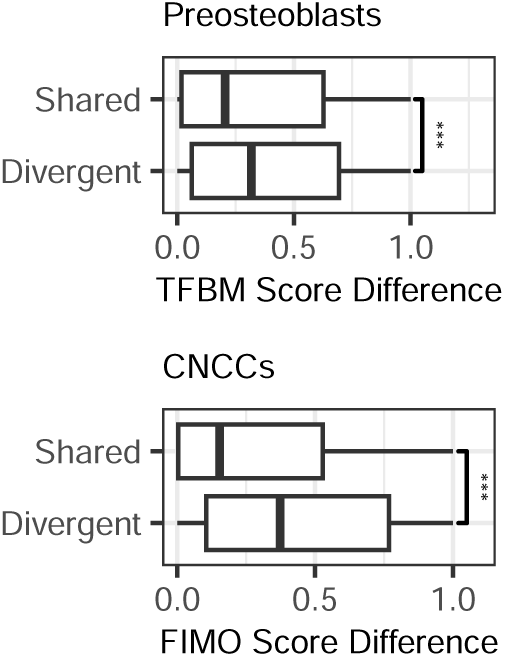
Differences in aggregated FIMO scores between pairs of orthologous TWARs with shared or divergent activity.

**Supplementary Figure 8.**
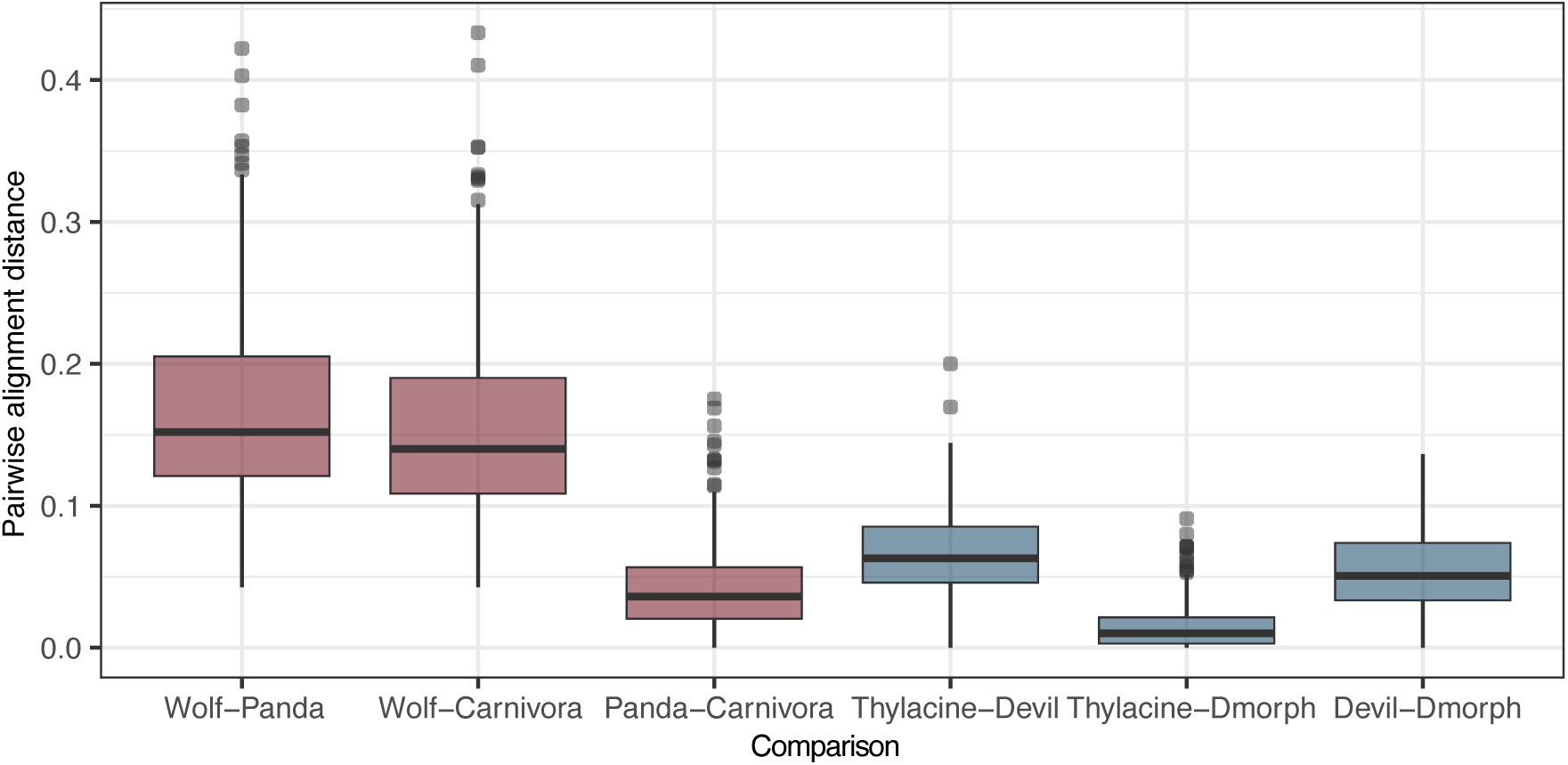
Distribution of pairwise sequence distances between orthologous TWARs.

**Supplementary Figure 9.**
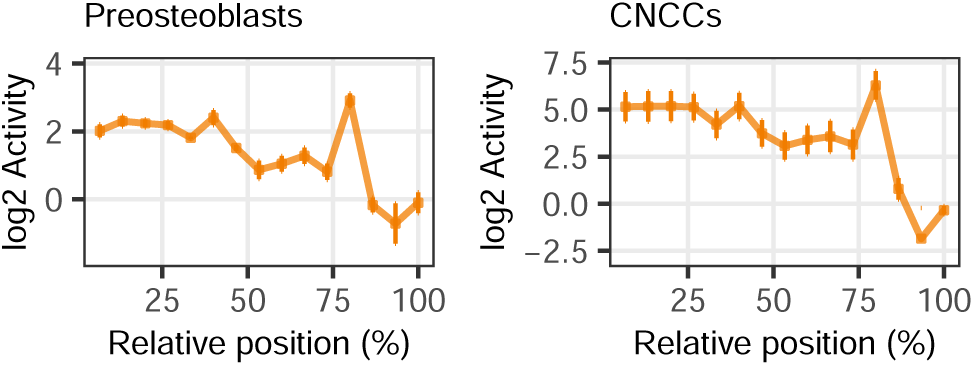
Activity across tiles of TWAR17.chr6 for mouse in each cell line.

**Supplementary Figure 10.**
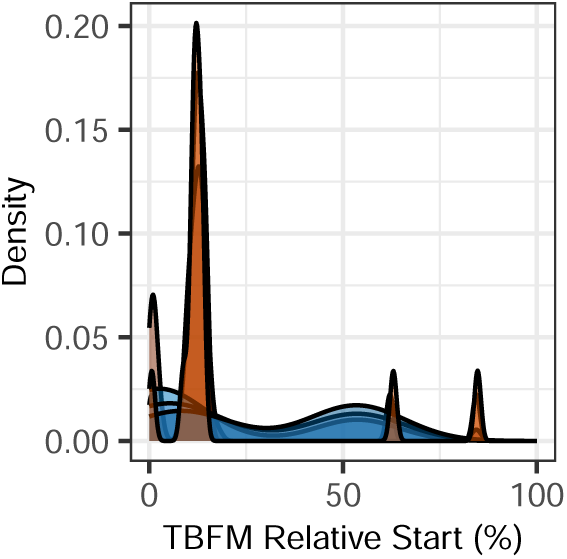
Positional distribution of transcription factor binding motifs for all six orthologs of TWAR17.chr6.

**Supplementary Figure 11.**
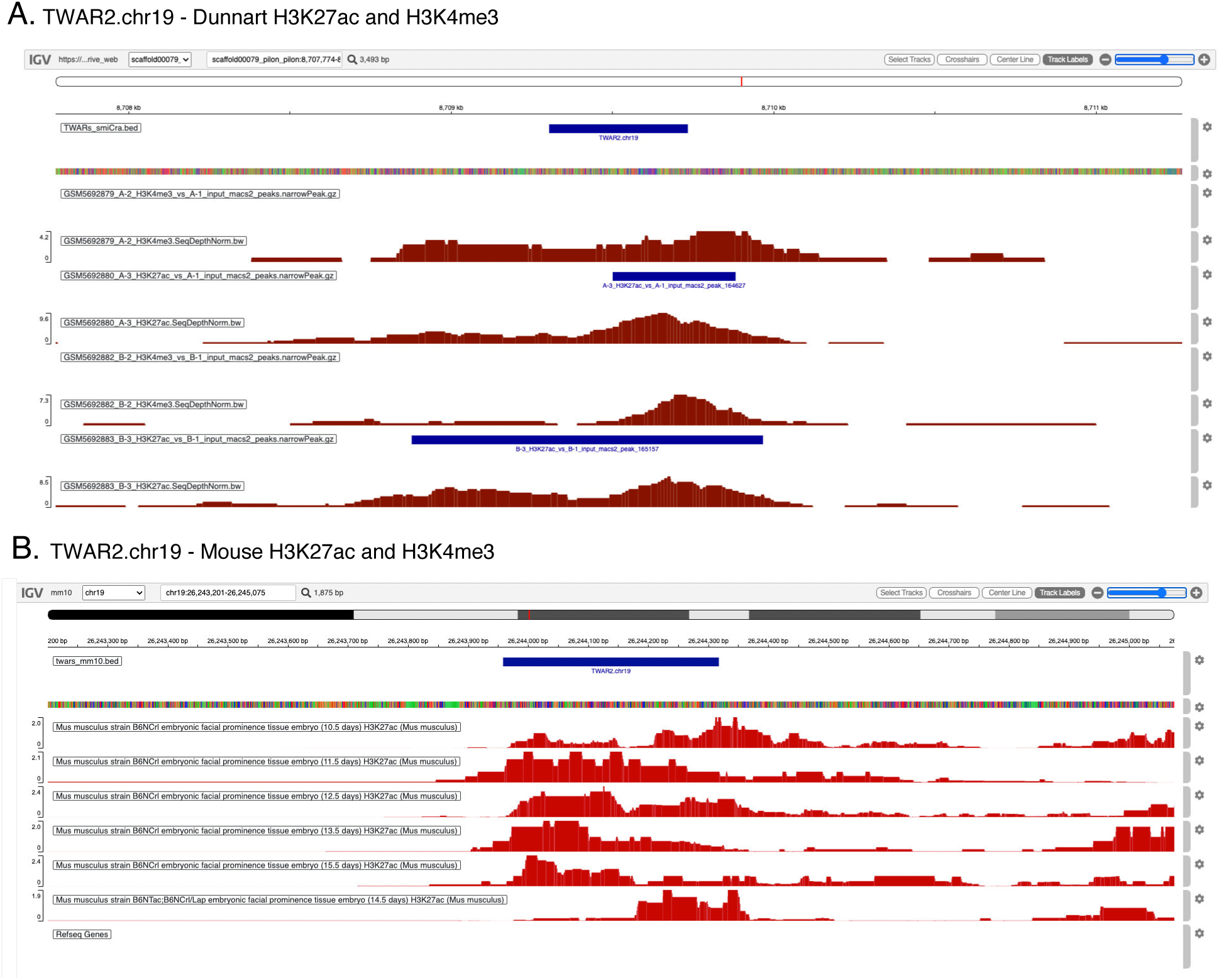
H3K27ac and H3K4me3 peaks for TWAR2.chr19 in **A.** dunnart pouch young (less than 24 hours old) craniofacial tissue from [83] and **B.** mouse embryonic (E10.5 to R15.5) craniofacial tissue from the ENCODE database [27].

**Supplementary Figure 12.**
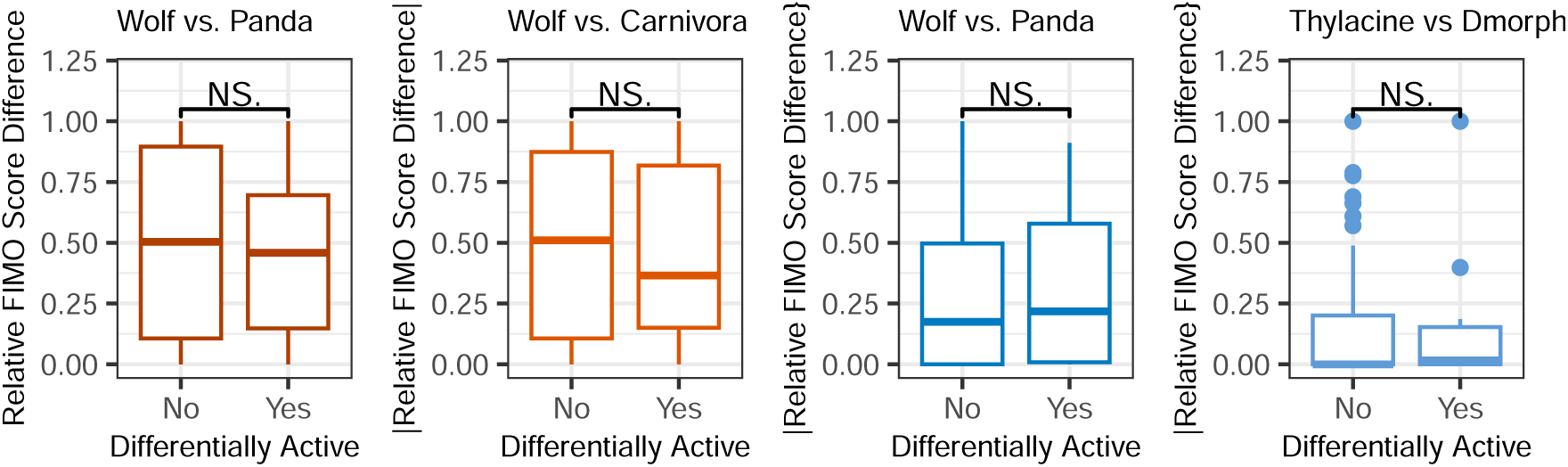
Differences in relative FIMO scores between the test and outgroup or ancestral orthologs for differentially active and non-differentially active TWARs.

**Supplementary Figure 13.**
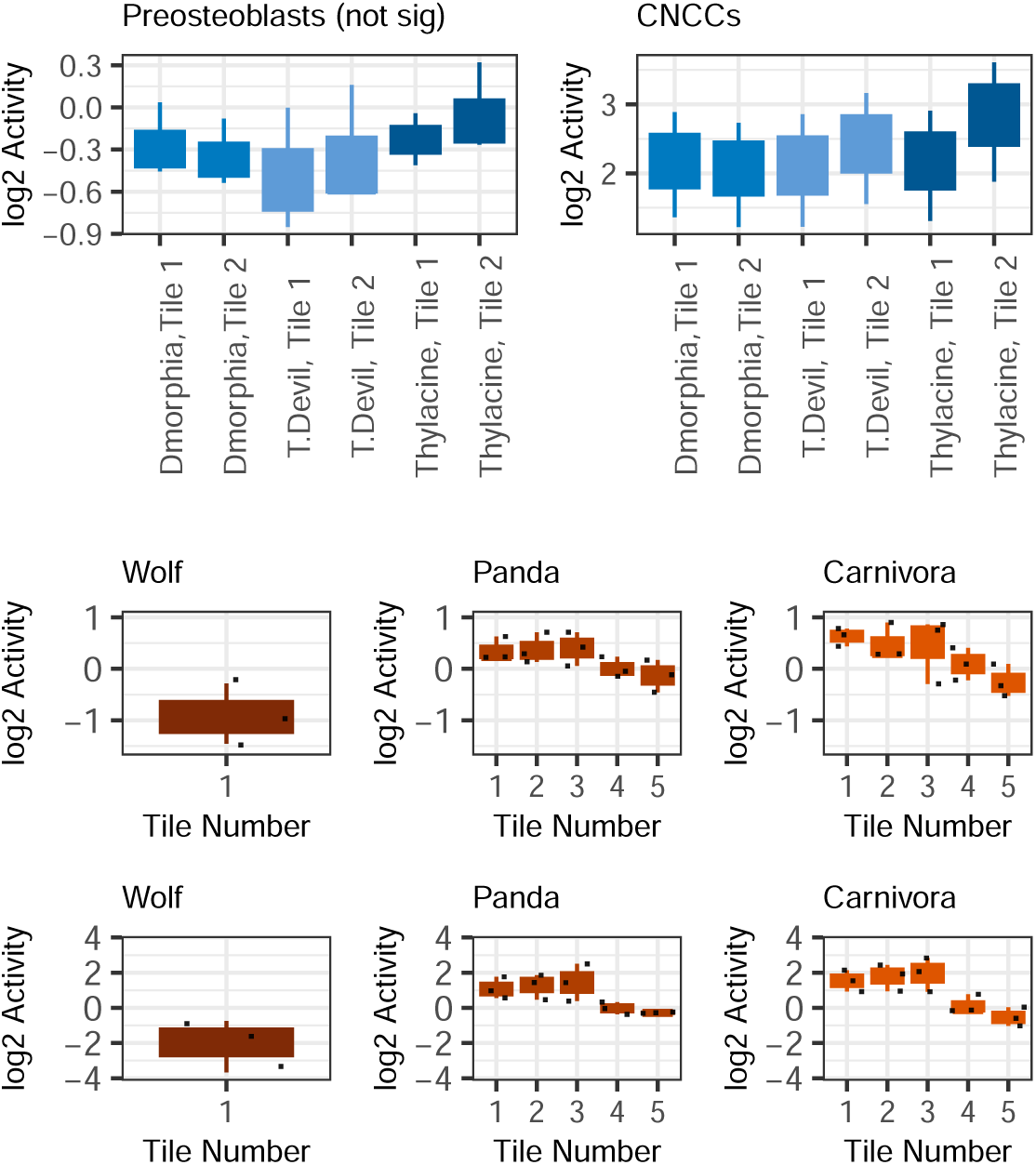
TWAR5.chr7 **A**. Distributions of activity for the marsupials, with each taxon containing two tiles. **B.** Activity across tiles for the placentals; the wolf sequence contains a large deletion.

**Supplementary Figure 14.**
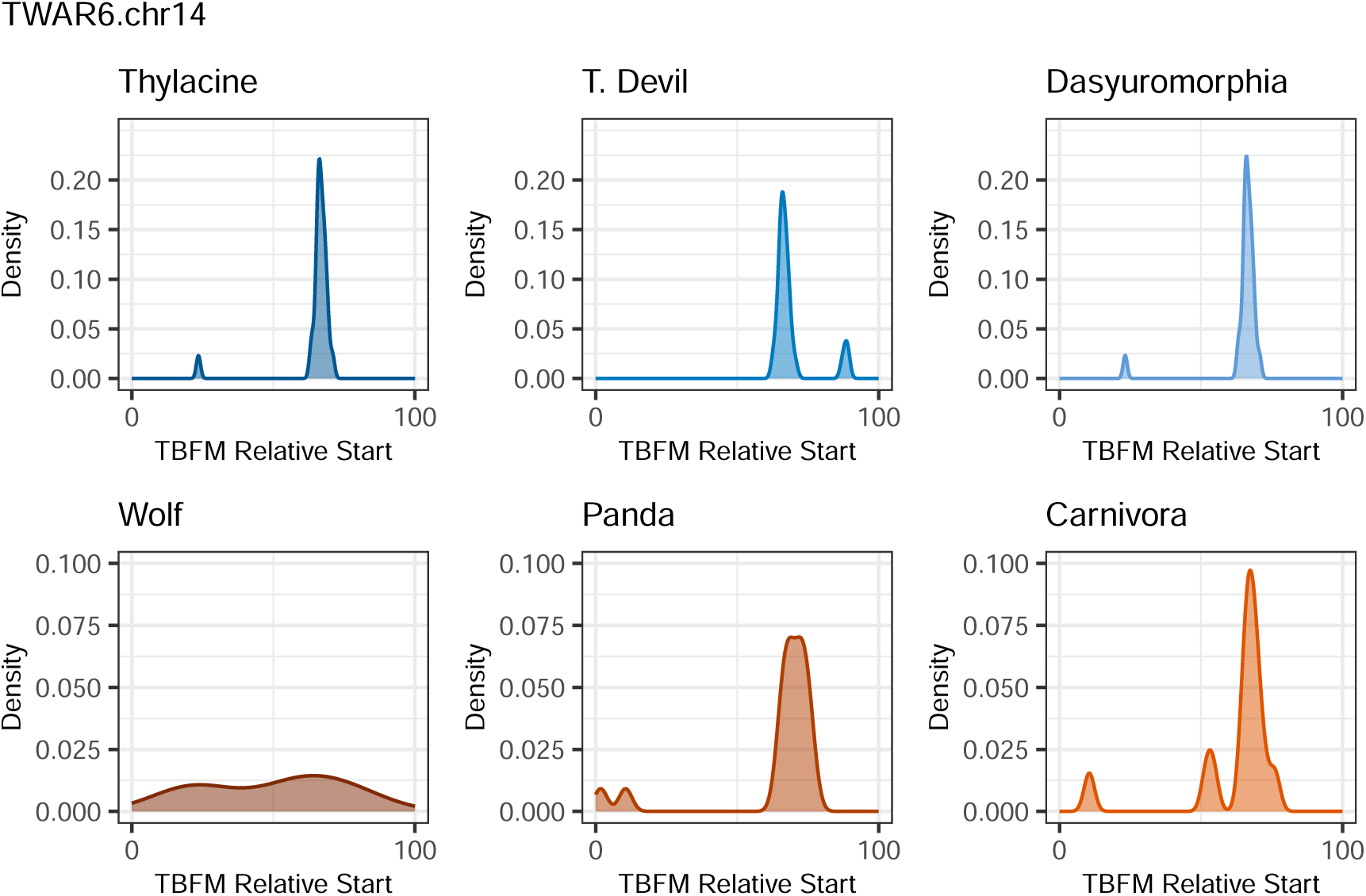
Positional distribution of binding motifs for all six orthologs of TWAR6.chr14.

**Supplementary Figure 15.**
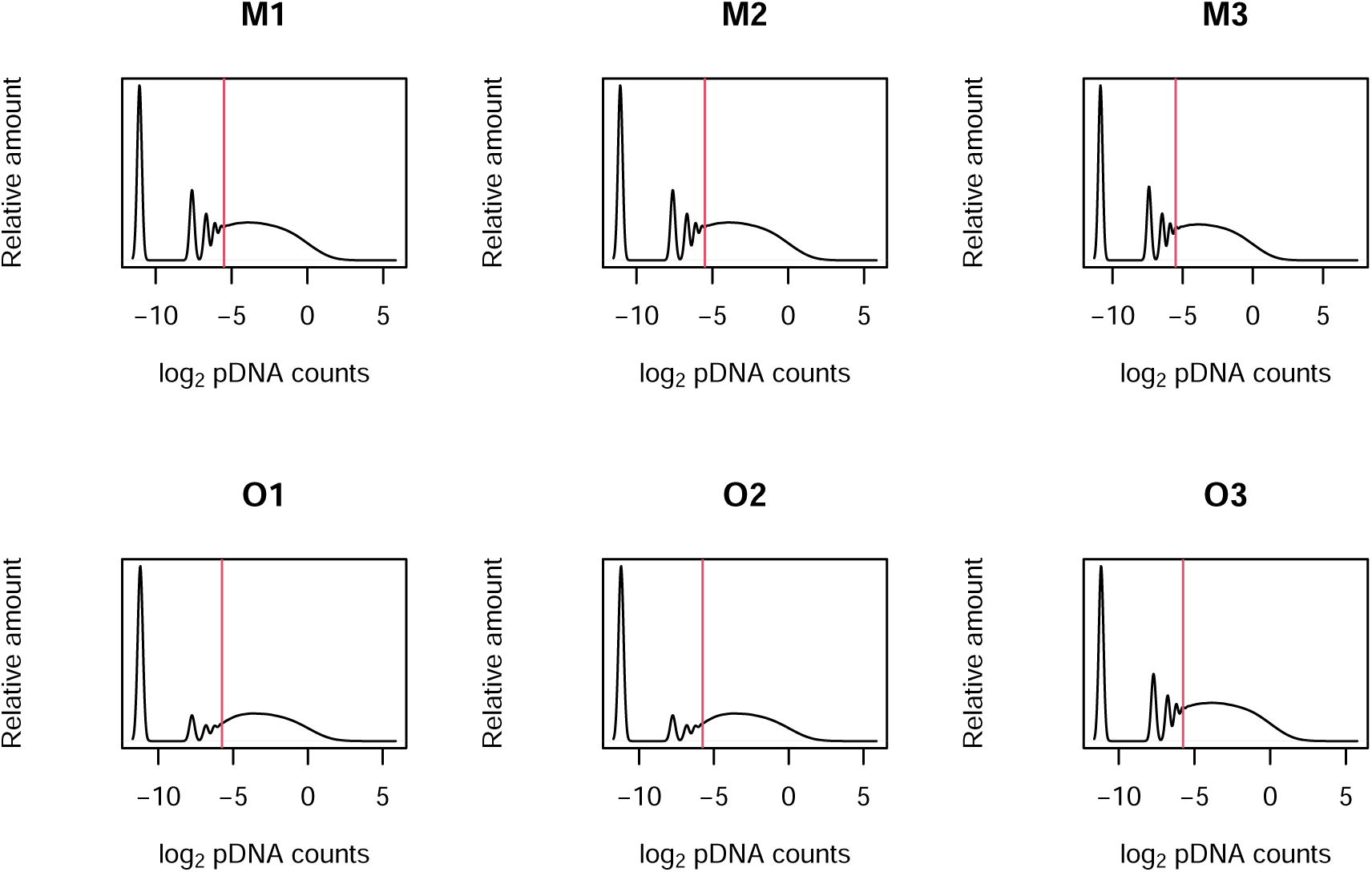
Distribution of *log*_2_ CPM for barcode sequences in the pDNA of each cell line replicate (M — MC3T3-E1 preosteoblasts; O — O91 CNCCs. The line is the threshold of filtering and barcodes with pDNA values lower than the threshold were removed.

## Supplementary Tables

**Supplementary Table 1**: All tested sequences

**Supplementary Table 2**: Pairwise t-tests comparing the activity distributions between the different categories of elements (cell-line positive controls, negative controls and TWARs) within each cell line replicate.

**Supplementary Table 3**: Pairwise t-tests comparing the activity distributions between different species within each cell line replicate.

**Supplementary Table 4**: Two-proportion tests comparing difference in proportion of active TWARs between each pair of species, for both cell lines.

**Supplementary Table 5**: List of TWARs found to be active in all species, all marsupials, or all placentals.

**Supplementary Table 6**: ANOVA for differential activity among marsupials in the MC3T3-E1 pre-osteoblast cells.

**Supplementary Table 7**: ANOVA for differential activity among placentals in the MC3T3-E1 pre-osteoblast cells.

**Supplementary Table 8**: ANOVA for differential activity among marsupials in the O91 cranial neural crest cells.

**Supplementary Table 9**: ANOVA for differential activity among marsupials in the O91 cranial neural crest cells.

**Supplementary Table 10**: Genomes used in reconstruction of the marsupial carnivore ancestor (Dasyuromorphia).

**Supplementary Table 11**: Genomes used in reconstruction of the placental carnivore ancestor (Carnivora).

**Supplementary Table 12**: List of primers.

**Supplementary Table 13**: Barcode sequencing reads post-filtering for each RNA and DNA library.

**Supplementary Table 14**: Coverage of each TWAR in the MC3T3-E1 pre-osteoblast cells.

**Supplementary Table 15**: Coverage of each TWAR in the O91 cranial neural crest cells.

**Supplementary Table 16**: Results of activity tests, MC3T3-E1 pre-osteoblast cells.

**Supplementary Table 17**: Results of activity tests, O91 cranial neural crest cells.

**Supplementary Table 18**: Mouse coordinates for and nearest genes to the thylacine and wolf accelerated regions.

## Supplementary Files

**Supplementary File 1**: All activity-by-tile plots for TWARs observed to be active in all six species.

**Supplementary File 2**: All activity-by-tile plots for TWARs displaying marsupial or placental-specific activity.

**Supplementary File 3**: Output of motif-finding with FIMO [72] for all TWARs.

