## Supplementary File 2 for "Dissecting functional regulatory convergence over 160 million years of therian evolution"

**Preosteoblasts, marsupial active, TWAR14.chr5**

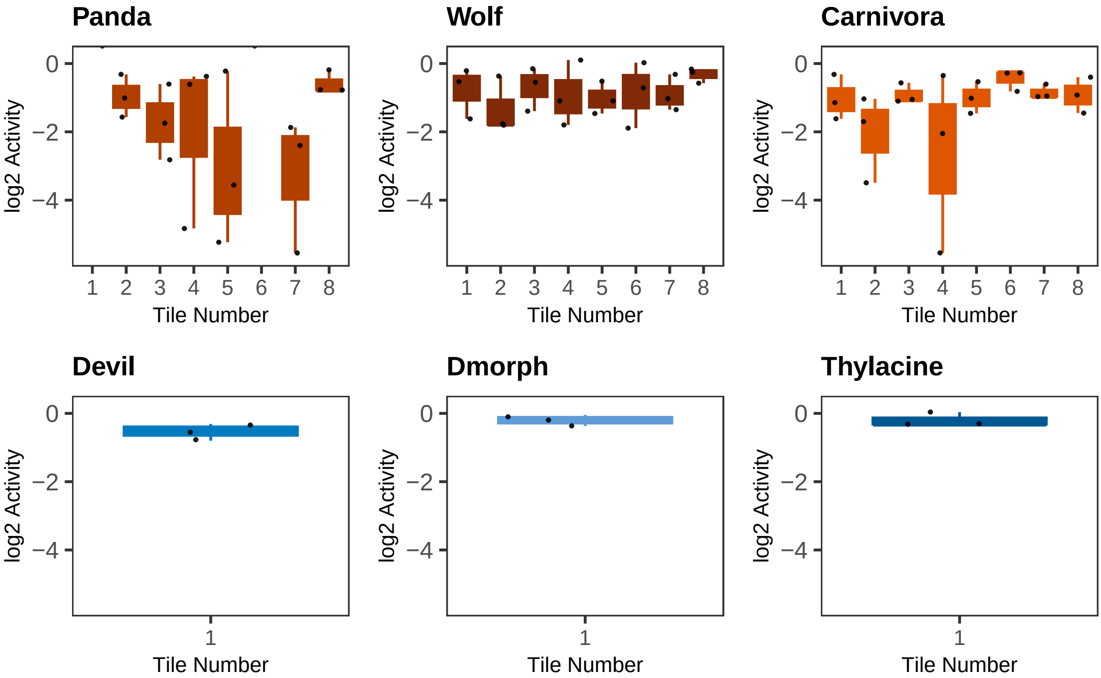

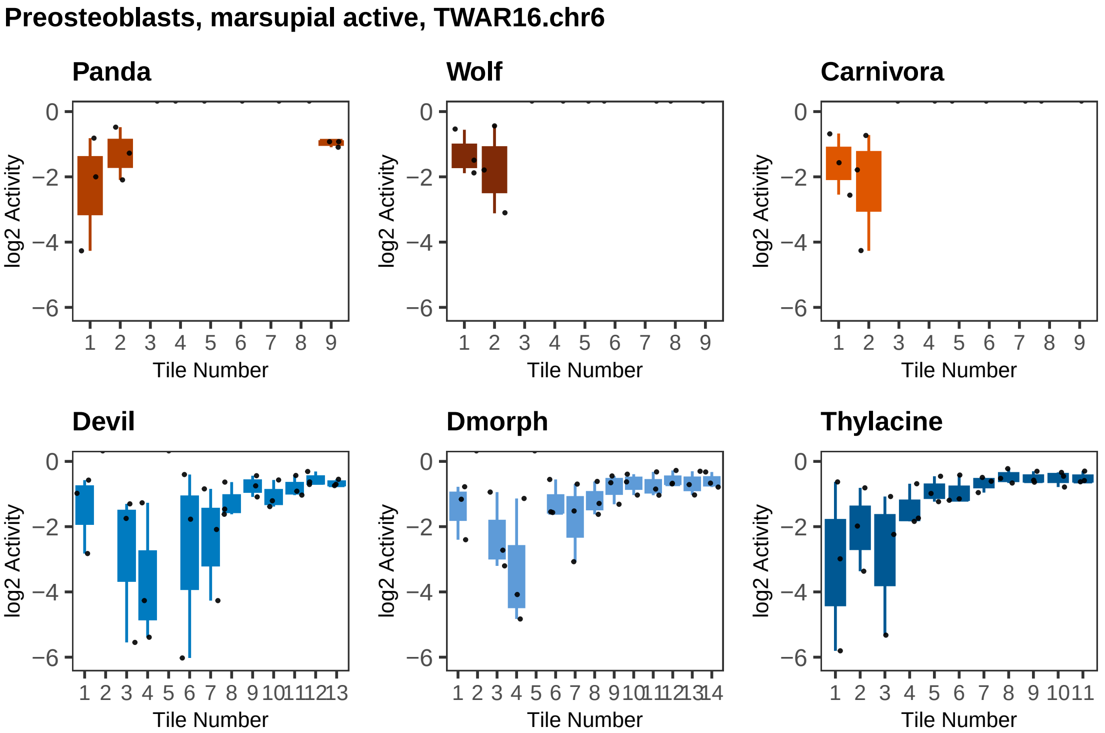
Preosteoblasts, marsupial active, TWAR16.chr6
Panda
Wolf
Carnivora
Devil
Dmorph
Thylacine
log2 Activity
Tile Number
0
−2
−4
−6
1 2 3 4 5 6 7 8 9
1 2 3 4 5 6 7 8 9 10 11 12 13
1 2 3 4 5 6 7 8 9 10 11 12 13 14
1 2 3 4 5 6 7 8 9 10 11

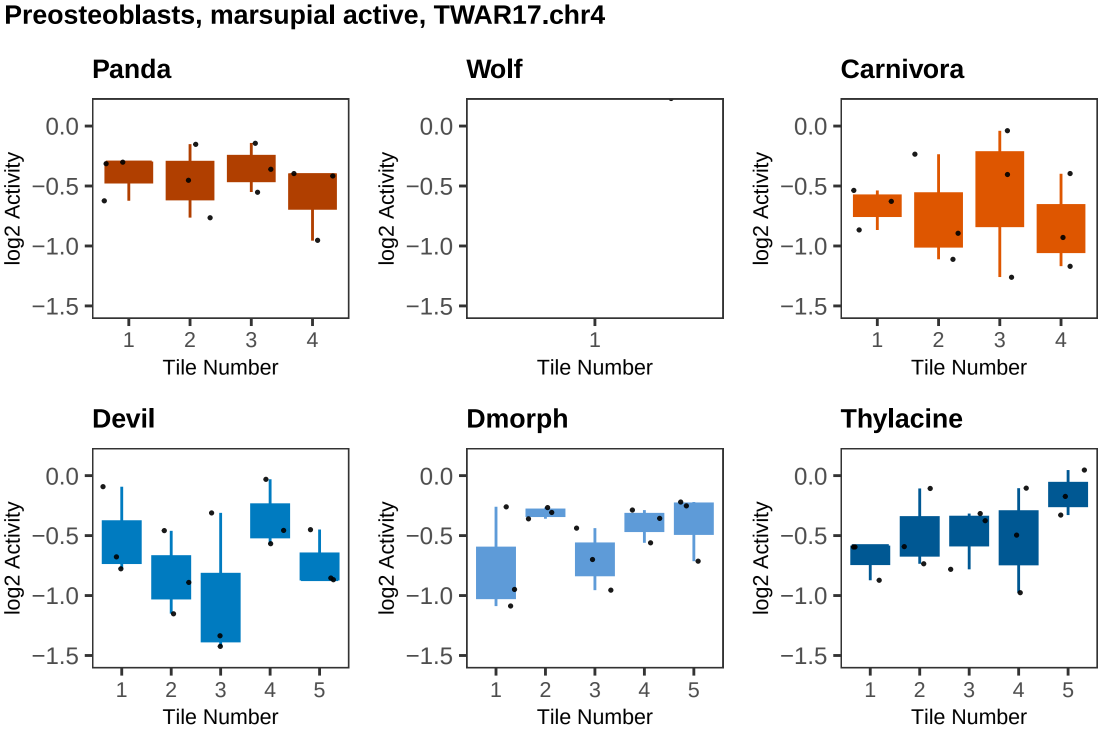
Preosteoblasts, marsupial active, TWAR17.chr4
Panda
Wolf
Carnivora
Devil
Dmorph
Thylacine
log2 Activity
Tile Number
0.0
−0.5
−1.0
−1.5
1
2
3
4
5

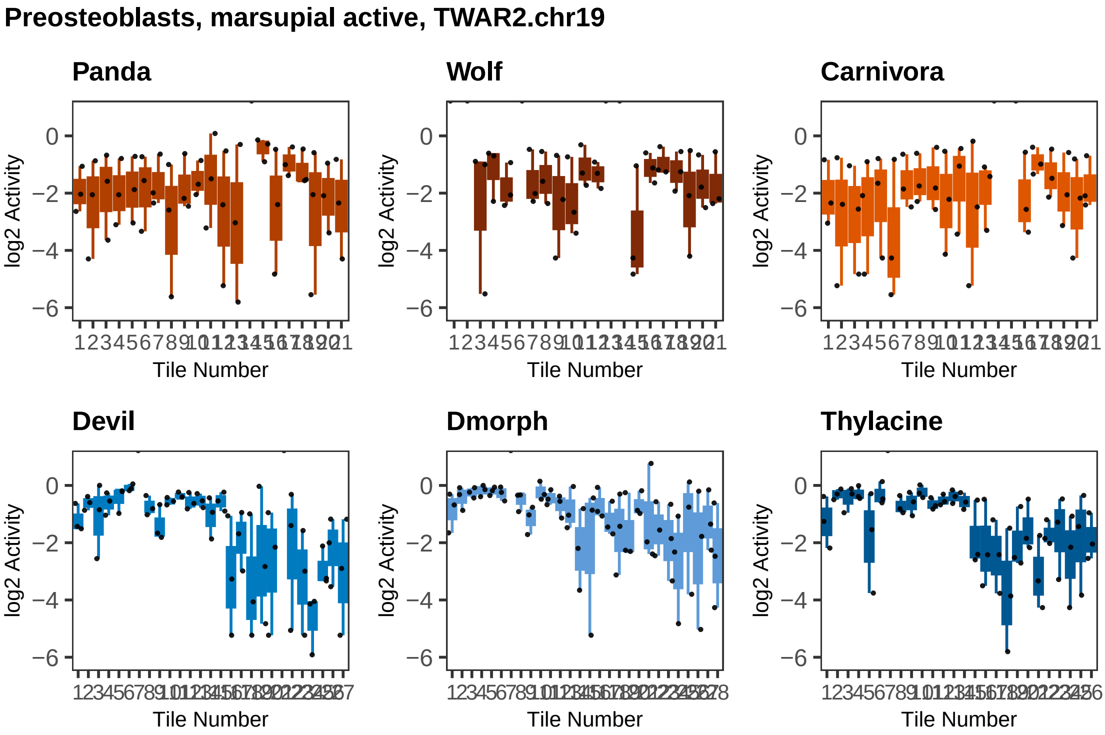
Preosteoblasts, marsupial active, TWAR2.chr19
Panda
Wolf
Carnivora
Devil
Dmorph
Thylacine
log2 Activity
Tile Number
0
−2
−4
−6

**Preosteoblasts, marsupial active, TWAR27.chr3**

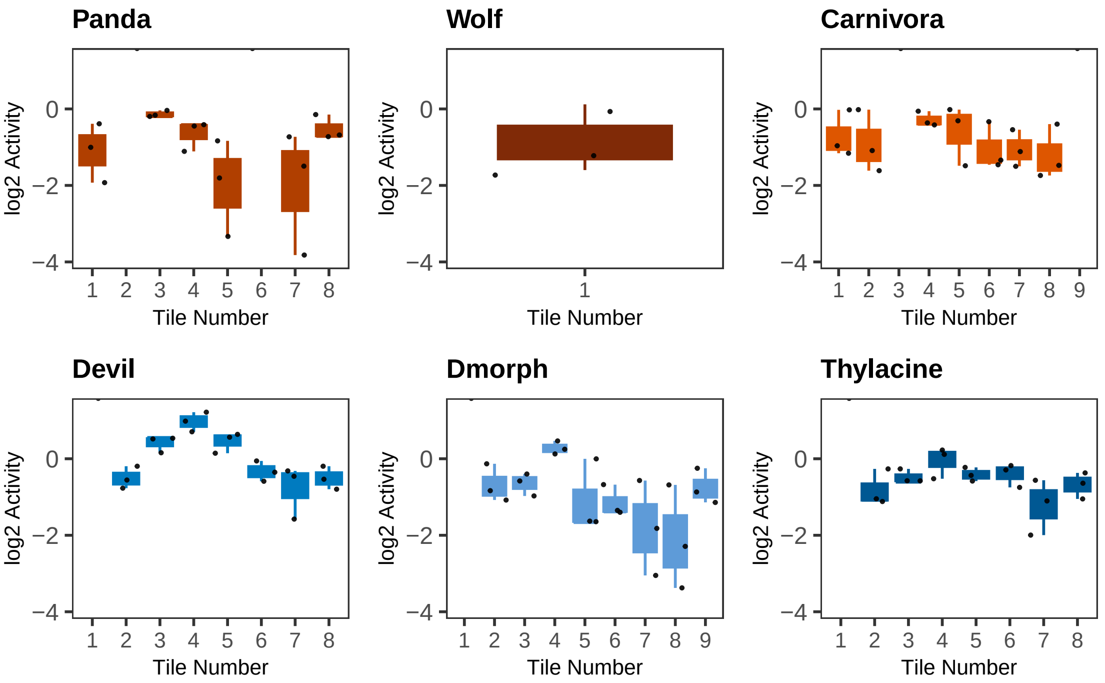

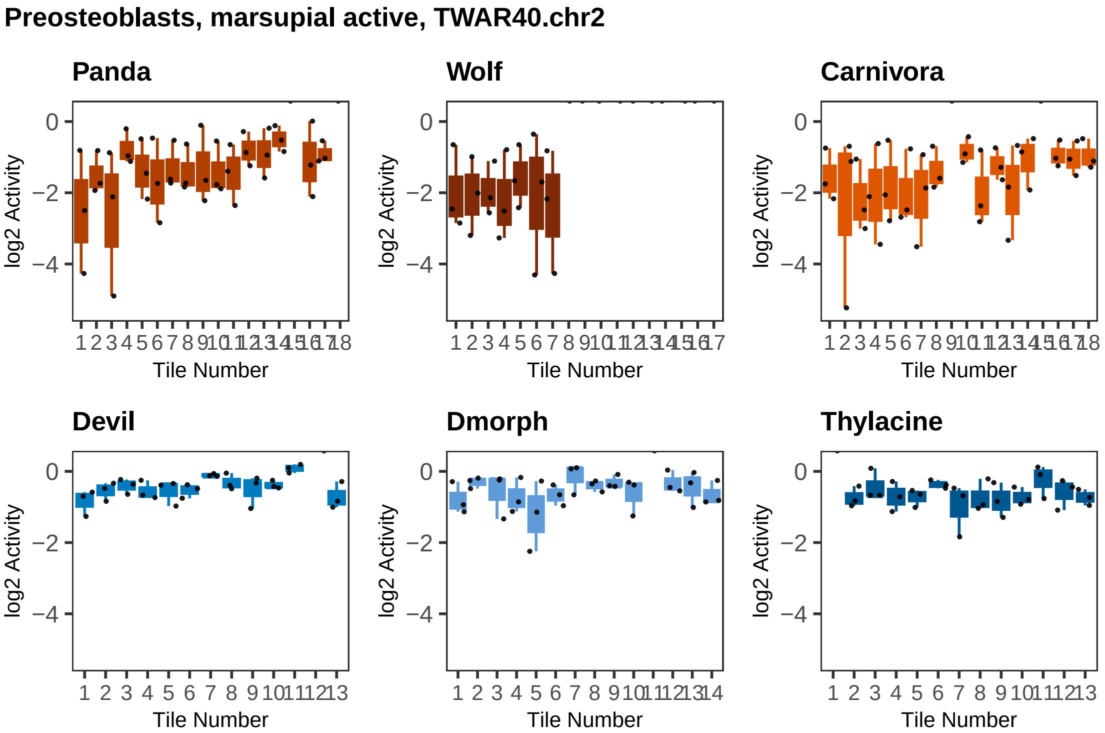
Preosteoblasts, marsupial active, TWAR40.chr2
Panda
Wolf
Carnivora
Devil
Dmorph
Thylacine
log2 Activity
Tile Number
0
−2
−4

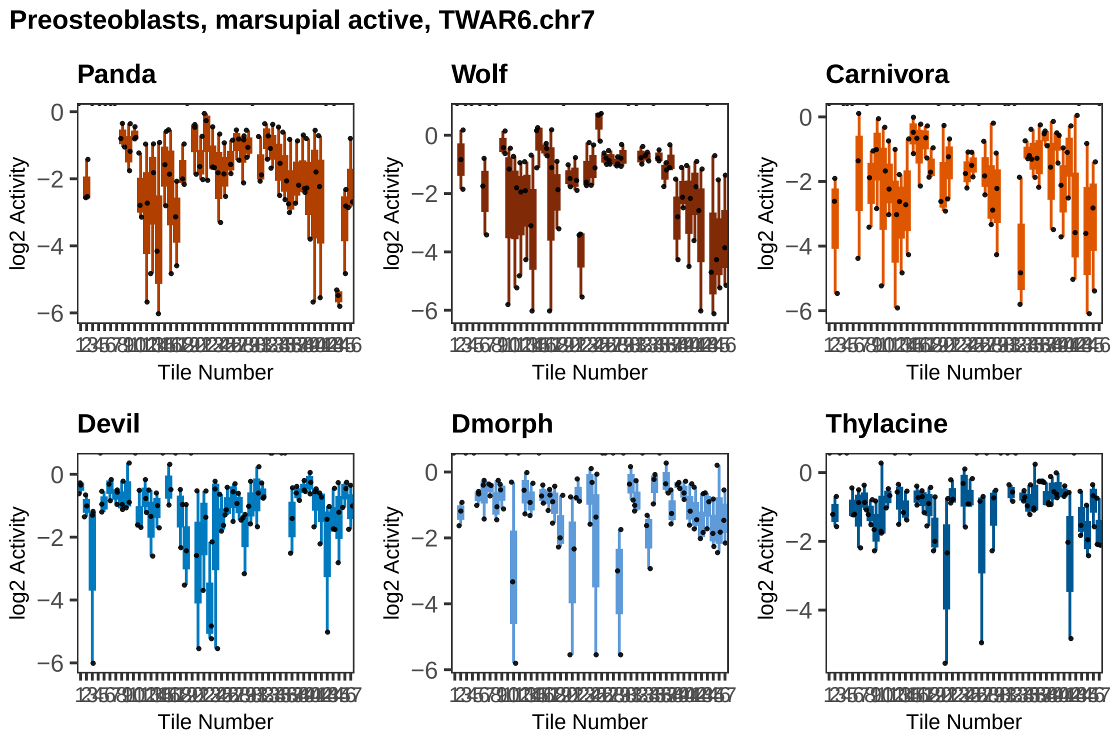

Preosteoblasts, marsupial active, TWAR6.chr7
Panda
Wolf
Carnivora
Devil
Dmorph
Thylacine
log2 Activity
Tile Number
0
−2
−4
−6

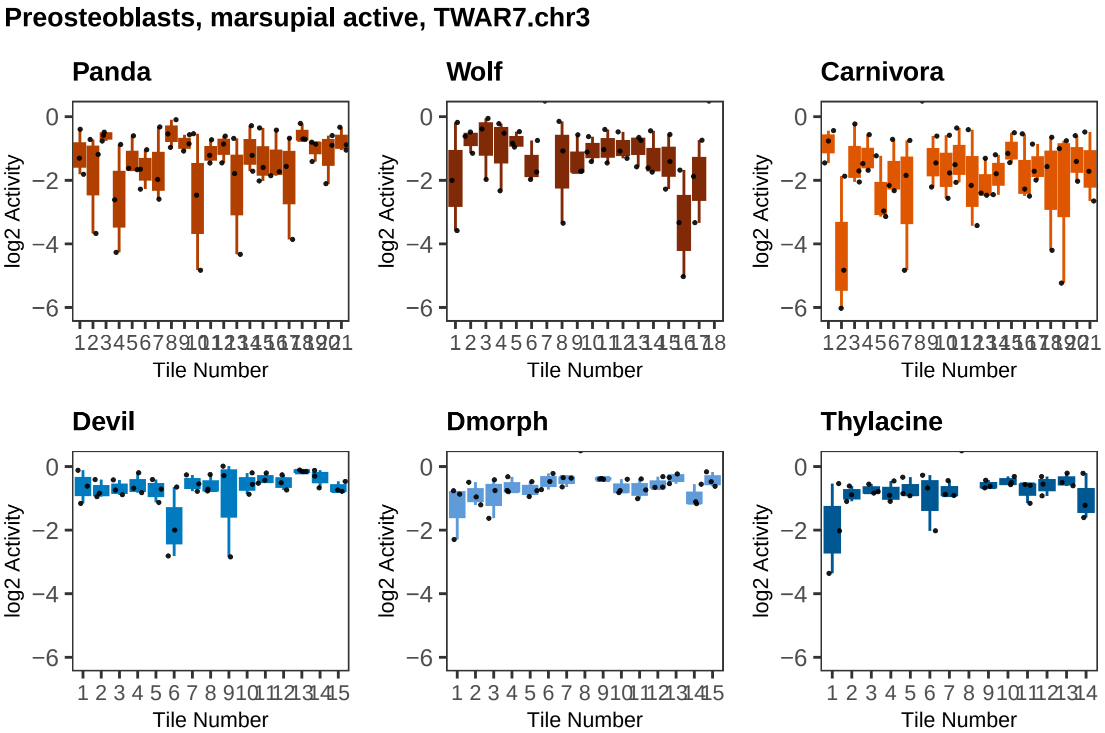
Preosteoblasts, marsupial active, TWAR7.chr3
Panda
Wolf
Carnivora
Devil
Dmorph
Thylacine
log2 Activity
0
−2
−4
−6
Tile Number

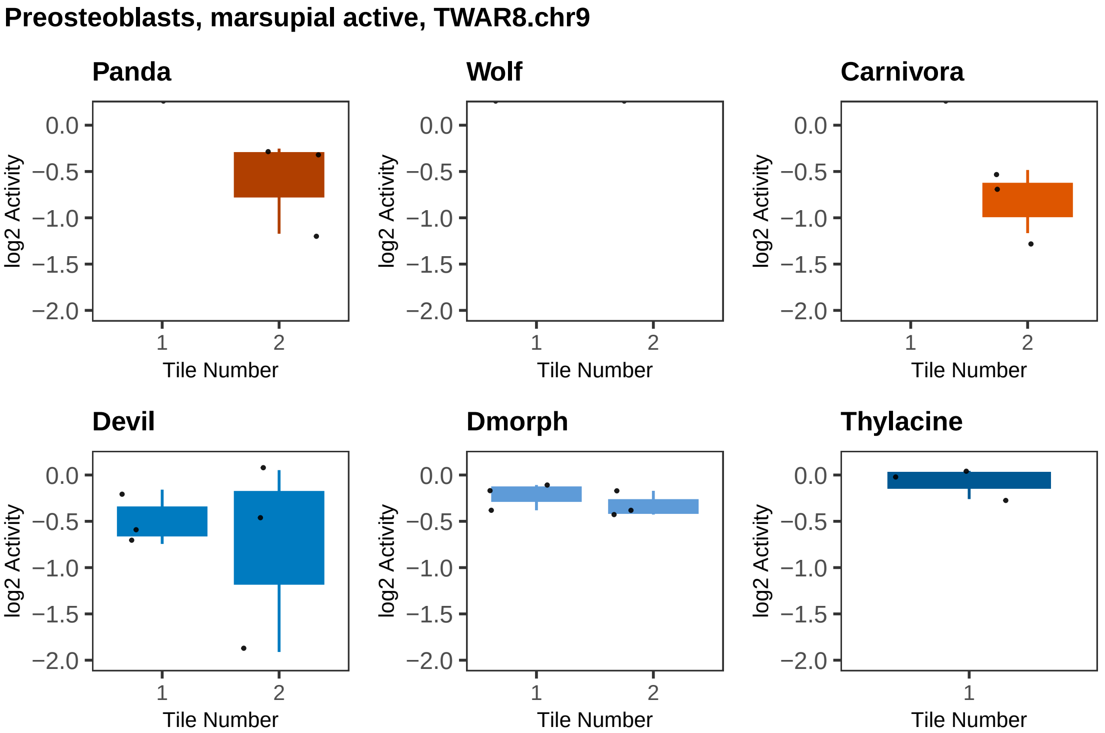

Preosteoblasts, marsupial active, TWAR8.chr9
Panda
Wolf
Carnivora
Devil
Dmorph
Thylacine
log2 Activity
0.0
−0.5
−1.0
−1.5
−2.0
1
2
Tile Number

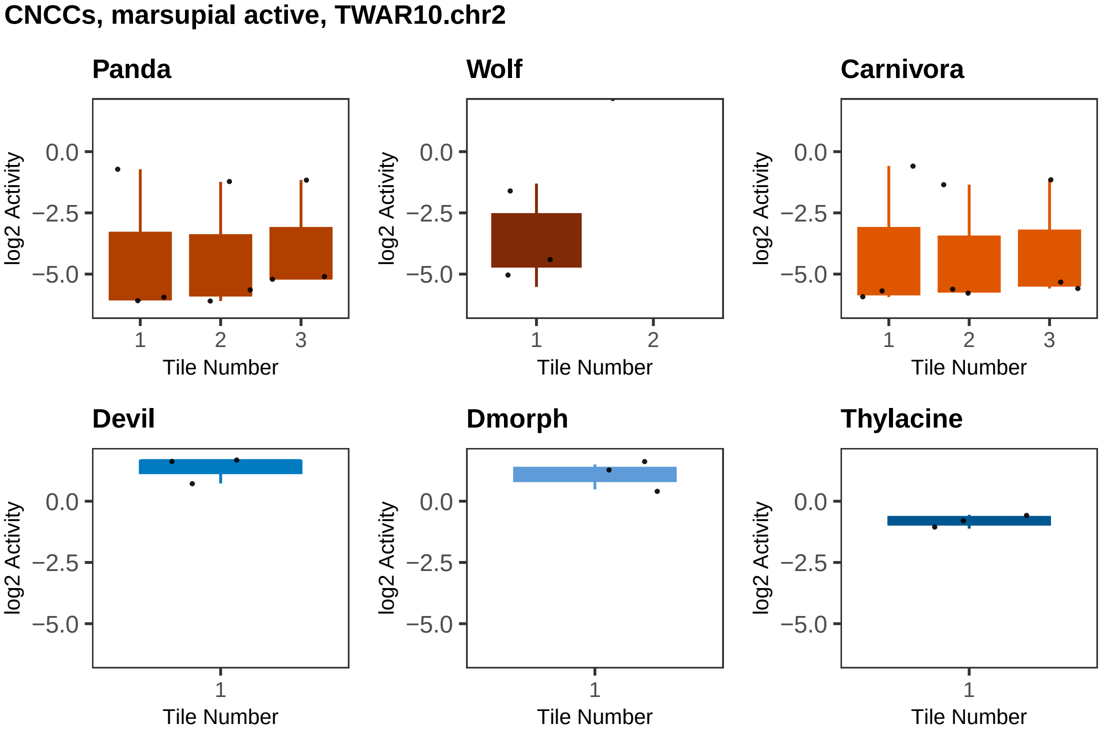

CNCCs, marsupial active, TWAR10.chr2
Panda
Wolf
Carnivora
Devil
Dmorph
Thylacine
log2 Activity
0.0
−2.5
−5.0
1
2
3
Tile Number

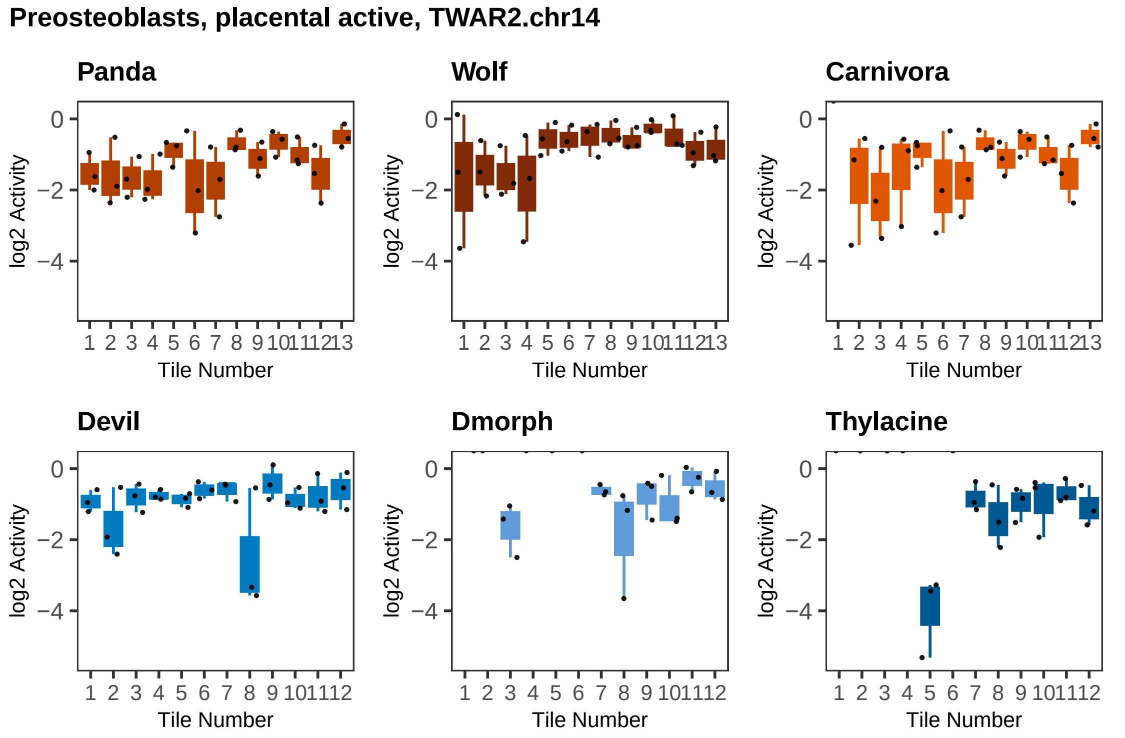

Preosteoblasts, placental active, TWAR2.chr14
Panda
Wolf
Carnivora
Devil
Dmorph
Thylacine
log2 Activity
0
−2
−4
Tile Number
1 2 3 4 5 6 7 8 9 10 11 12 13

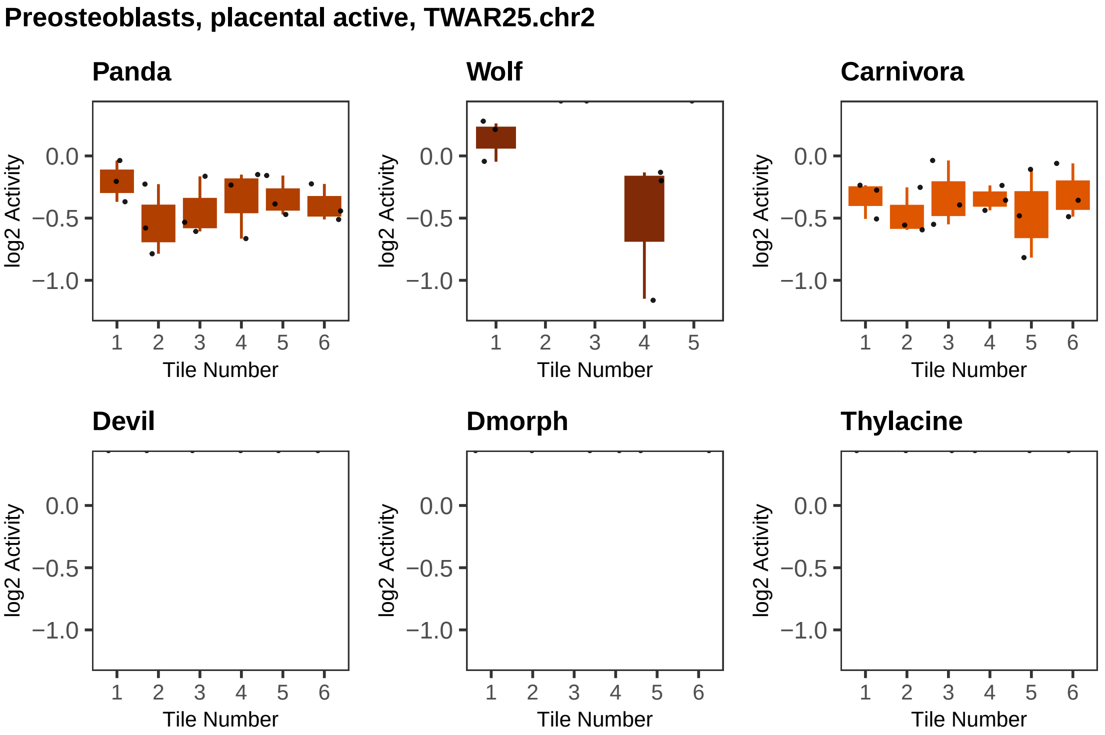

Preosteoblasts, placental active, TWAR25.chr2
Panda
Wolf
Carnivora
Devil
Dmorph
Thylacine
log2 Activity
0.0
−0.5
−1.0
Tile Number
1
2
3
4
5
6

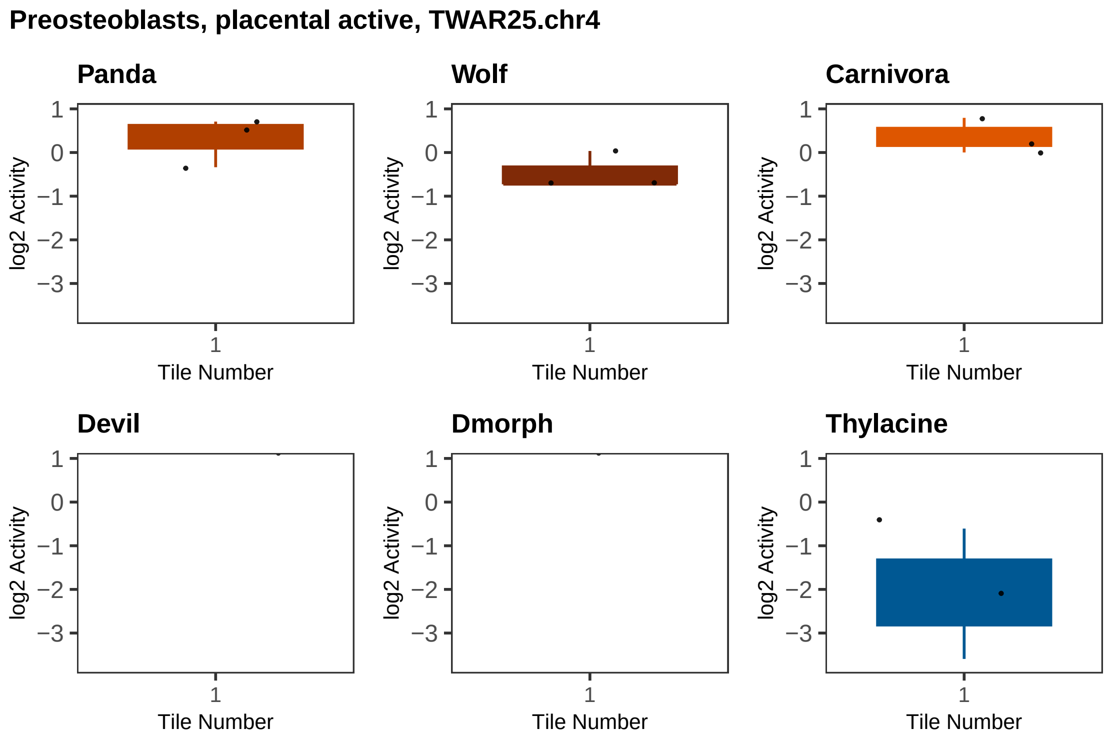
Preosteoblasts, placental active, TWAR25.chr4
Panda
Wolf
Carnivora
Devil
Dmorph
Thylacine
log2 Activity
1
0
−1
−2
−3
Tile Number
1

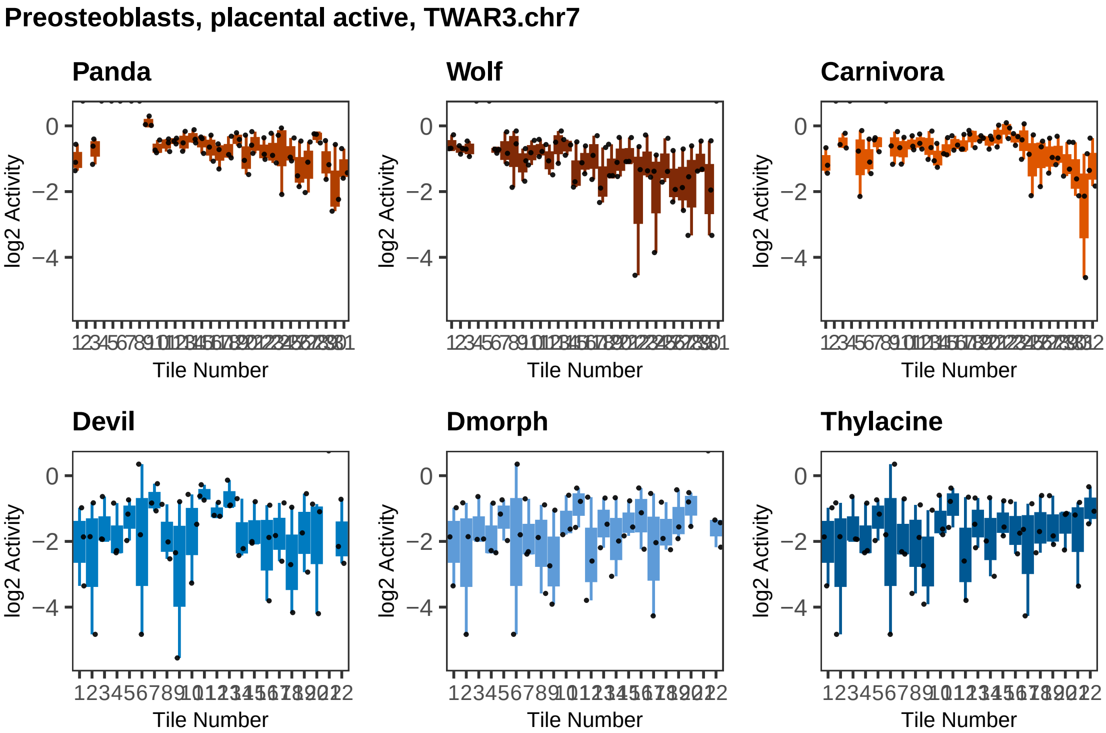
Preosteoblasts, placental active, TWAR3.chr7
Panda
Wolf
Carnivora
Devil
Dmorph
Thylacine
log2 Activity
0
−2
−4
Tile Number

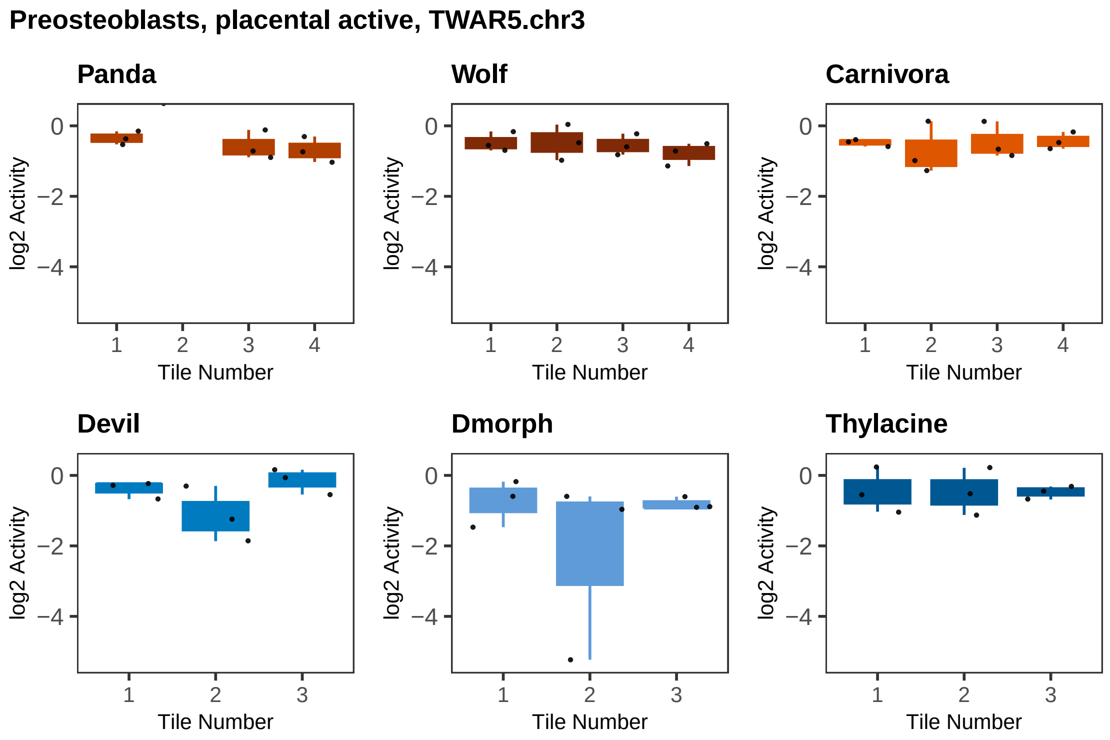

Preosteoblasts, placental active, TWAR5.chr3
Panda
Wolf
Carnivora
Devil
Dmorph
Thylacine
log2 Activity
Tile Number

### Preosteoblasts, placental active, TWAR7.chr17

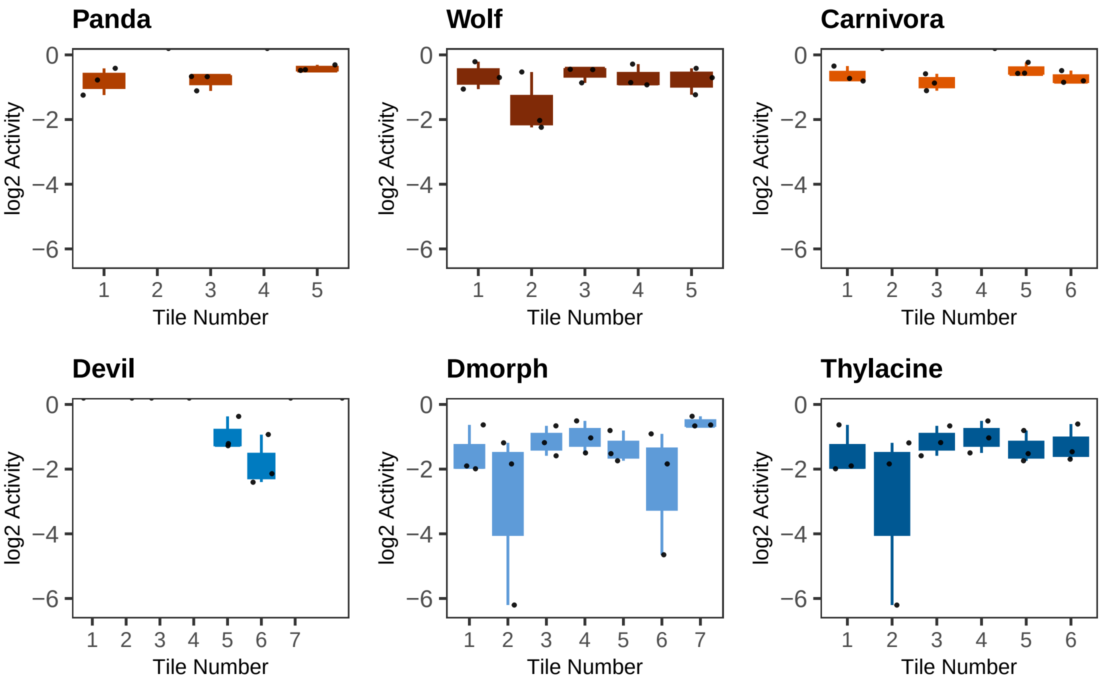

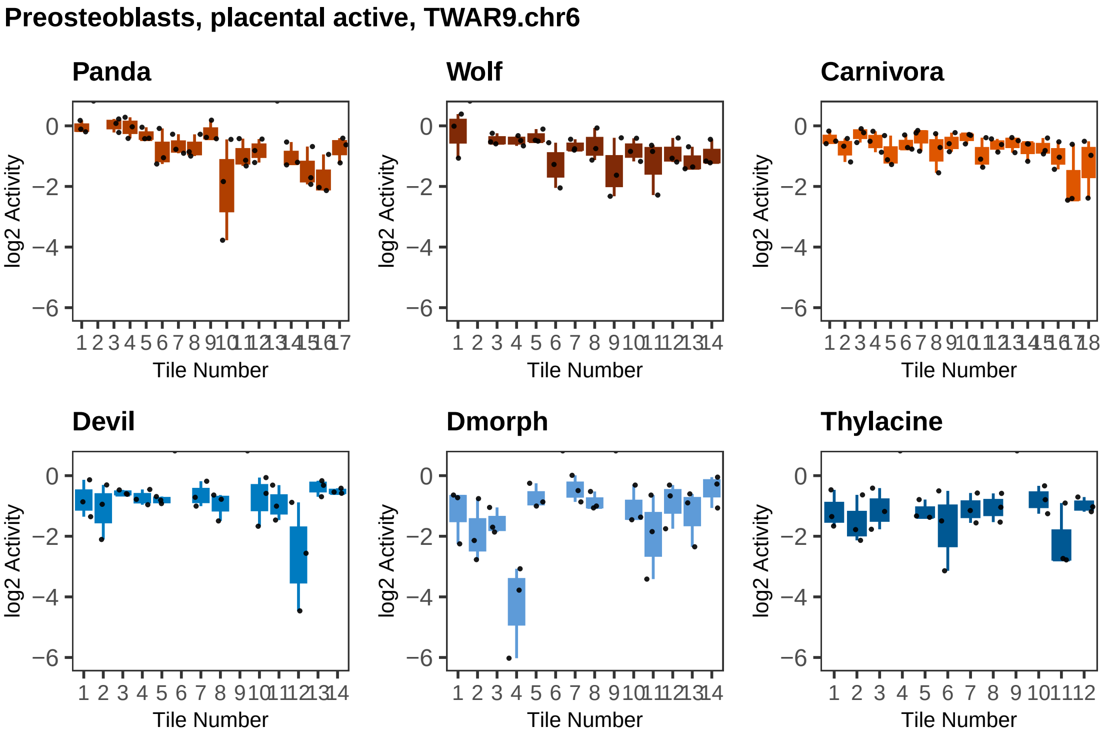

Preosteoblasts, placental active, TWAR9.chr6
Panda
Wolf
Carnivora
Devil
Dmorph
Thylacine
log2 Activity
0
−2
−4
−6
Tile Number

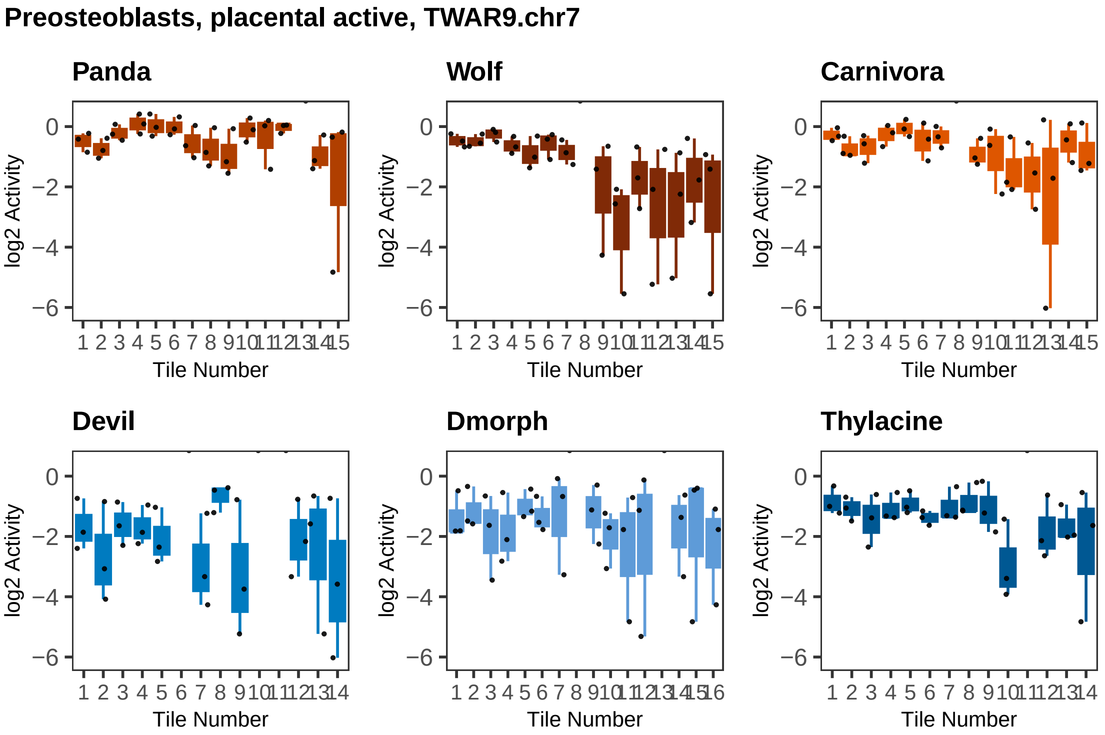

Preosteoblasts, placental active, TWAR9.chr7
Panda
Wolf
Carnivora
Devil
Dmorph
Thylacine
log2 Activity
Tile Number

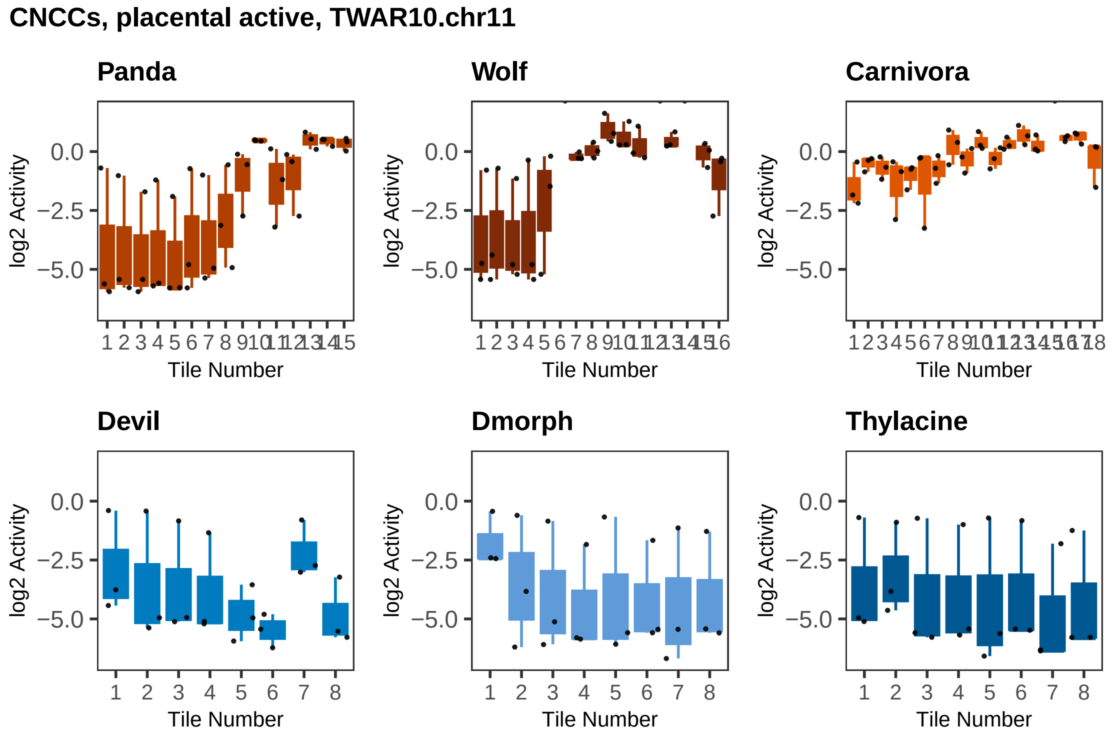

CNCCs, placental active, TWAR10.chr11
Panda
Wolf
Carnivora
Devil
Dmorph
Thylacine
log2 Activity
0.0
−2.5
−5.0
Tile Number

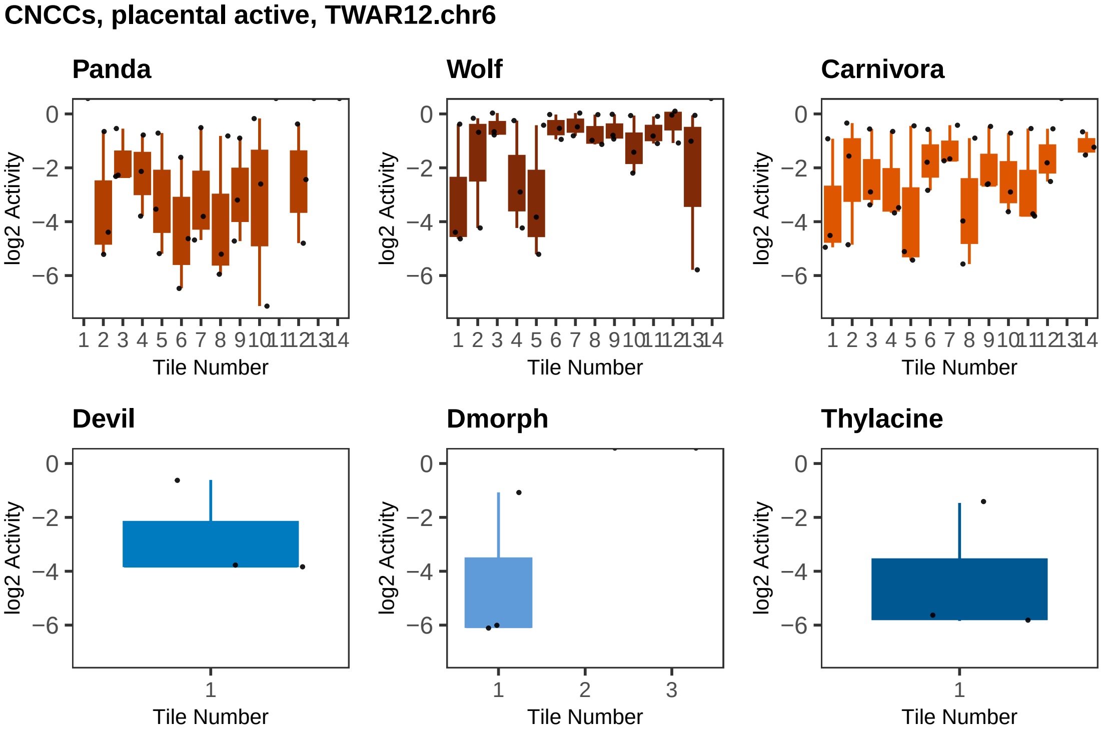

CNCCs, placental active, TWAR12.chr6
Panda
Wolf
Carnivora
Devil
Dmorph
Thylacine
log2 Activity
Tile Number

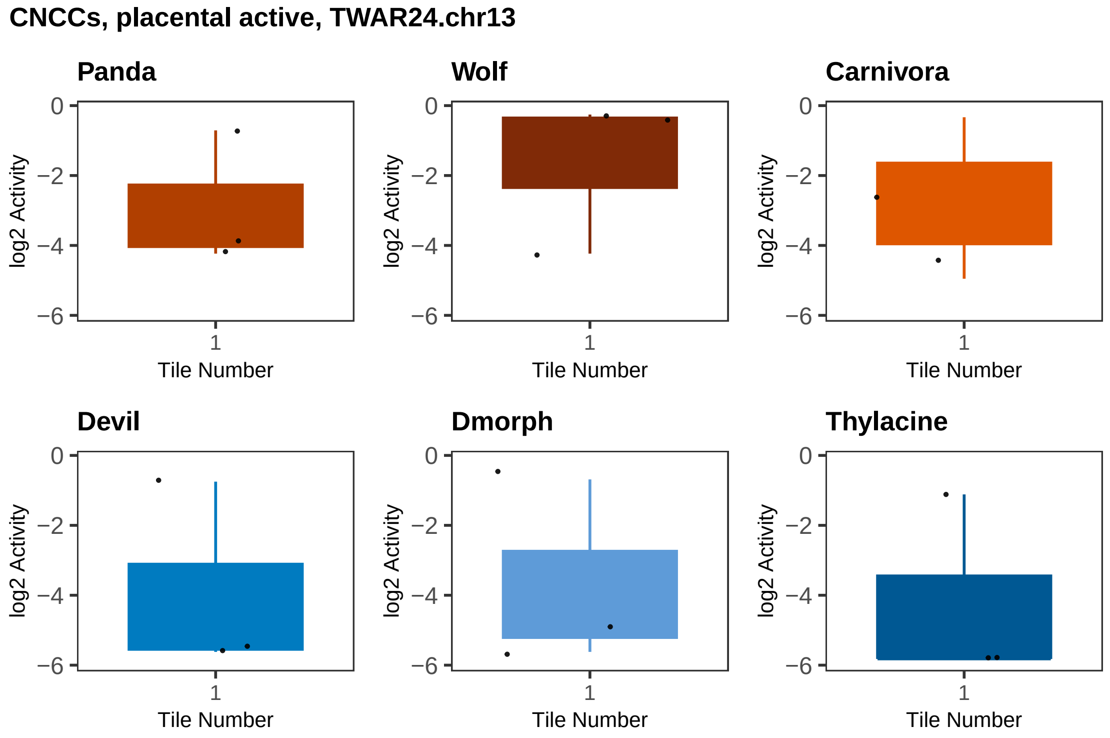
CNCCs, placental active, TWAR24.chr13
Panda
Wolf
Carnivora
Devil
Dmorph
Thylacine
log2 Activity
Tile Number
0
−2
−4
−6
1

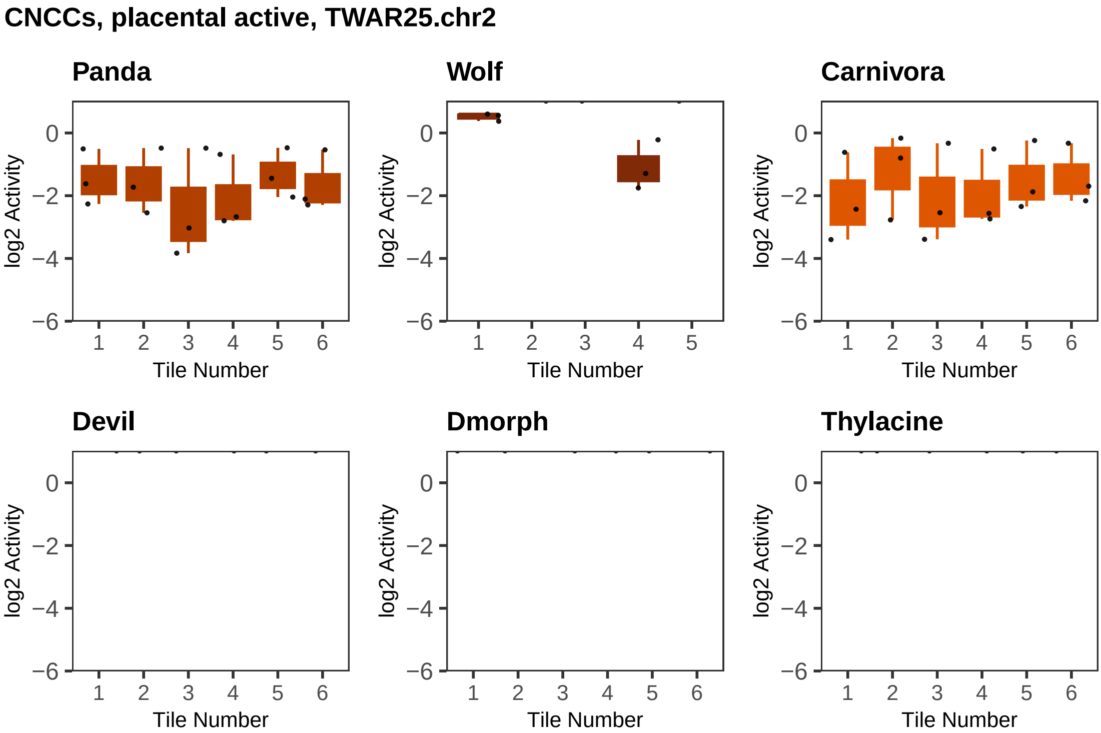

CNCCs, placental active, TWAR25.chr2
Panda
Wolf
Carnivora
Devil
Dmorph
Thylacine
log2 Activity
Tile Number

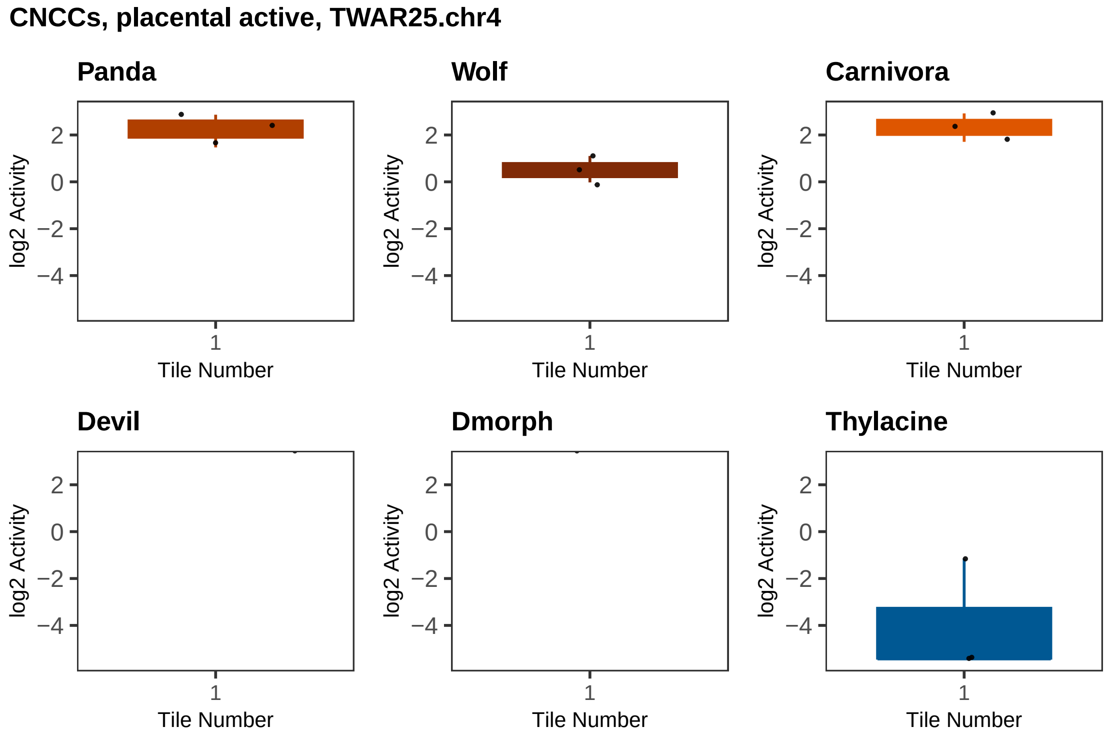
CNCCs, placental active, TWAR25.chr4
Panda
Wolf
Carnivora
Devil
Dmorph
Thylacine
log2 Activity
2
0
−2
−4
1
Tile Number

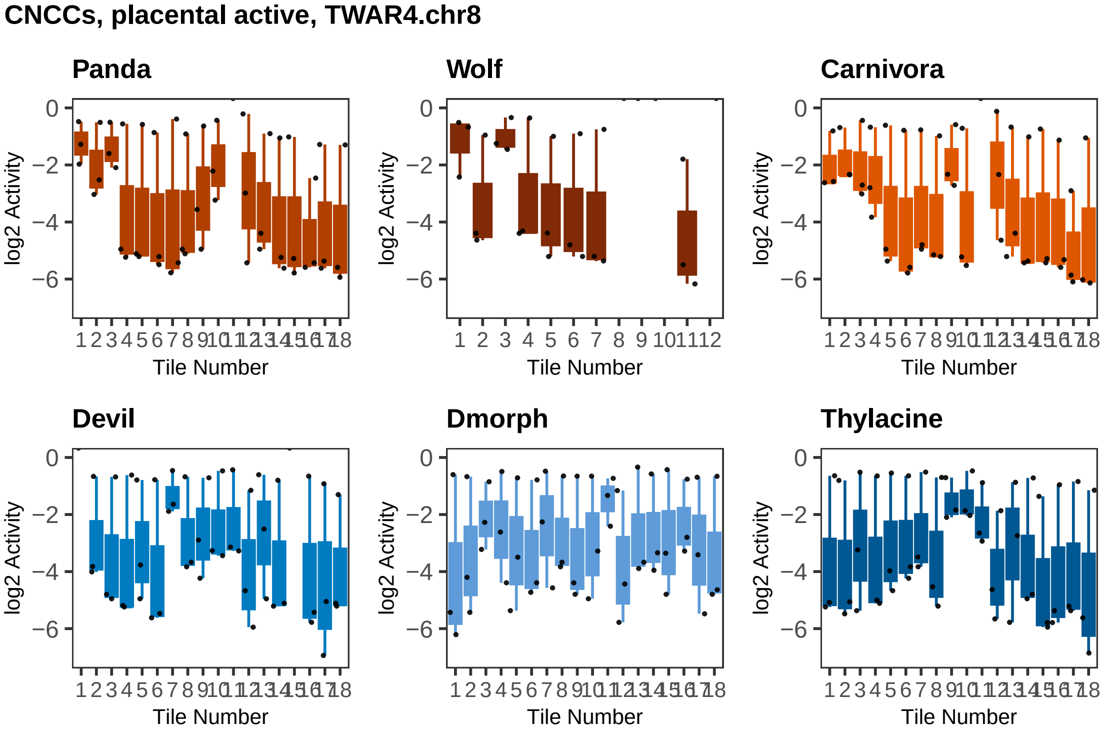

CNCCs, placental active, TWAR4.chr8
Panda
Wolf
Carnivora
Devil
Dmorph
Thylacine
log2 Activity
Tile Number
0
−2
−4
−6
1 2 3 4 5 6 7 8 9 10 11 12 13 14 15 16 17 18
1 2 3 4 5 6 7 8 9 10 11 12

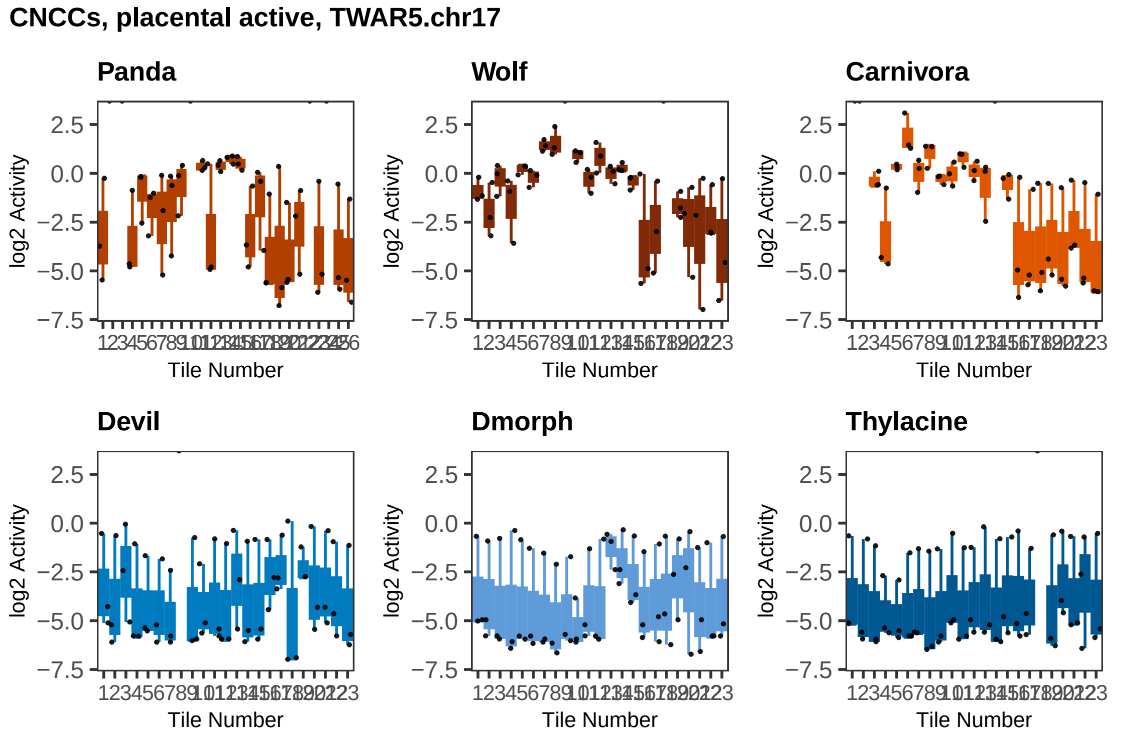

CNCCs, placental active, TWAR5.chr17
Panda
Wolf
Carnivora
Devil
Dmorph
Thylacine
log2 Activity
Tile Number
2.5
0.0
−2.5
−5.0
−7.5

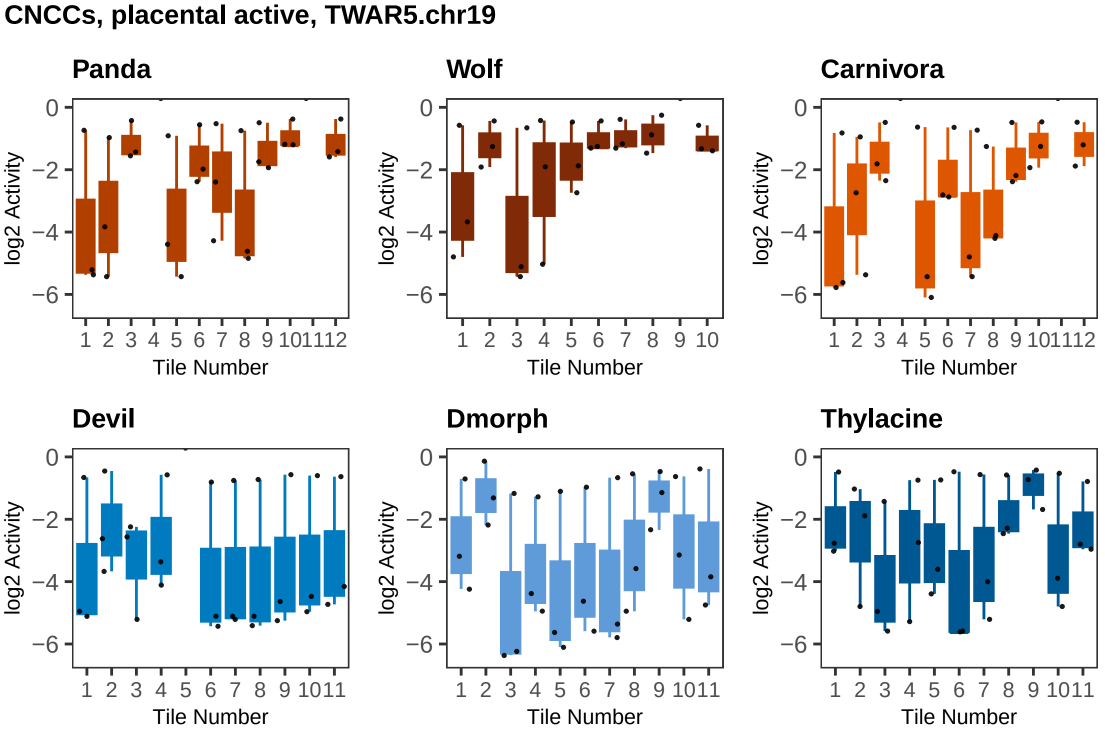

CNCCs, placental active, TWAR5.chr19
Panda
Wolf
Carnivora
Devil
Dmorph
Thylacine
log2 Activity
0
−2
−4
−6
1 2 3 4 5 6 7 8 9 10 11 12
1 2 3 4 5 6 7 8 9 10
1 2 3 4 5 6 7 8 9 10 11
Tile Number

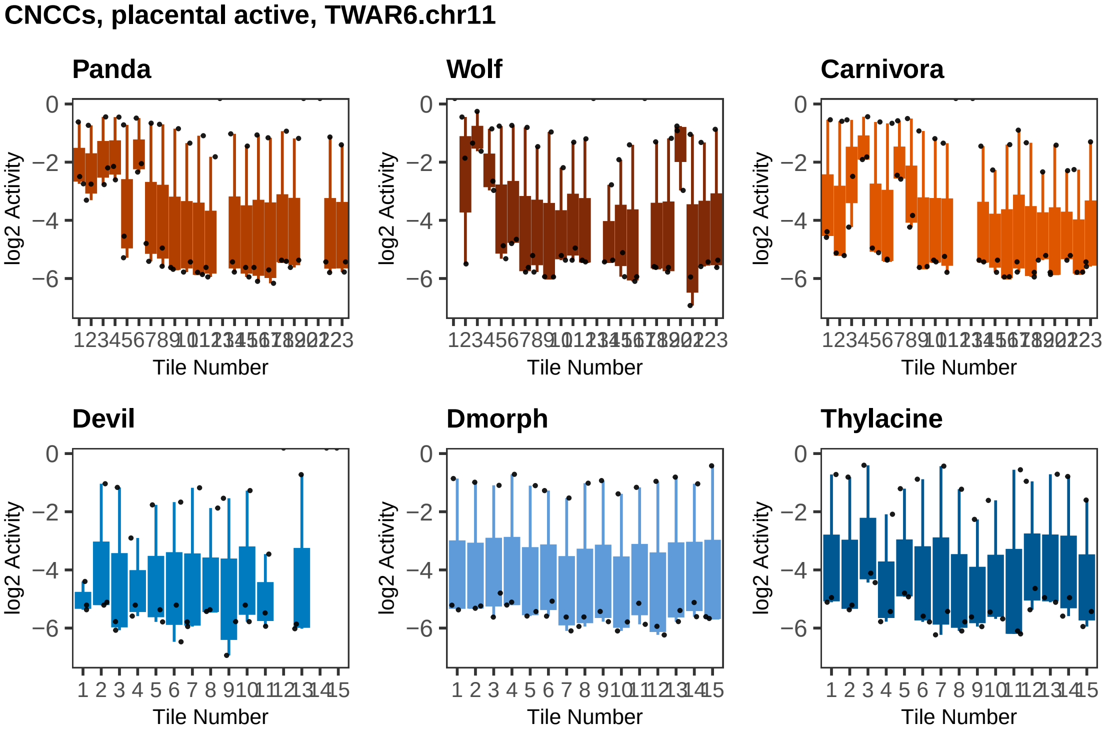
CNCCs, placental active, TWAR6.chr11
Panda
Wolf
Carnivora
Devil
Dmorph
Thylacine
log2 Activity
0
−2
−4
−6
Tile Number
1 2 3 4 5 6 7 8 9 10 11 12 13 14 15

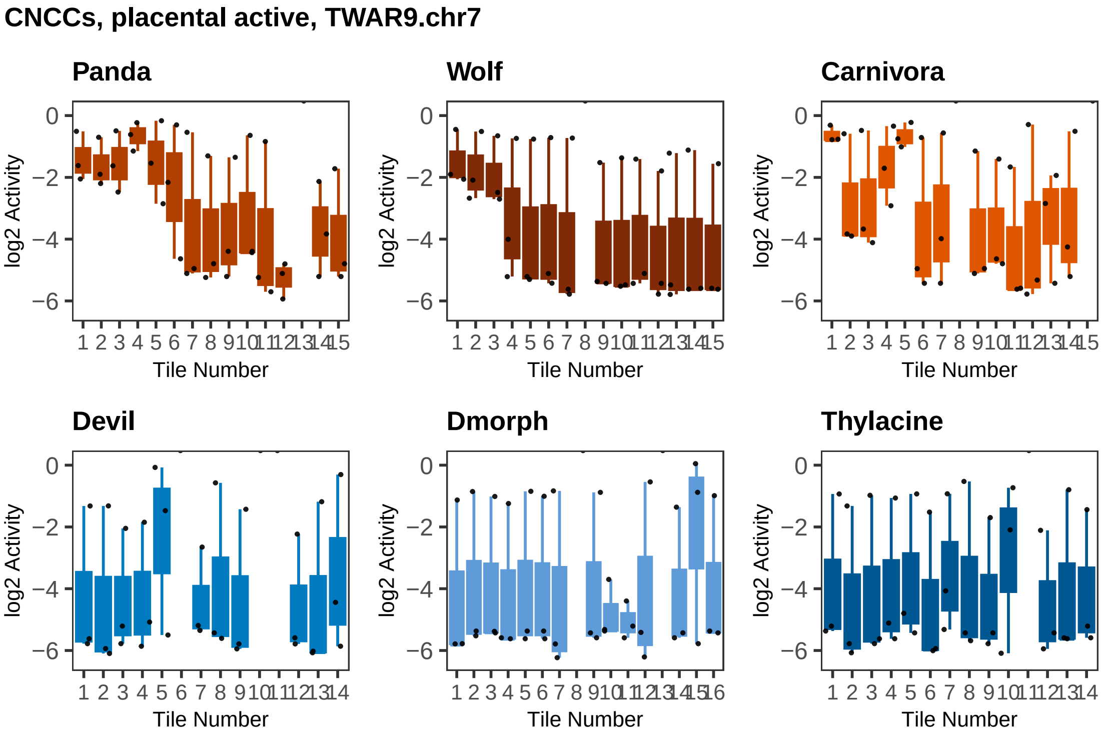
CNCCs, placental active, TWAR9.chr7
Panda
Wolf
Carnivora
Devil
Dmorph
Thylacine
log2 Activity
0
−2
−4
−6
Tile Number
1 2 3 4 5 6 7 8 9 10 11 12 13 14 15
1 2 3 4 5 6 7 8 9 10 11 12 13 14
1 2 3 4 5 6 7 8 9 10 11 12 13 14 15 16
