## Supplementary File 1 for "Dissecting functional regulatory convergence over 160 million years of therian evolution"

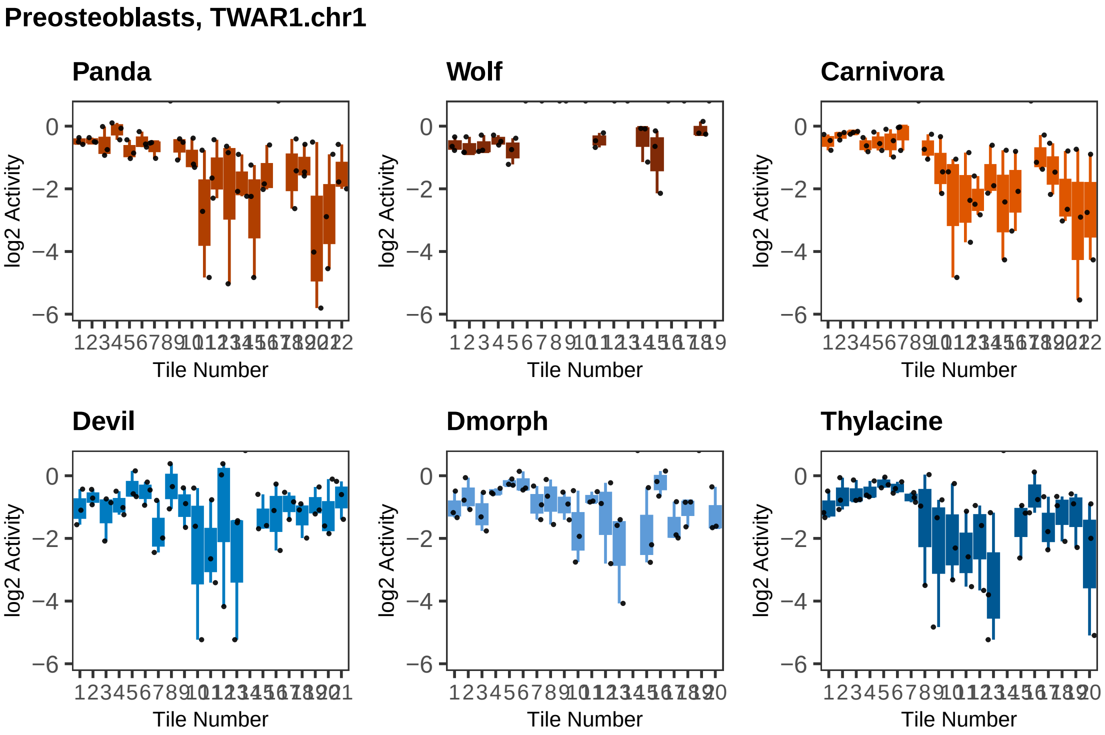
Preosteoblasts, TWAR1.chr1
Panda
Wolf
Carnivora
Devil
Dmorph
Thylacine
log2 Activity
0
−2
−4
−6
Tile Number

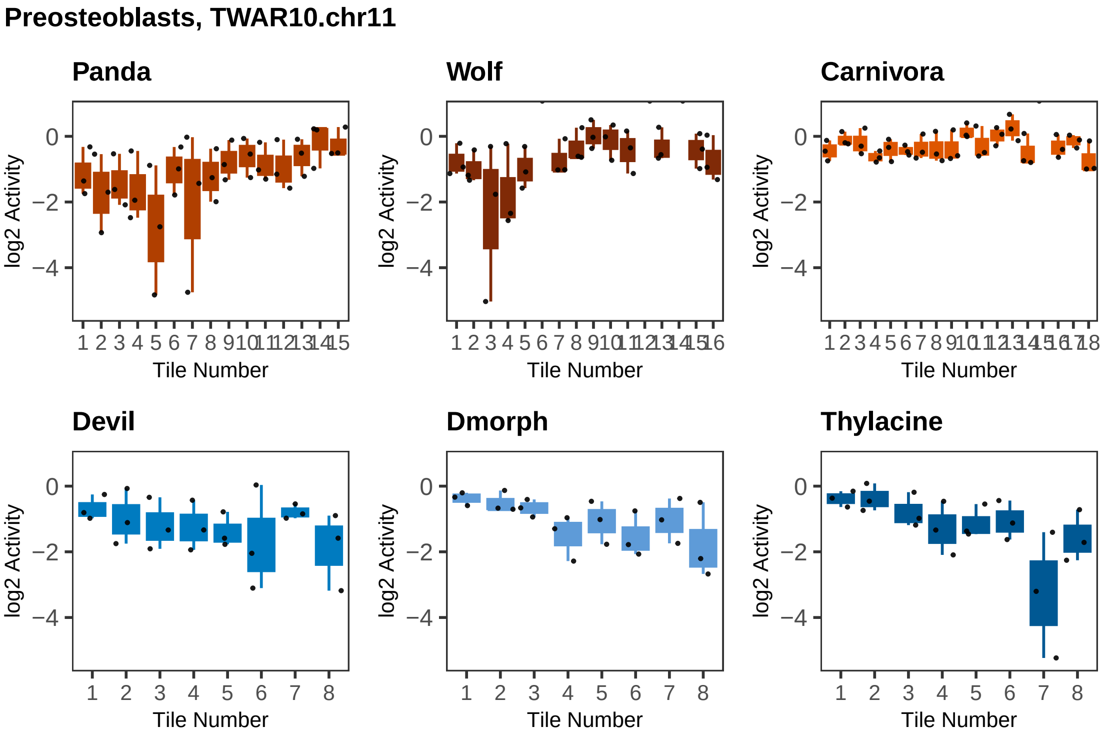
Preosteoblasts, TWAR10.chr11
Panda
Wolf
Carnivora
Devil
Dmorph
Thylacine
log2 Activity
Tile Number
0
−2
−4

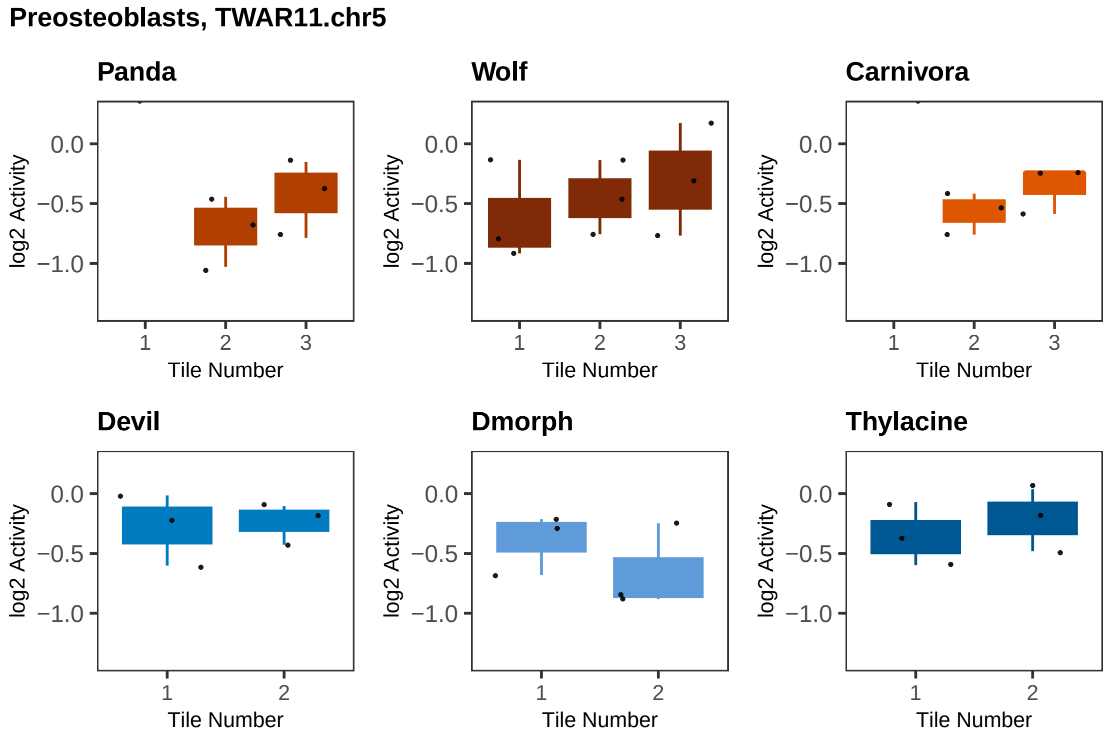

Preosteoblasts, TWAR11.chr5
Panda
Wolf
Carnivora
Devil
Dmorph
Thylacine
log2 Activity
Tile Number
0.0
−0.5
−1.0
1
2
3

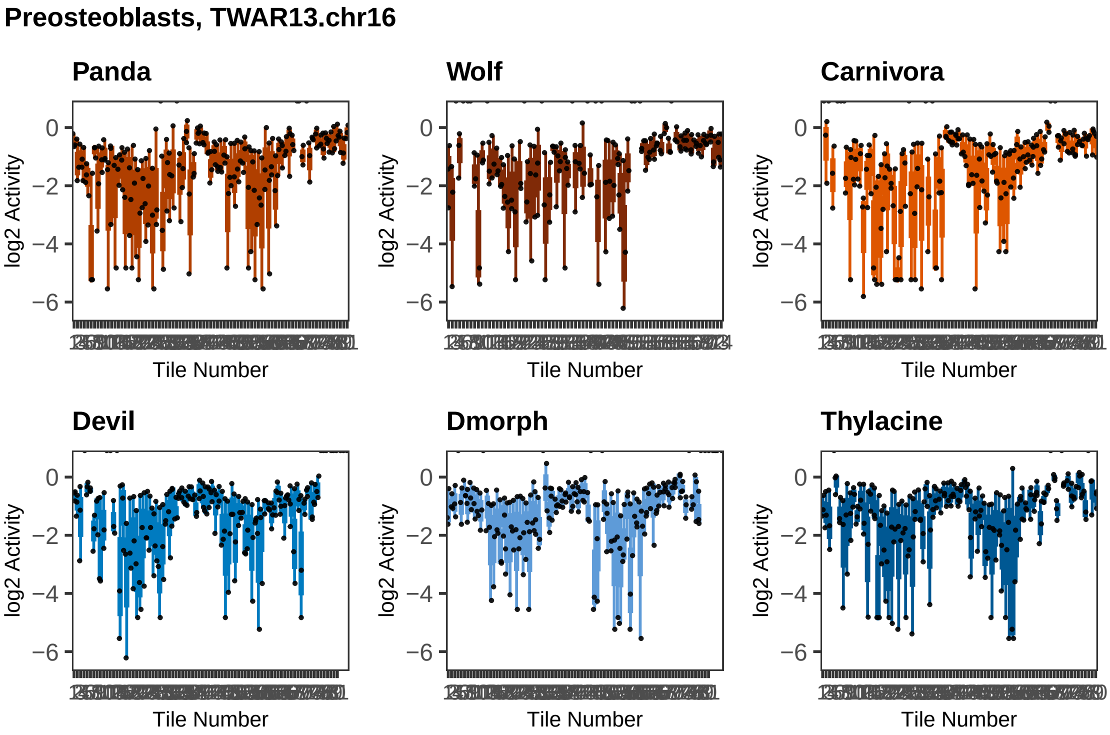

Preosteoblasts, TWAR13.chr16
Panda
Wolf
Carnivora
Devil
Dmorph
Thylacine
log2 Activity
Tile Number
0
−2
−4
−6

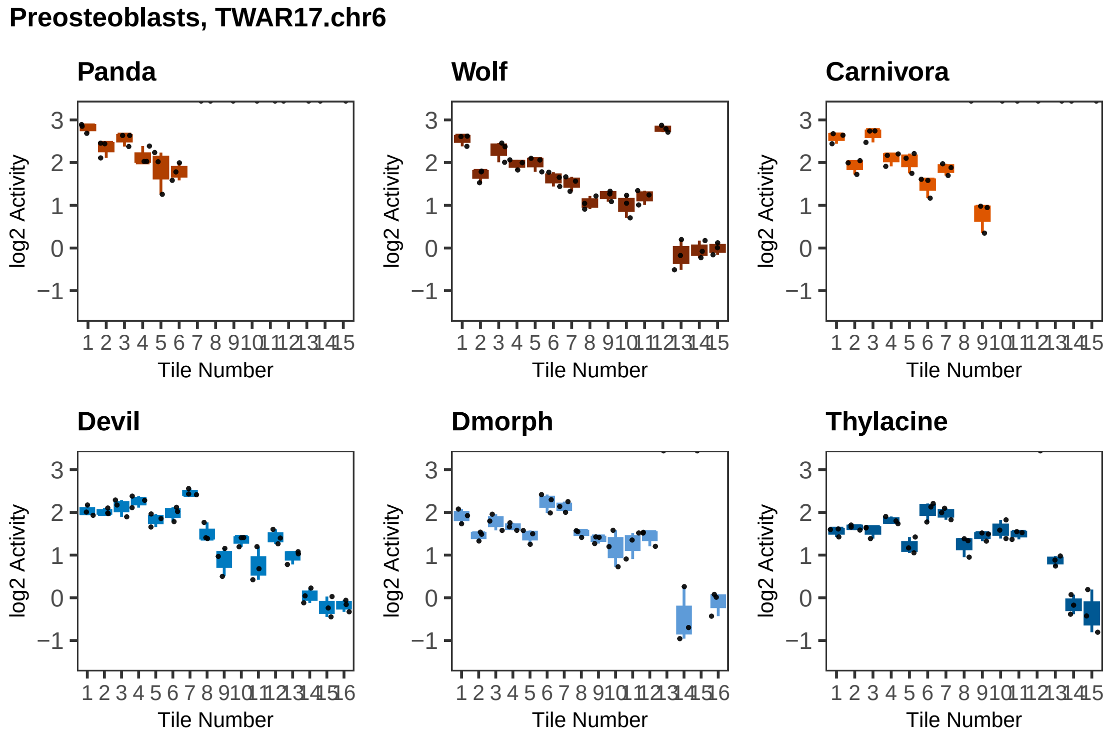

Preosteoblasts, TWAR17.chr6
Panda
Wolf
Carnivora
Devil
Dmorph
Thylacine
log2 Activity
Tile Number

**Preosteoblasts, TWAR2.chr1**

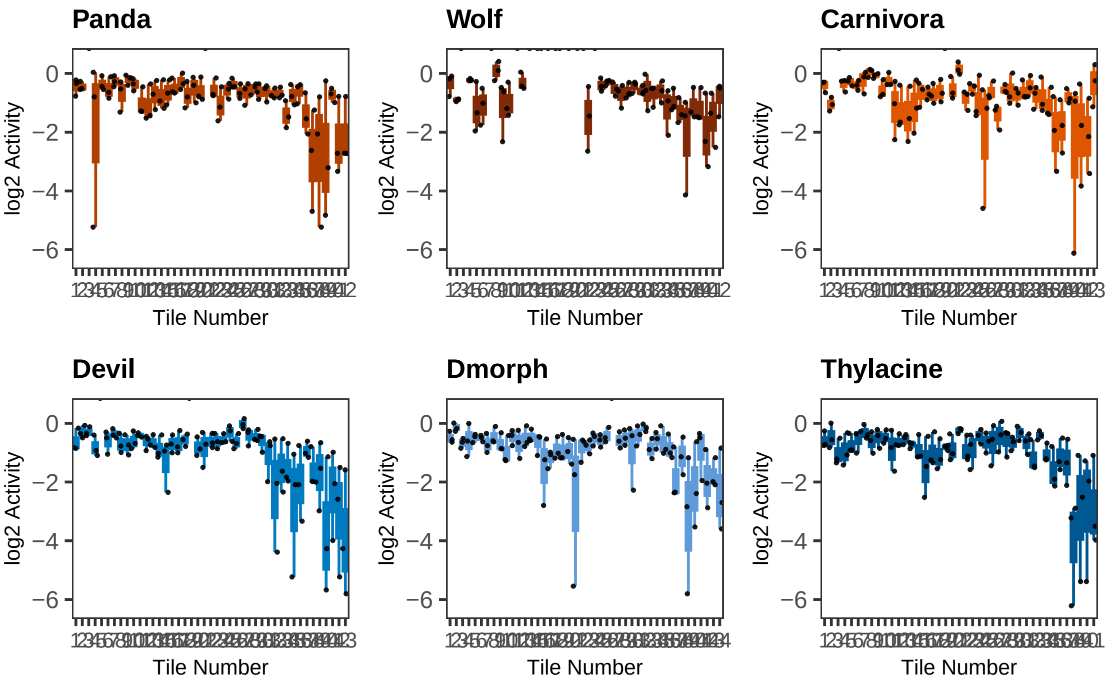

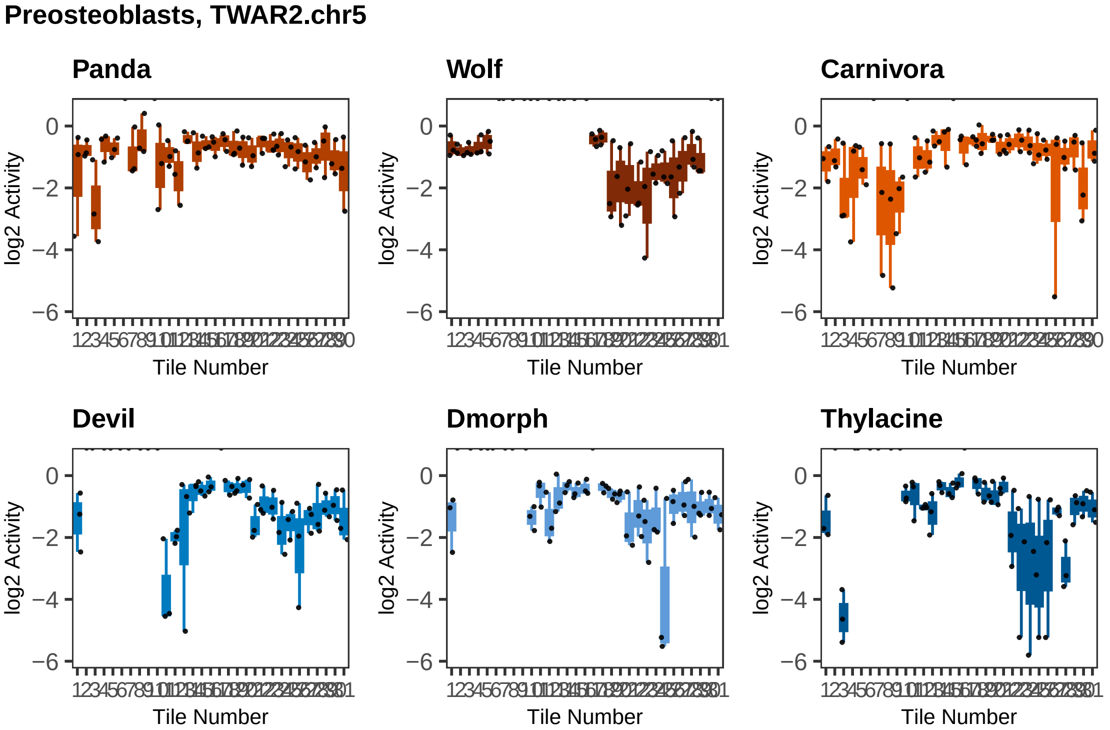
Preosteoblasts, TWAR2.chr5
Panda
Wolf
Carnivora
Devil
Dmorph
Thylacine
log2 Activity
Tile Number
0
−2
−4
−6

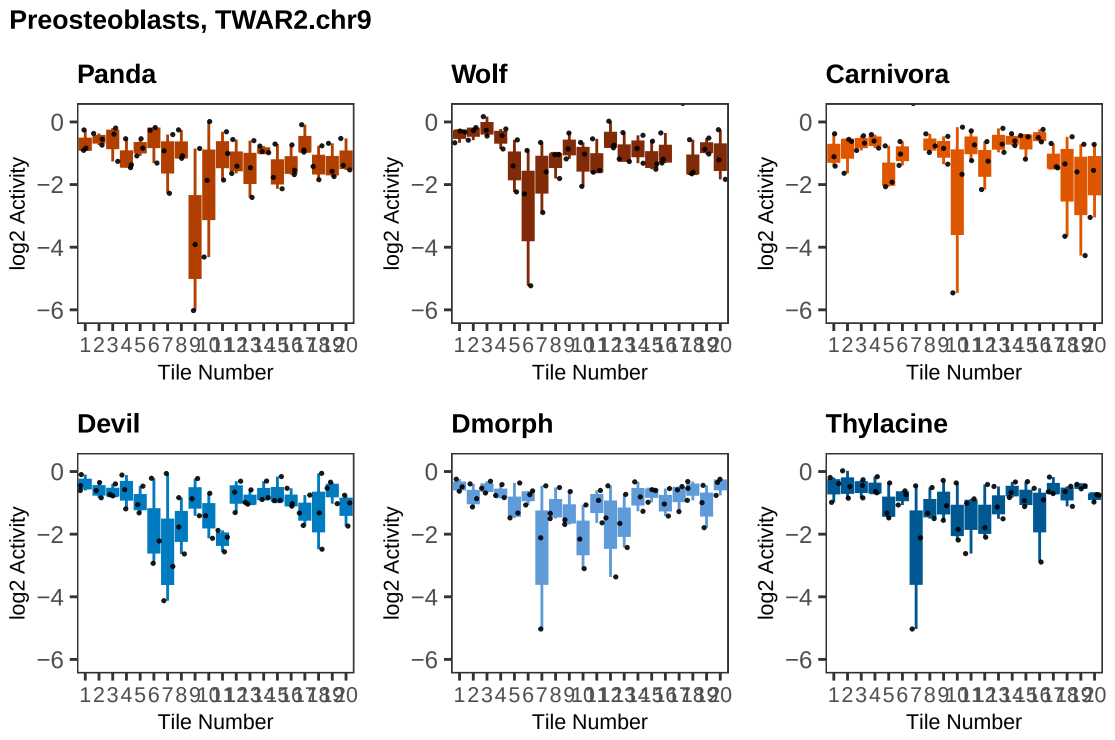
Preosteoblasts, TWAR2.chr9
Panda
Wolf
Carnivora
Devil
Dmorph
Thylacine
log2 Activity
0
−2
−4
−6
Tile Number
1 2 3 4 5 6 7 8 9 10 11 12 13 14 15 16 17 18 19 20

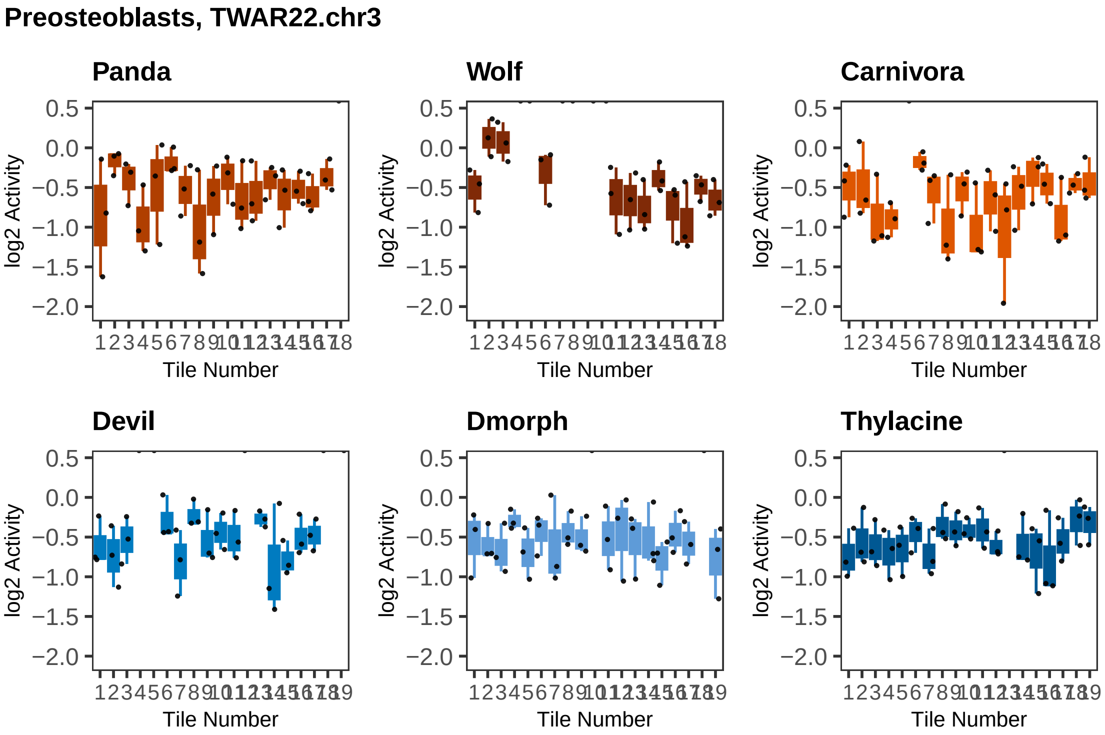
Preosteoblasts, TWAR22.chr3
Panda
Wolf
Carnivora
Devil
Dmorph
Thylacine
log2 Activity
Tile Number
0.5
0.0
−0.5
−1.0
−1.5
−2.0

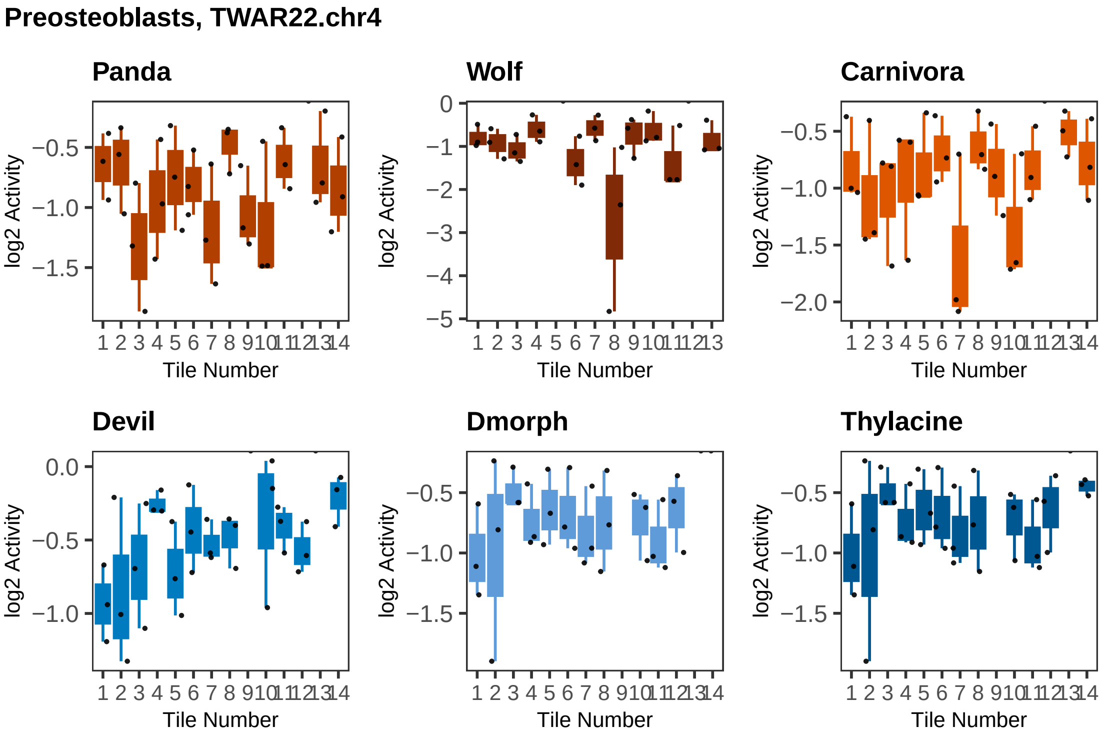

Preosteoblasts, TWAR22.chr4
Panda
Wolf
Carnivora
Devil
Dmorph
Thylacine
log2 Activity
Tile Number

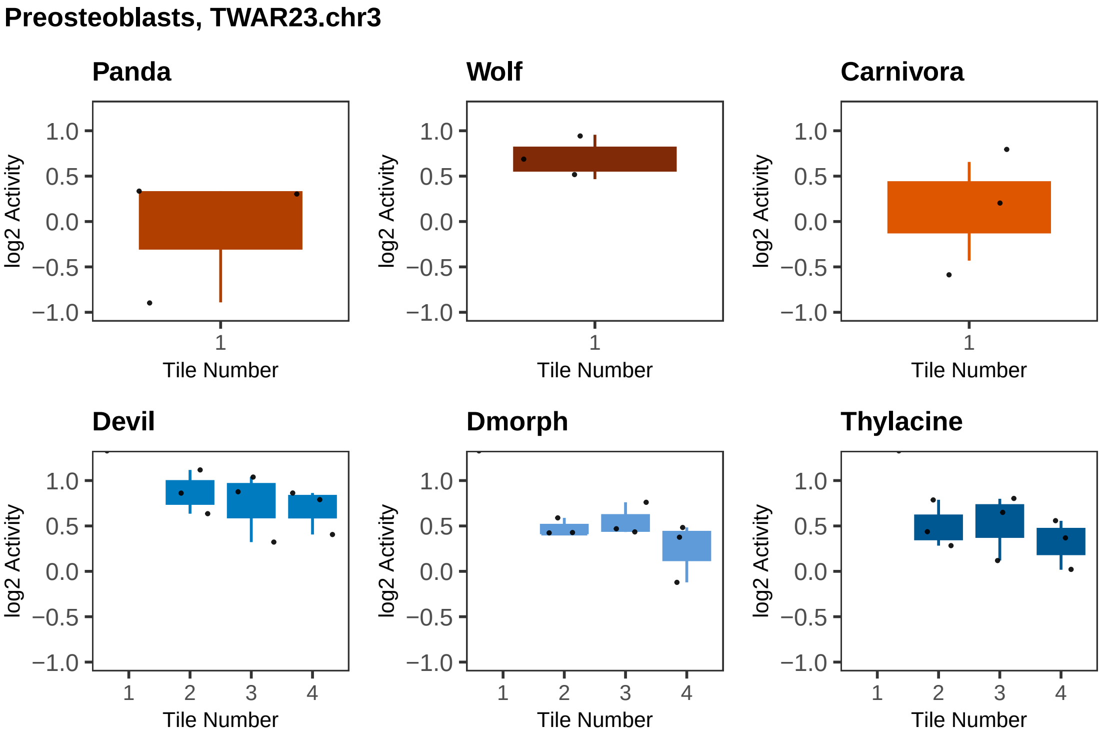
Preosteoblasts, TWAR23.chr3
Panda
Wolf
Carnivora
Devil
Dmorph
Thylacine
log2 Activity
1.0
0.5
0.0
−0.5
−1.0
Tile Number
1
2
3
4

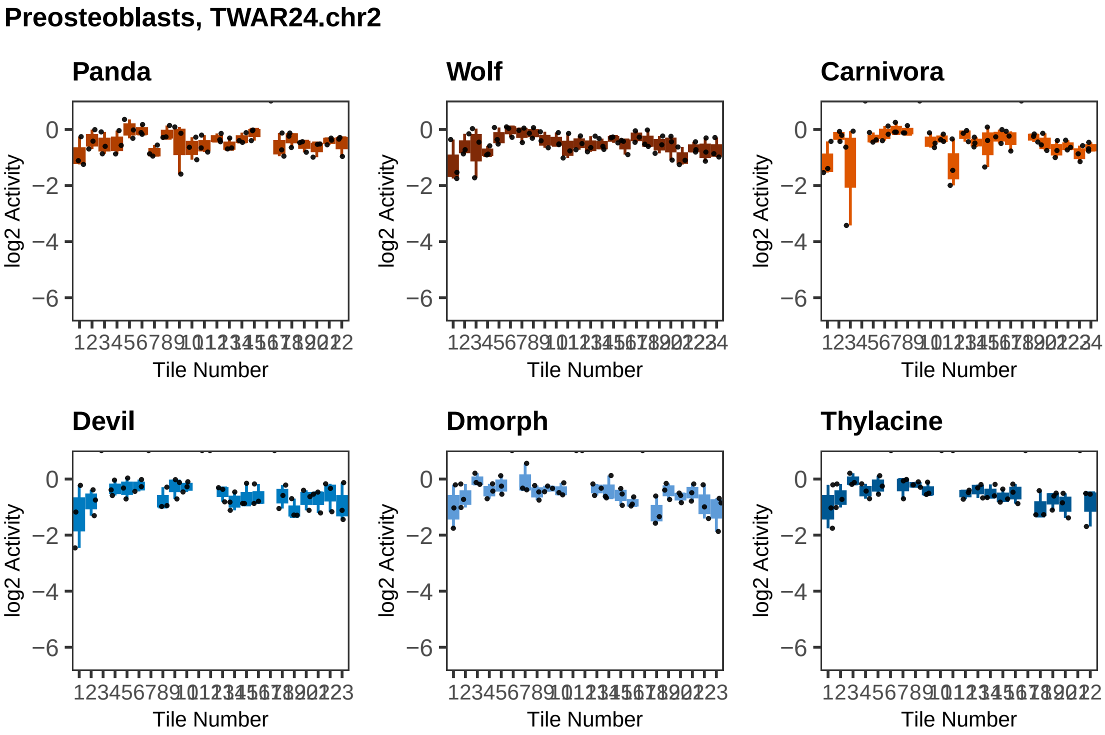
Preosteoblasts, TWAR24.chr2
Panda
Wolf
Carnivora
Devil
Dmorph
Thylacine
log2 Activity
0
−2
−4
−6
Tile Number

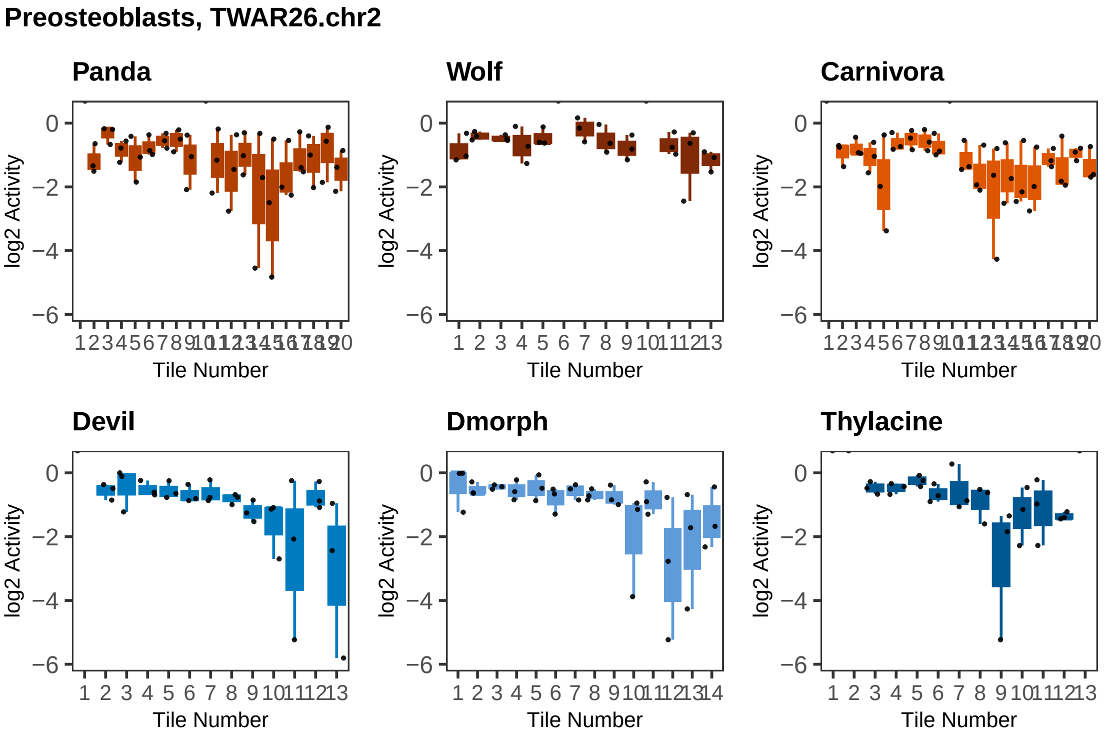

Preosteoblasts, TWAR26.chr2
Panda
Wolf
Carnivora
Devil
Dmorph
Thylacine
log2 Activity
Tile Number
0
−2
−4
−6

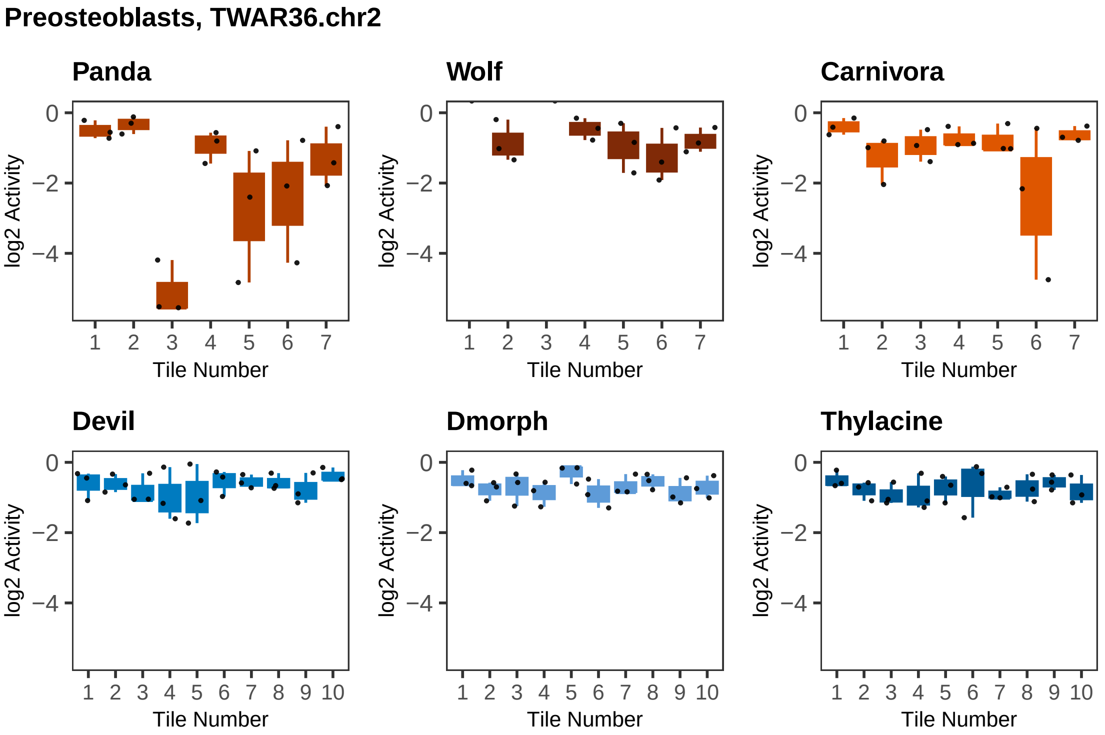

Preosteoblasts, TWAR36.chr2
Panda
Wolf
Carnivora
Devil
Dmorph
Thylacine
log2 Activity
Tile Number
0
−2
−4
1 2 3 4 5 6 7
1 2 3 4 5 6 7 8 9 10

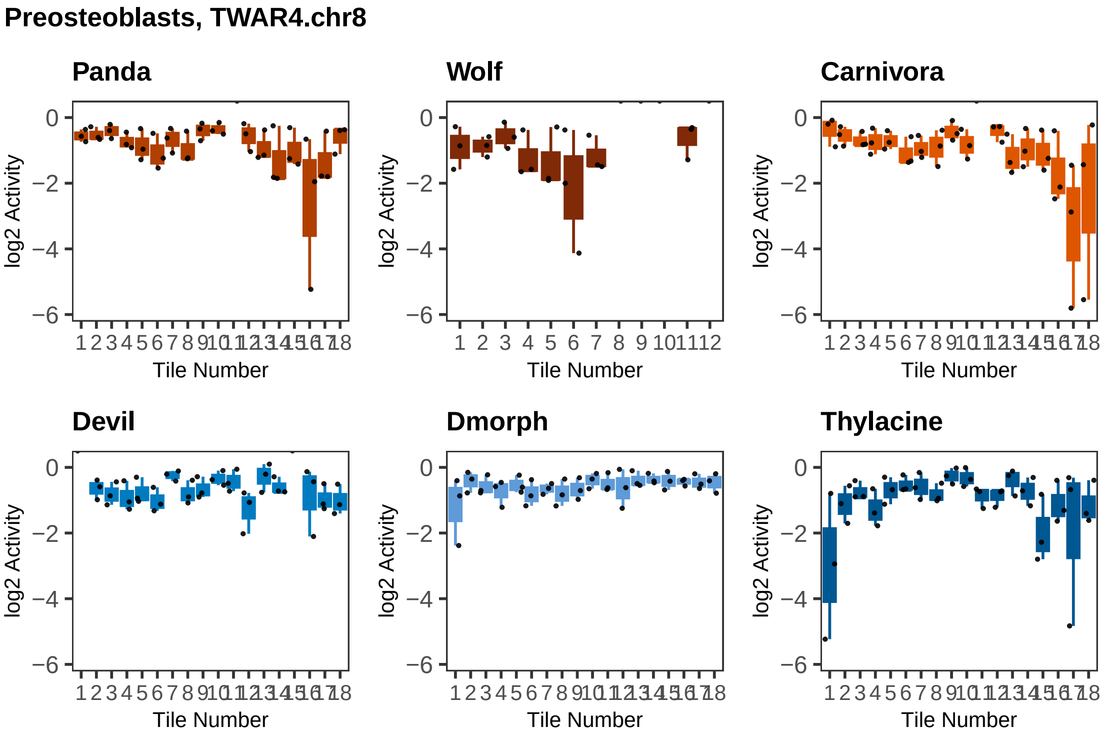

Preosteoblasts, TWAR4.chr8
Panda
Wolf
Carnivora
Devil
Dmorph
Thylacine
log2 Activity
Tile Number
0
−2
−4
−6

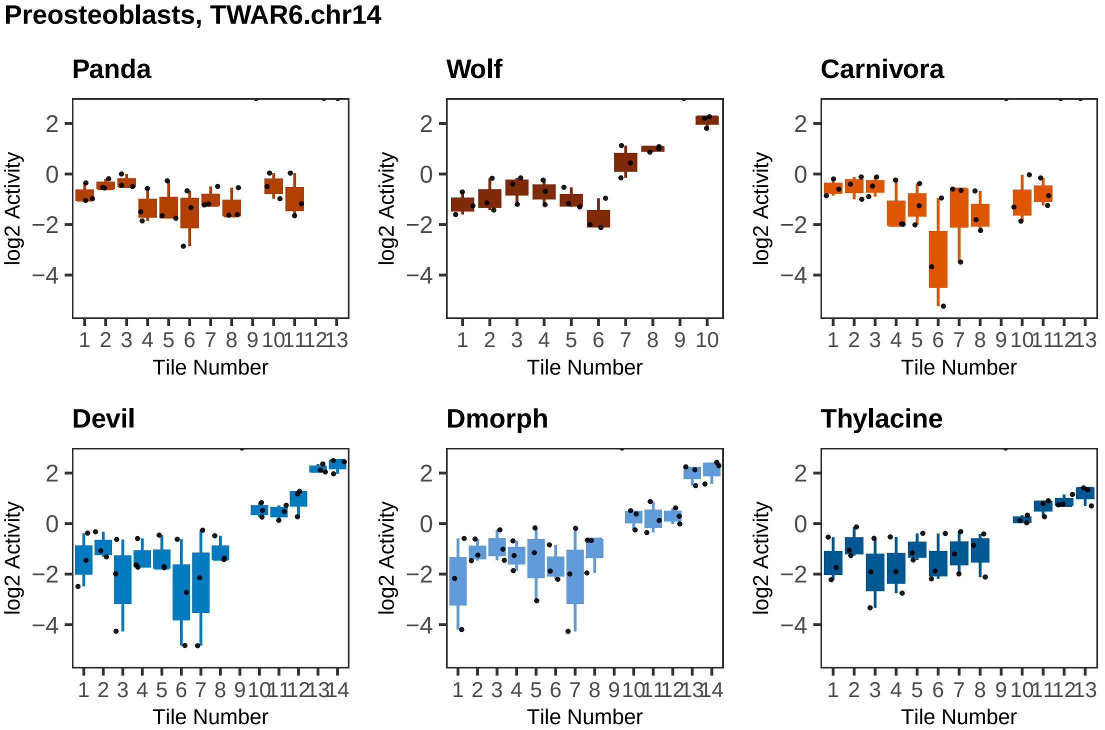
Preosteoblasts, TWAR6.chr14
Panda
Wolf
Carnivora
Devil
Dmorph
Thylacine
log2 Activity
Tile Number
2
0
−2
−4
1 2 3 4 5 6 7 8 9 10 11 12 13 14

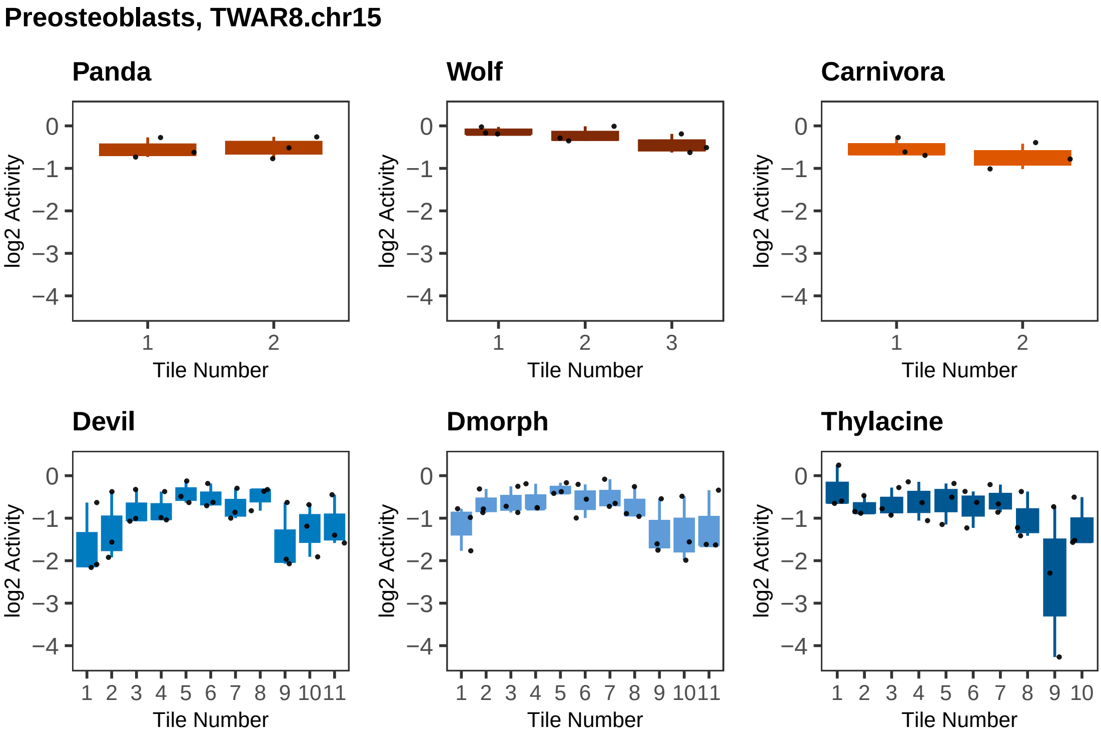

Preosteoblasts, TWAR8.chr15
Panda
Wolf
Carnivora
Devil
Dmorph
Thylacine
log2 Activity
Tile Number

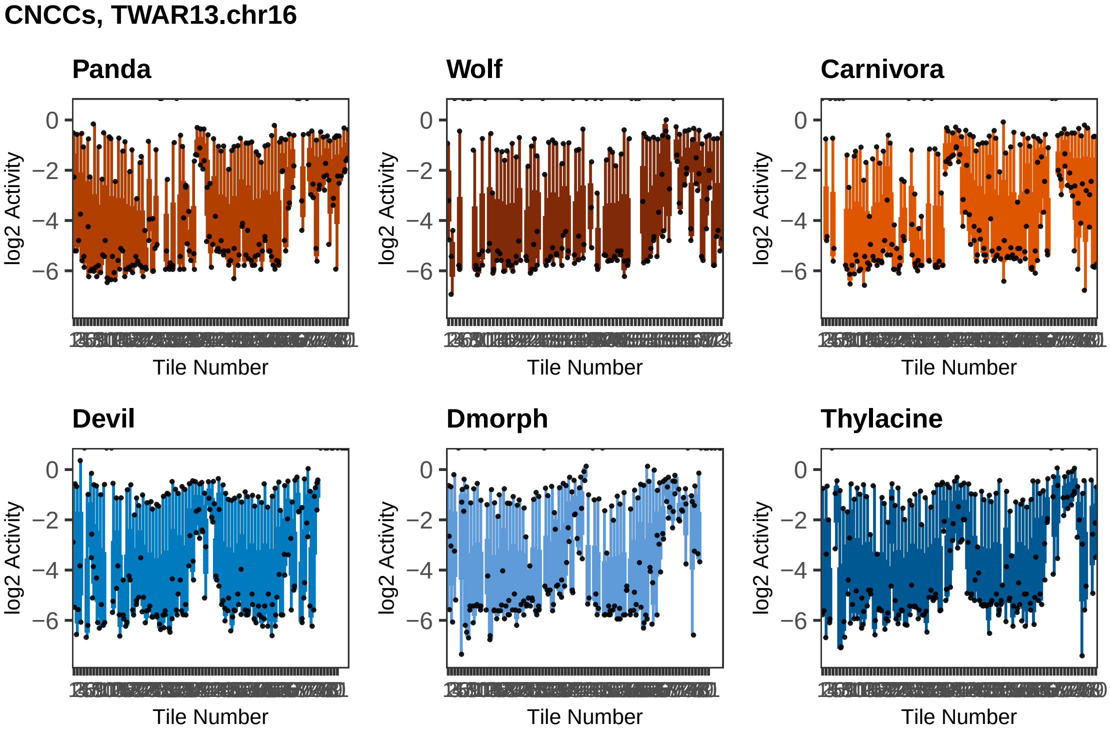
CNCCs, TWAR13.chr16
Panda
Wolf
Carnivora
Devil
Dmorph
Thylacine
log2 Activity
0
−2
−4
−6
Tile Number

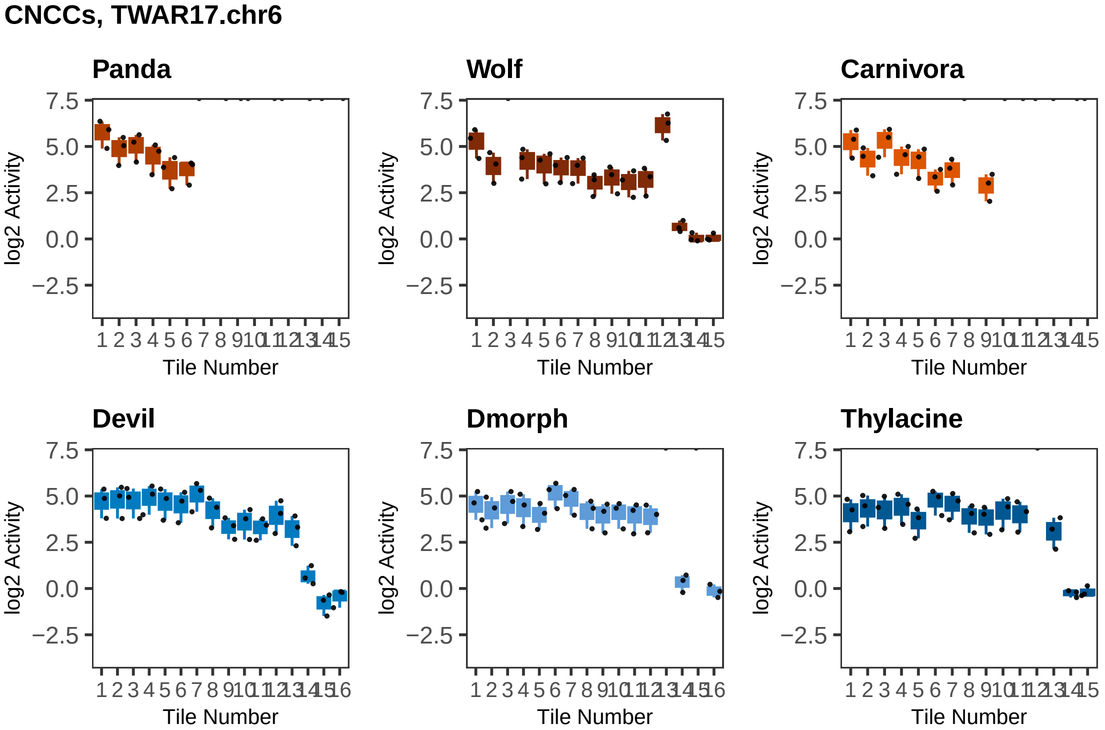

CNCCs, TWAR17.chr6
Panda
Wolf
Carnivora
Devil
Dmorph
Thylacine
log2 Activity
Tile Number
7.5
5.0
2.5
0.0
−2.5

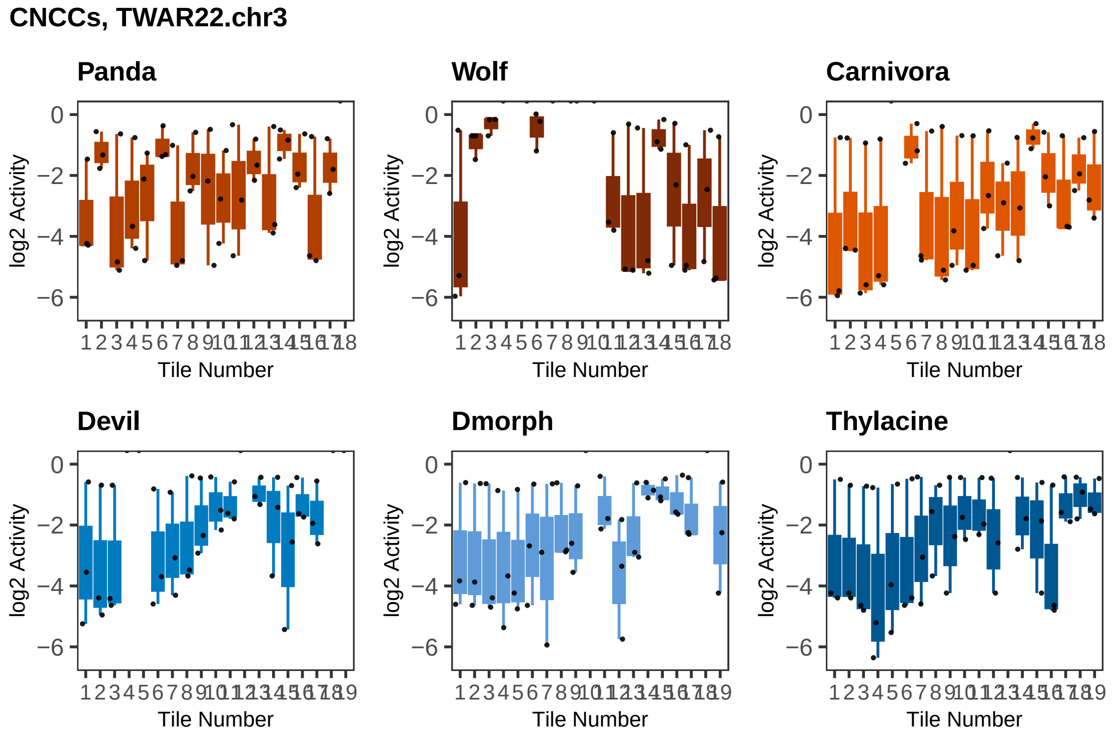
CNCCs, TWAR22.chr3
Panda
Wolf
Carnivora
Devil
Dmorph
Thylacine
log2 Activity
0
−2
−4
−6
Tile Number
1 2 3 4 5 6 7 8 9 10 11 12 13 14 15 16 17 18
1 2 3 4 5 6 7 8 9 10 11 12 13 14 15 16 17 18 19

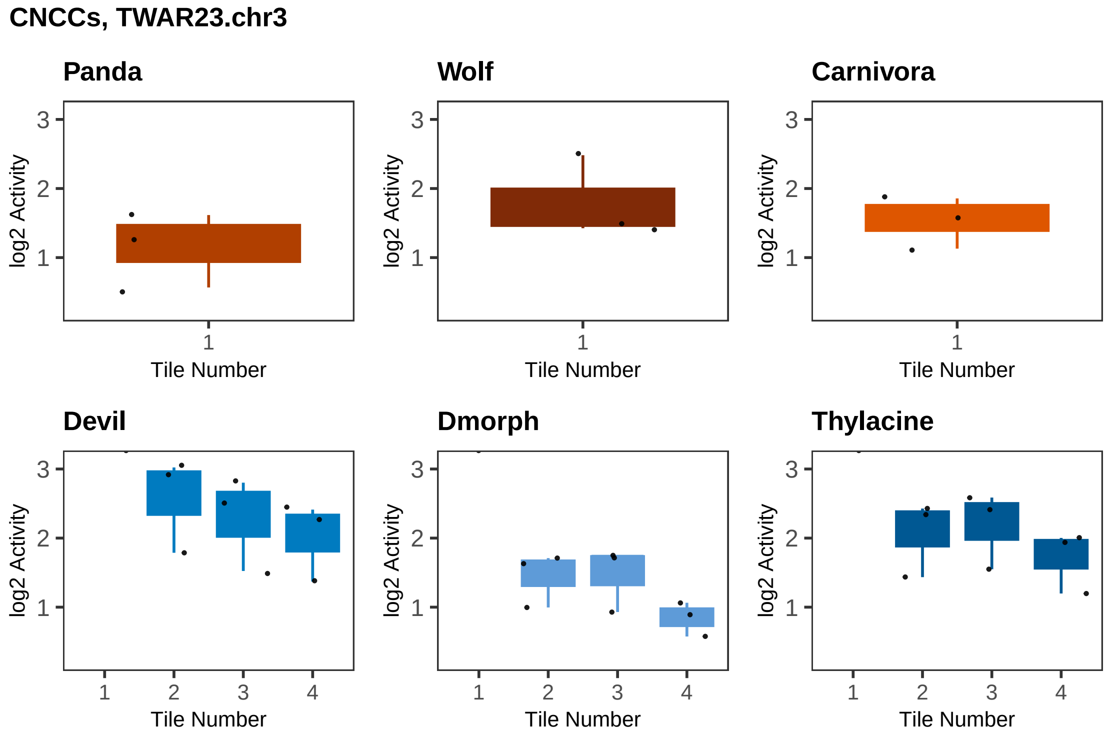
CNCCs, TWAR23.chr3
Panda
Wolf
Carnivora
Devil
Dmorph
Thylacine
log2 Activity
Tile Number
1
2
3
4

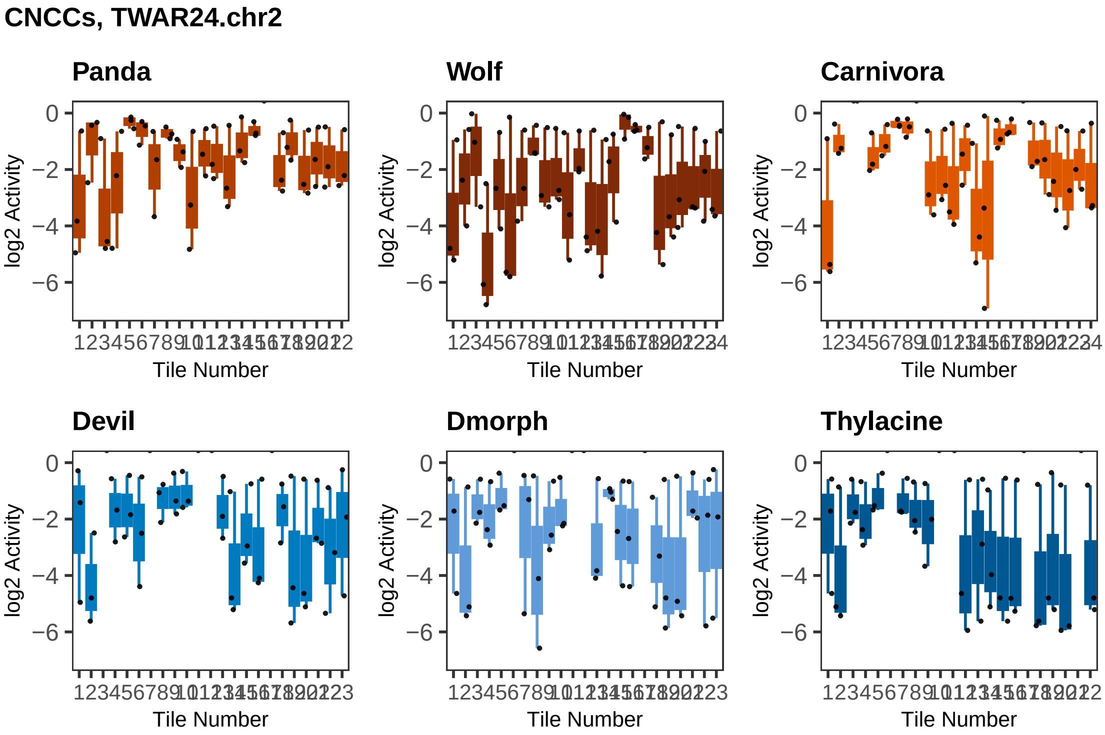
CNCCs, TWAR24.chr2
Panda
Wolf
Carnivora
Devil
Dmorph
Thylacine
log2 Activity
Tile Number
0
−2
−4
−6

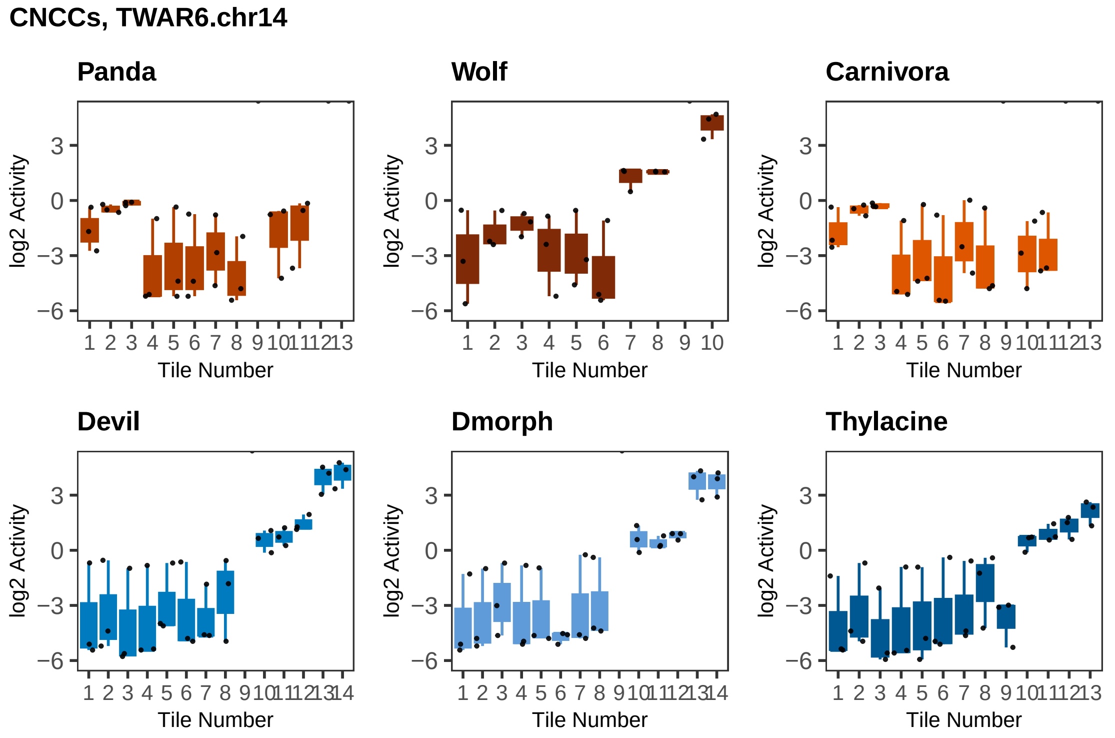

CNCCs, TWAR6.chr14
Panda
Wolf
Carnivora
Devil
Dmorph
Thylacine
log2 Activity
Tile Number
